# Computational prediction of SARS-CoV-2 encoded miRNAs and their putative host targets

**DOI:** 10.1101/2020.11.02.365049

**Authors:** Sonia Verma, Abhisek Dwivedy, Neeraj Kumar, Bichitra K. Biswal

## Abstract

Over the past two decades, there has been a continued research on the role of small non-coding RNAs including microRNAs (miRNAs) in various diseases. Studies have shown that viruses modulate the host cellular machinery and hijack its metabolic and immune signalling pathways by miRNA mediated gene silencing. Given the immensity of coronavirus disease 19 (COVID-19) pandemic and the strong association of viral encoded miRNAs with their pathogenesis, it is important to study Severe Acute Respiratory Syndrome Coronavirus-2 (SARS-CoV-2) miRNAs. To address this unexplored area, we identified 8 putative novel miRNAs from SARS-CoV-2 genome and explored their possible human gene targets. A significant proportion of these targets populated key immune and metabolic pathways such as MAPK signalling pathway, maturity-onset diabetes, Insulin signalling pathway, endocytosis, RNA transport, TGF-β signalling pathway, to name a few. The data from this work is backed up by recently reported high-throughput transcriptomics datasets obtains from SARS-CoV-2 infected samples. Analysis of these datasets reveal that a significant proportion of the target human genes were down-regulated upon SARS-CoV-2 infection. The current study brings to light probable host metabolic and immune pathways susceptible to viral miRNA mediated silencing in a SARS-CoV-2 infection, and discusses its effects on the host pathophysiology.

## Background

SARS-CoV-2 is the genesis of 2019–20 coronavirus pandemic, COVID-19. SARS-CoV-2 belongs to the genera Betacoronavirus of subfamily *Orthocoronavirinae* of the *Coronaviridae* family and order *Nidovirales* (1). SARS-CoV-2 is an enveloped virus with single-stranded RNA genome of positive polarity. The 29903 nucleotides long RNA genome consists of 10 Open Reading Frames (ORFs) and encodes 26 proteins (NCBI Reference Sequence: NC_045512.2, SARS-CoV-2 Isolate Wuhan Hu-1) (2). Two additional betacoronaviruses, Severe Acute Respiratory Syndrome Coronavirus (SARS-CoV) and Middle East Respiratory Syndrome Coronavirus (MERS-CoV), have evolved and caused viral epidemics in the past twenty years. SARS-CoV and MERS-CoV, respectively have infected over 8000 and 857 individuals globally and are associated with high mortality rates (3). The SARS-CoV-2 has infected over 40 million people worldwide and caused over 1.1 million deaths as of October 2020, resulting in a high fatality rate (4). The common symptoms of COVID-19 are fever, sore throat, dry cough, breathlessness, malaise, and fatigue. In rare cases, patients develop acute respiratory distress syndrome and sepsis followed by respiratory and heart failure (5). One of the challenges to control this virus is unavailability of effective therapeutics. The problem is further compounded by the lack of our understanding regarding molecular mechanisms involved in the pathogenesis of SARS-CoV-2 including the host-pathogen interactions.

MiRNAs are 18-22 nucleotides long noncoding RNAs that are found in animals, plants, and some viruses. They regulate genetic network pathways by targeting messenger RNAs (mRNAs) to translational repression or degradation. The biogenesis of miRNAs begins by RNA polymerase II mediated transcription of genome into long primary-miRNAs (pri-miRNAs). The nuclear Ribonuclease type III enzyme, Drosha cleaves pri-miRNA into small hairpin precursor-miRNAs (pre-miRNAs) which are then exported to the cytoplasm by nuclear membrane protein, Exportin5. In the cytoplasm, another Ribonuclease III enzyme-Dicer processes the pre-miRNA to generate a miRNA duplex intermediate (miRNA: miRNA*). One strand of this intermediate, mature miRNA binds to protein Argonaute-1 generating the miRNA inducing silencing complex (RISC) which guides the binding of mature miRNA to target mRNAs (6). Besides eukaryotes, viruses also encode miRNAs to target host genes involved in important biological processes like cell growth, cell differentiation, and cellular defense mechanisms as well as to regulate viral gene expressions to modulate the viral replication cycle (7). Therefore, viral miRNAs aid virus particles in their continuous proliferation and evasion of the host immune system (8,9).

Various studies have demonstrated a strong association between viral miRNAs and infection (8,9). However, there is no evidence to suggest the presence of miRNA sequences in SARS-CoV-2 genome. To address this problem, we applied computational approaches to analyze the SARS-CoV-2 genome sequence for possible miRNAs. This was of high interest and therefore the secondary goal of the study was to investigate the potential targets of these miRNAs along with their associated Gene Ontologies (GO) and signalling pathways. Our data provides a list of potential viral miRNAs from SARS-CoV-2 and predicts their roles in COVID-19 pathophysiology, thus facilitating further works on the potential biological roles of these miRNAs.

## Results

### The putative miRNAs in SARS-CoV-2 genome

As pre-miRNAs form hair-pin loop structures during their biogenesis, we first searched for such arrangements in the SARS-CoV-2 genomic sequence using the VMir software (10). The analysis demonstrated that the direct and reverse genomic strands consist of twenty-four and twenty-two hair-pin loop sequences, respectively. The length of these sequences ranged between 70 and 148 nucleotides, and they had VMir score ≥115, hairpin size ≥ 70, and window count ≥ 35 (**Figure 1A, Supplementary Material - Sheet 1**).

**Figure 1.**
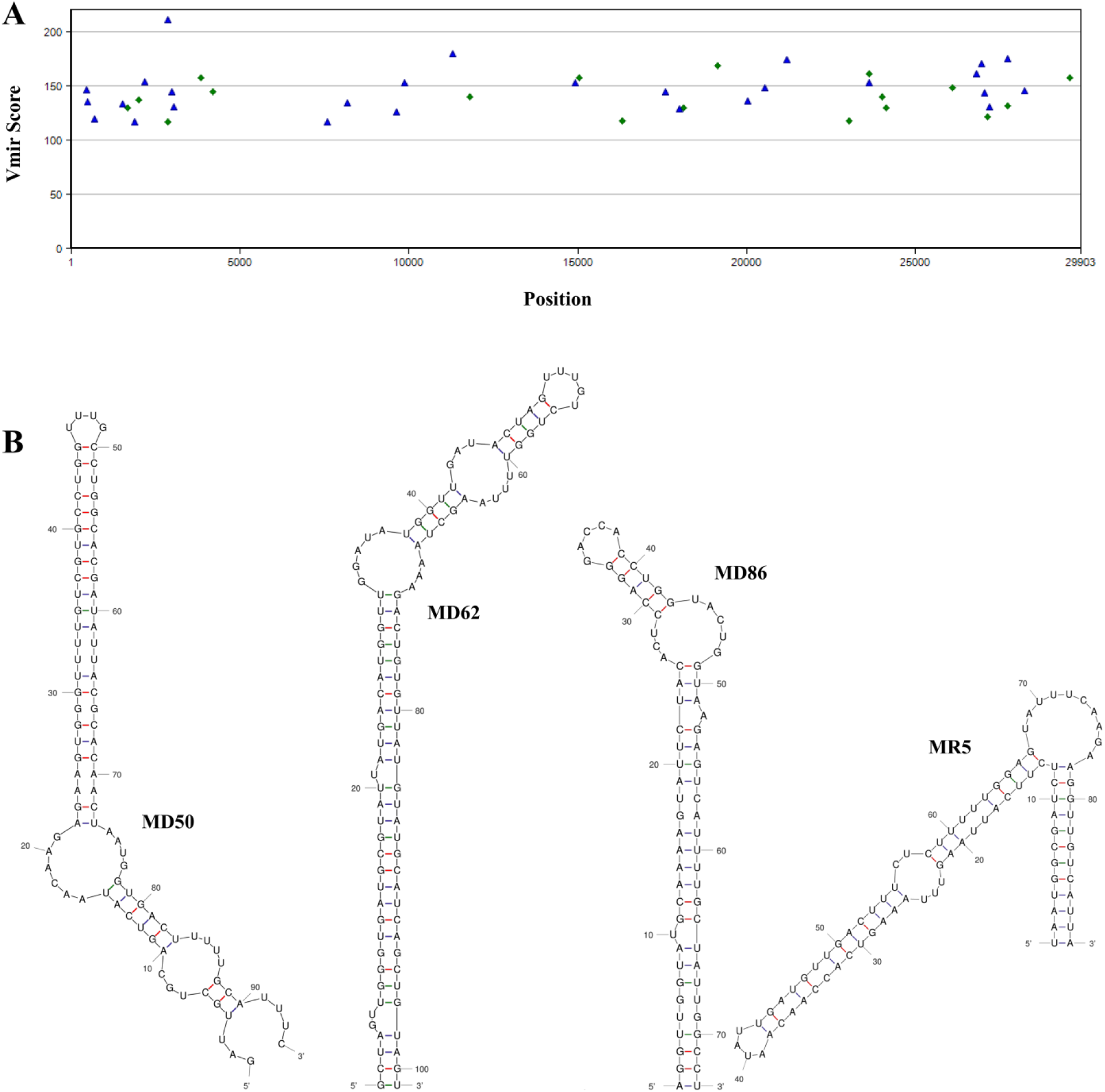
Potential stem-loop structures of pre-miRNAs of SARS-CoV-2. **(A)** VMir analysis of the SARS-CoV-2 genomic RNA; all hairpins with VMir score ≥115, hairpin size ≥ 70, and window count ≥ 35 are shown. Hairpins are plotted according to genomic position (X-axis) and VMir score (Y-axis). (Hairpins in direct orientation represented as blue triangles; and hairpins in reverse orientation represented as green diamonds). **(B)** Secondary structures of the predicated stem-loops of most likely pre-miRNAs candidates generated using MFold.

Next, using MFold web server the Minimal Folding Free Energy (MFE) of each sequence was calculated because MFE of the pre-miRNA folding is inversely proportional to the thermodynamic stability of the molecule (11). The secondary structure of these sequences was also predicted using MFold. In total, forty-two potential pre-miRNAs (direct strand-23; reverse strand-18), with MFE ≤ −18 kcal/mol, were recognized to be thermodynamically stable **(Supplementary Material − Sheet 2)**.

The Minimal Folding Free Energy Index (MFEI) distinguishes miRNAs from other coding and non-coding RNAs as pre-miRNAs display higher MFEI value than mRNAs (0.66), tRNAs (0.64) and rRNAs (0.59) (12). Therefore, the MFEI for the forty-one potential pre-miRNAs was calculated, and the values ranged from −0.60 to −0.93 **(Supplementary Material-Sheet 3)**. The pre-miRNAs MD50, MD62, and MD86 on the direct strand and MR5 on the reverse strand, with MFEI ≤ −0.85 kcal/mol, were chosen as the best candidates for the prediction of mature miRNAs via MatureBayes tool **(Figure 1B)** (13). The MatureBayes prediction revealed 22-nucleotide long 8 mature miRNAs from the pre-miRNA sequences on their 5’ and 3’ stem locations **(Table 1)**. As both the strands can serve as mature miRNA depending on the assembly of RISC complex, mature miRNAs on both the arms of stem-loop were used for further analysis.

**Table 1.**
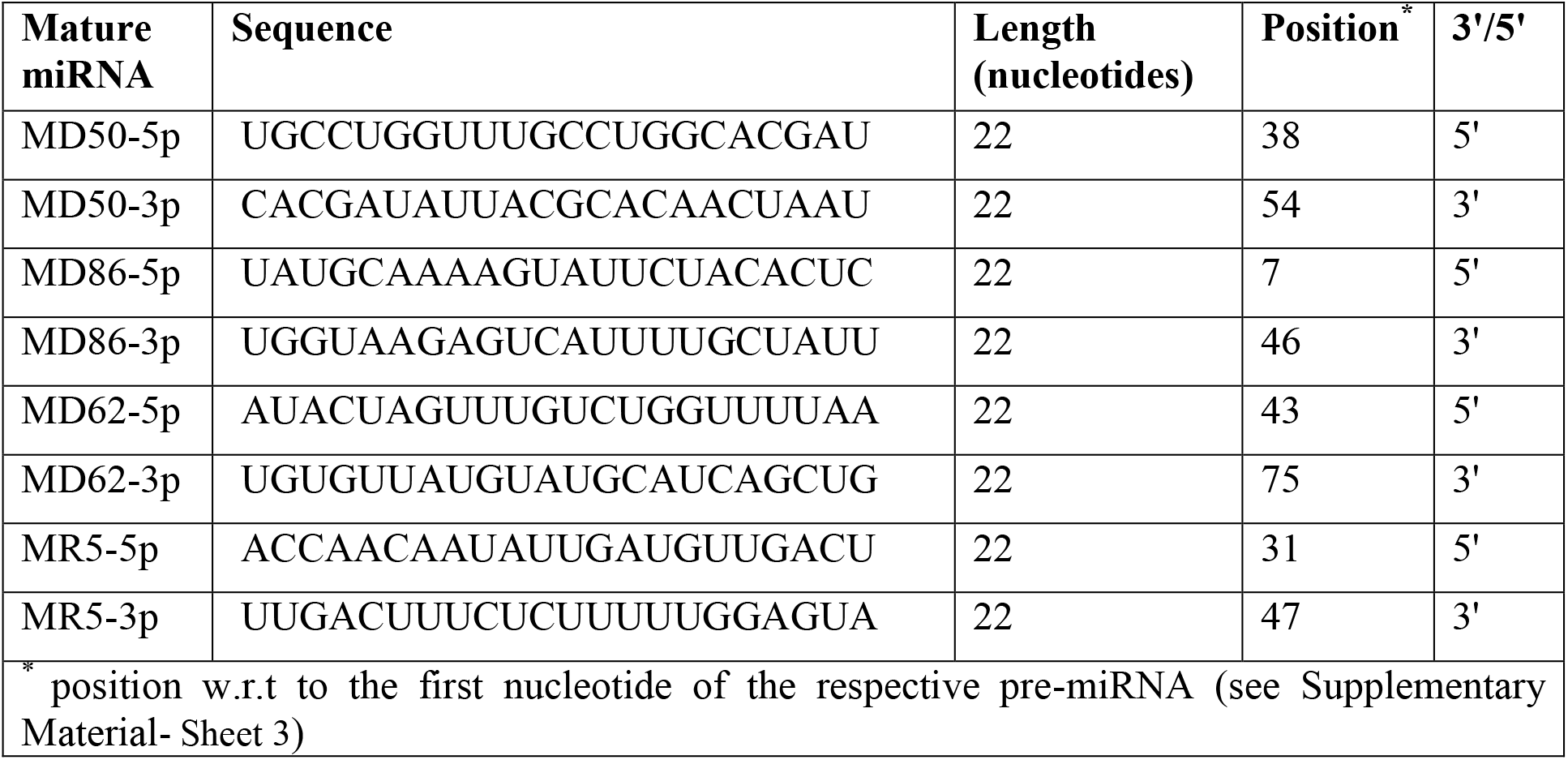
List of putative misRNAs predicted by Mature-Bayes.

This work is the first to report that SARS-CoV-2 genome may encode miRNAs and therefore careful validation of these findings was carried out against previously published data. A systematic and comprehensive literature search for RNA sequencing data of SARS-CoV-2 infected samples extracted out a study from the Blanco-Melo *et al*. group. The study explored the transcriptional response of SARS-CoV-2 infected epithelial cell lines, A549, and A549-ACE-2 cells (14). They reported that over 50% of the sequencing reads from each infected sample aligned with viral sequences while the mock-treated cells showed no viral reads. An alignment search using the mature miRNA sequences was executed against the sequencing reads provided for these samples in the Gene Expression Omnibus (GEO) Dataset: GSE147507. The search identified MR5-5p and MR5-3p together in 15 and 38758 reads from the infected A549 and A549-ACE-2 cells, respectively. The lengths of these sequences varied between 48 to 151 nucleotides proposing the likelihood of the intermediate products of miRNA maturation process. This further implies for the physiological presence of pre-miRNA like molecules in SARS-CoV-2 infected tissues.

### Predicting the target mRNAs of the SARS-CoV-2 encoded miRNAs in the human genome

The miRNAs exert their effects by targeting one or more mRNAs. Therefore, predicting the targets of putative miRNAs of SARS-CoV-2 would help understand the pathogenesis of the disease and shed light on the possible host-pathogen interactions. *In silico* prediction of targets by miRDB (15) for all 8 mature miRNAs retrieved 1,121 unique targets in the human genome with score ≥ 80 **(Supplementary Material-Sheet 4)**. Interestingly, some of these targets have already been associated with pulmonary diseases. For instance, the gene coding for IL-1α was one of the predicted targets for SARS-CoV-2 encoded miRNAs. Interleukin-1α/β (IL-1α/β) are key pro-inflammatory cytokines that mediate acute pulmonary inflammation and enhances survival during influenza virus infection by recruiting CD4^+^ T cells and aiding to the IgM antibody response (16). Likewise, another predicted target, Bone morphogenetic protein receptor 2 maintains pulmonary arterial hypertension, a form of high blood pressure that affects the arteries of the lungs and the heart (17). Of note, high blood pressure is a significant risk factor in COVID-19 (18). Defects in CHRNA3 and CHRNA5, members of nicotinic acetylcholine receptor family of proteins, are linked with various lung cancers and smoking-related lung disorders (19). Studies showed increased susceptibility of smokers to contract COVID-19 (20) and the genes coding for CHRNA3 and CHRNA5 were also reported amongst the SARS-CoV-2 encoded miRNAs targets in our study.

Blanco-Melo *et al*. reported 77 and 96 genes down-regulated in infected A549 and A549-ACE-2 cell lines, respectively. Of these, 6 and 4 genes respectively were found to be the targets of SARS-Cov-2 miRNAs in current study **(Figure 2Ai, 2Aii, and 2B, Supplementary Material-Sheet 5)**. The group also reported 2408 genes down-regulated in the post mortem lung samples from COVID-19 patients; of which 126 were predicted as the targets of SARS-Cov-2 miRNAs in our study **(Figure 2Aiii and 2B, Supplementary Material-Sheet 5)** (14). Our literature mining retrieved two additional studies on transcriptional response of COVID-19 patients. Xiong *et al*. performed a transcriptomic analysis of both broncho-alveolar lavage fluid (BALF) and peripheral blood of COVID-19 patients and compared with samples from healthy donors (21). Of the 325 and 316 genes reported to be down-regulated in BALF and peripheral blood samples, we found 15 and 7 genes respectively as putative targets of the SARS-Cov-2 miRNAs **(Figure 2Aiv, 2Av, and 2B, Supplementary Material-Sheet 5)**. Zhou *et al*. reported a metatranscriptome sequencing of BALF from COVID-19 patients and found 739 genes to be down-regulated. 56 genes reported as targets of SARS-CoV-2 miRNAs were also found to be down-regulated in their analysis **(Figure 2Avi and 2B, Supplementary Material-Sheet 5)** (22). Examples of such genes include ABCA1 and CDC42. ABCA1 regulates cholesterol efflux and in association with CDC42 regulates the formation of lipid rafts (23)(24). Lipid rafts are key features of many cardiovascular and inflammatory diseases such as diabetes, which have also been associated with COVID-19 (25). Taken together, many of the targets that we predicted for the SARS-Cov-2 miRNAs have also been reported to be down-regulated during COVID-19 in other studies, signifying a possible viral miRNA mediated repression of human genes.

**Figure 2.**
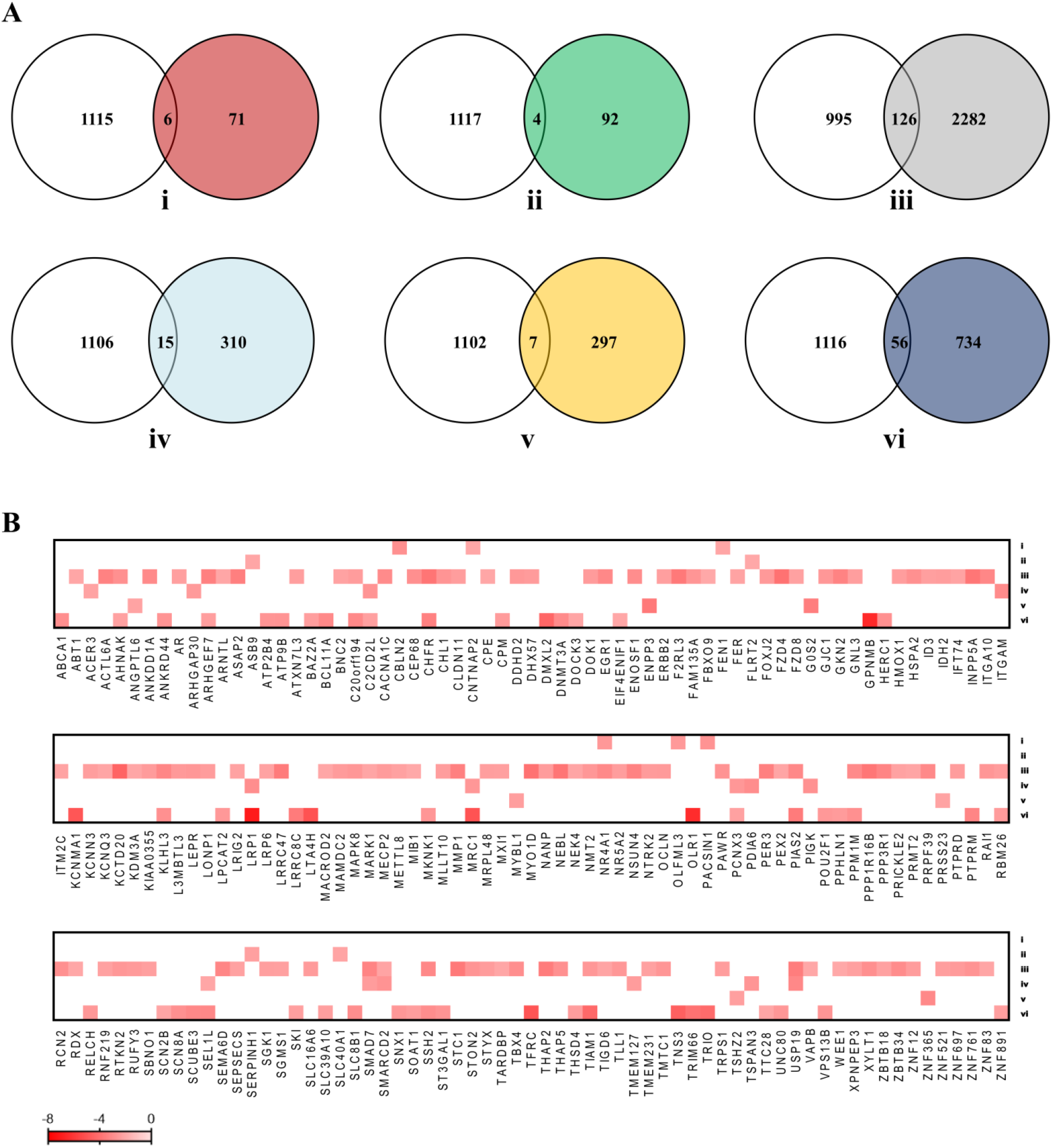
Target mRNAs are reported to be down-regulated in SARS-CoV-2 infected samples. **(A)** Venn diagrams showing the number of overlapping genes between the SARS-CoV-2 miRNAs target genes predicted in the current study (white circles) and the down-regulated genes in the SARS-CoV-2 infected samples of three independent transcriptomic studies (colored circles)-A549 cells (Blanco-Melo *et al*.) **(i)**, A549-ACE-2 cells (Blanco-Melo *et al*.) **(ii)**, Post-mortem lung biopsies of COVID-19 patients (Blanco-Melo *et al*.) **(iii)**, Bronchio-Alveolar Lavage Fluid of COVID-19 patients (Xiong *et al*.) **(iv)**, peripheral blood of COVID-19 patients (Xiong *et al*.) **(v)** and Bronchio-Alveolar Lavage Fluid of COVID-19 patients (Zhuo Zhou *et al*.) **(vi)**. **(B)** Heat map showing the expression profiles of overlapping genes in the studies **(i-vi)** mentioned above (white indicates the absence of the expression levels in the respective study).

### The potential physiological effects of the SARS-CoV-2 miRNAs on a host cell

The GO analysis of the 1121 potential targets of the SARS-CoV-2 miRNAs was executed using the Cytoscape plugin, ClueGo (26,27) and the results indicated that the major Cellular Component terms populated were the nucleus, cytoplasm, cell junction, nucleoplasm, and endosome **(Figure 3A, Supplementary Material-Sheet 6)**. The Molecular Function enrichment depicted functions such as protein binding, mRNA 3’-untranslated regions (UTR) binding, zinc ion binding, transcription factor activity, and sequence-specific DNA binding **(Figure 3B, Supplementary Material-Sheet 6)**. The GO-Biological Processes analysis suggested that the target gene products primarily populate the negative regulation of transcription from RNA polymerase II promoter, positive regulation of transcription (DNA-templated), positive regulation of transcription from RNA polymerase II promoter, transcription (DNA-templated), and mitotic cell cycle **(Figure 3C, Supplementary Material-Sheet 6)**. The Kyoto Encyclopedia of Genes and Genomes (KEGG) pathway analysis showed that the target genes were enriched in MAPK signalling pathway, ErbB signalling pathway, maturity-onset diabetes of the young, Insulin signalling pathway, Ras signalling pathway, endocytosis, RNA transport, TGF-β signalling pathway, to name a few **(Figure 3D, Figure 4A-D, Supplementary Material-Sheet 6)**. Interestingly, studies have demonstrated a strong and consistent link between syndromes associated with disruption of few of these pathways and COVID-19. For instance, COVID-19 patients with no prior history of type-2 diabetes have been reported to show hyperglycaemia, lowered insulin levels, and pre-diabetic symptoms (28). The genes coding for PAX4 and PAX6, key proteins involved in the development of pancreatic islets which are often found mutated and/or dysregulated in type 2 diabetes, were found to be potential targets for SARS-CoV-2 miRNAs in our study (29,30).

**Figure 3.**
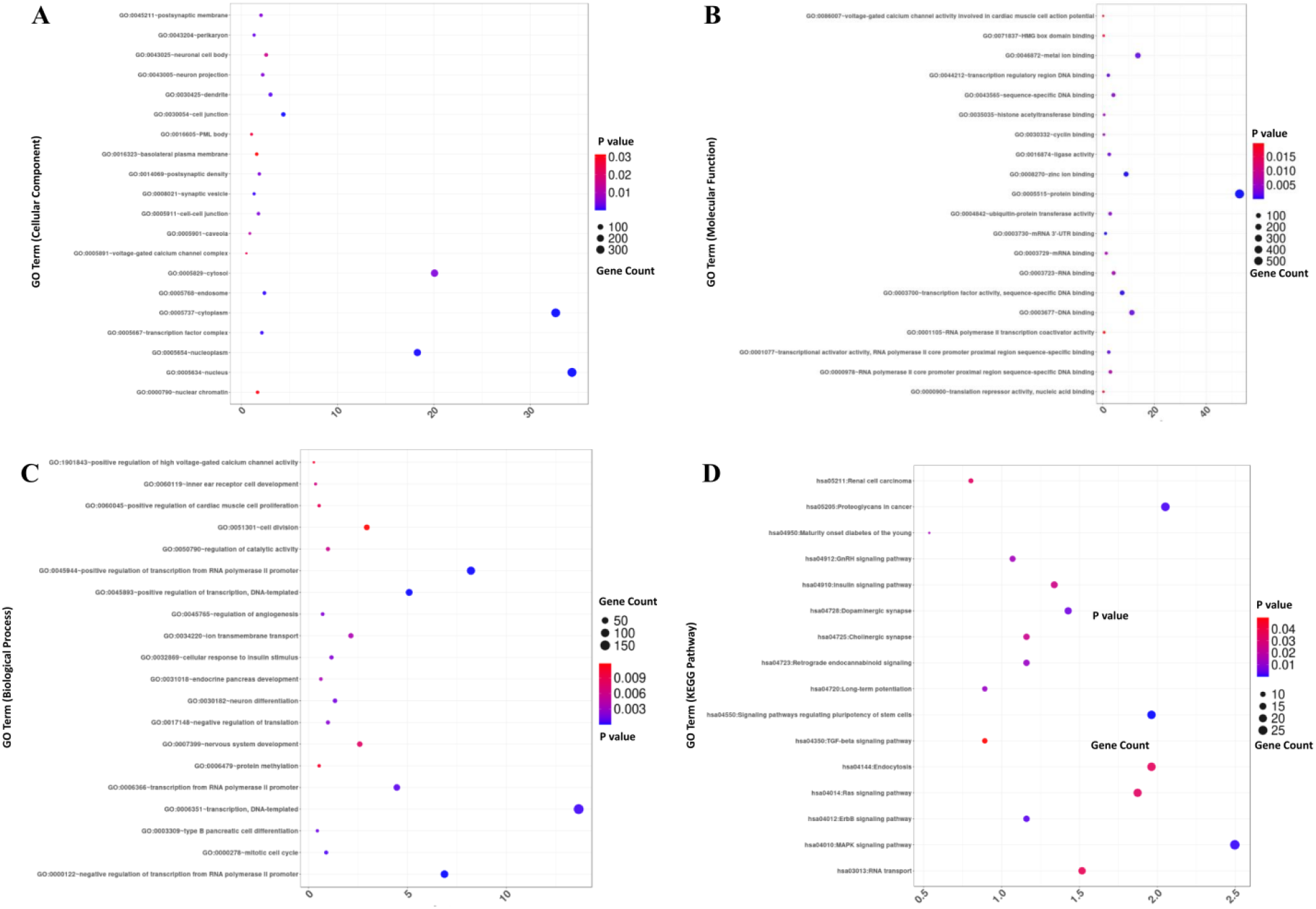
Gene Ontology and Pathway Analysis of host target genes for SARS-CoV-2 miRNAs. Top 20 most significant cellular components **(A)**, molecular functions **(B)**, biological processes **(C),** and KEGG pathway enrichment **(D)** analysis of 1121 target genes are shown.

**Figure 4.**
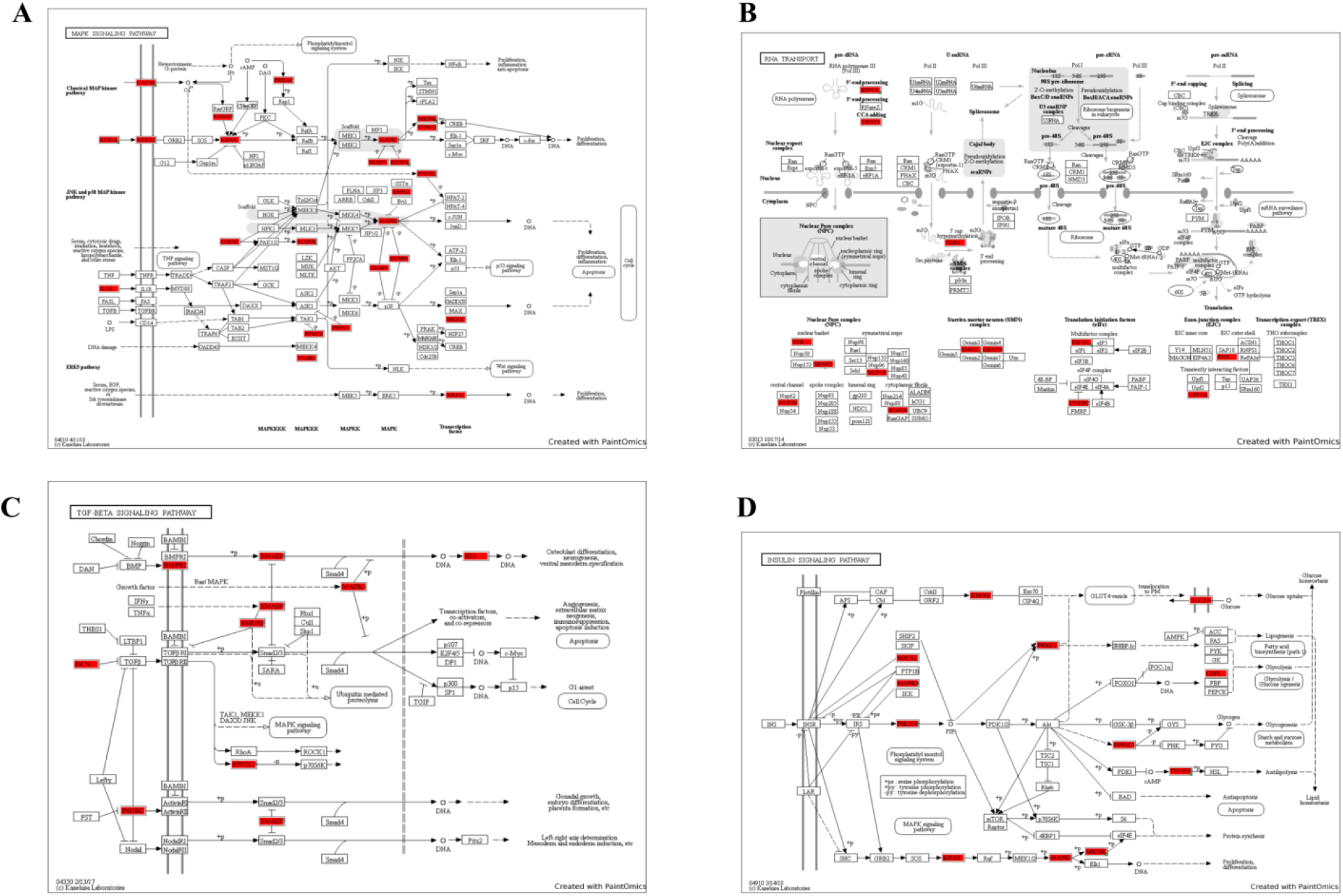
KEGG pathways populated with the host target genes for SARS-CoV-2 miRNAs. Painted MAPK signalling pathways **(A)**, RNA transport **(B)**, TGF-β signalling pathway **(C)**, and Insulin signalling pathways **(D)** with genes in red boxes identified as targets of predicted SARS-CoV-2 miRNAs.

In a cell, different proteins interact with each other to form a complex protein-protein interaction (PPI) network that comprises of over thousands of interactions. Knowing the interactions between the SARS-CoV-2 miRNA targets would provide an understanding of how the virus controls various molecular and biological processes of its host thus aiding in its survival and replication, while simultaneously evading the host responses. A PPI network of the 1121 targets was constructed using the ReactomeFI plugin in Cytoscape **(Figure 5A)** (27). The network included 308 proteins/nodes and 593 interactions/edges. To identify the significant interactions within the PPI network, the Molecular COmplex DEtection (MCODE) plugin (31) in Cytoscape was utilized, which predicted 16 significant interaction modules. The graphical representation of the top modules with an interaction score greater than 4 is shown in **Figure 5B**. In total these modules populated 85 genes, the majority of them are involved in MAPK signalling pathway, RNA transport, and mRNA surveillance. Of note, the RNA transport pathway was connected to the MAPK signalling pathway and the mRNA surveillance pathway through PPP2R3A and UPF3A, respectively **(Figure 6, Table 2)**. PPP2R3A is a key serine/threonine phosphatase that negatively regulates cell growth, while UPF3A detects truncated mRNA and initiates nonsense-mediated decay (32,33). Taken together, the SARS-CoV-2 hijacks the host metabolism and cellular processes for evading host-mediated RNA degradation and actively maintaining the cell in a growth phase, thus successfully its viral cycle (34,35).

**Figure 5.**
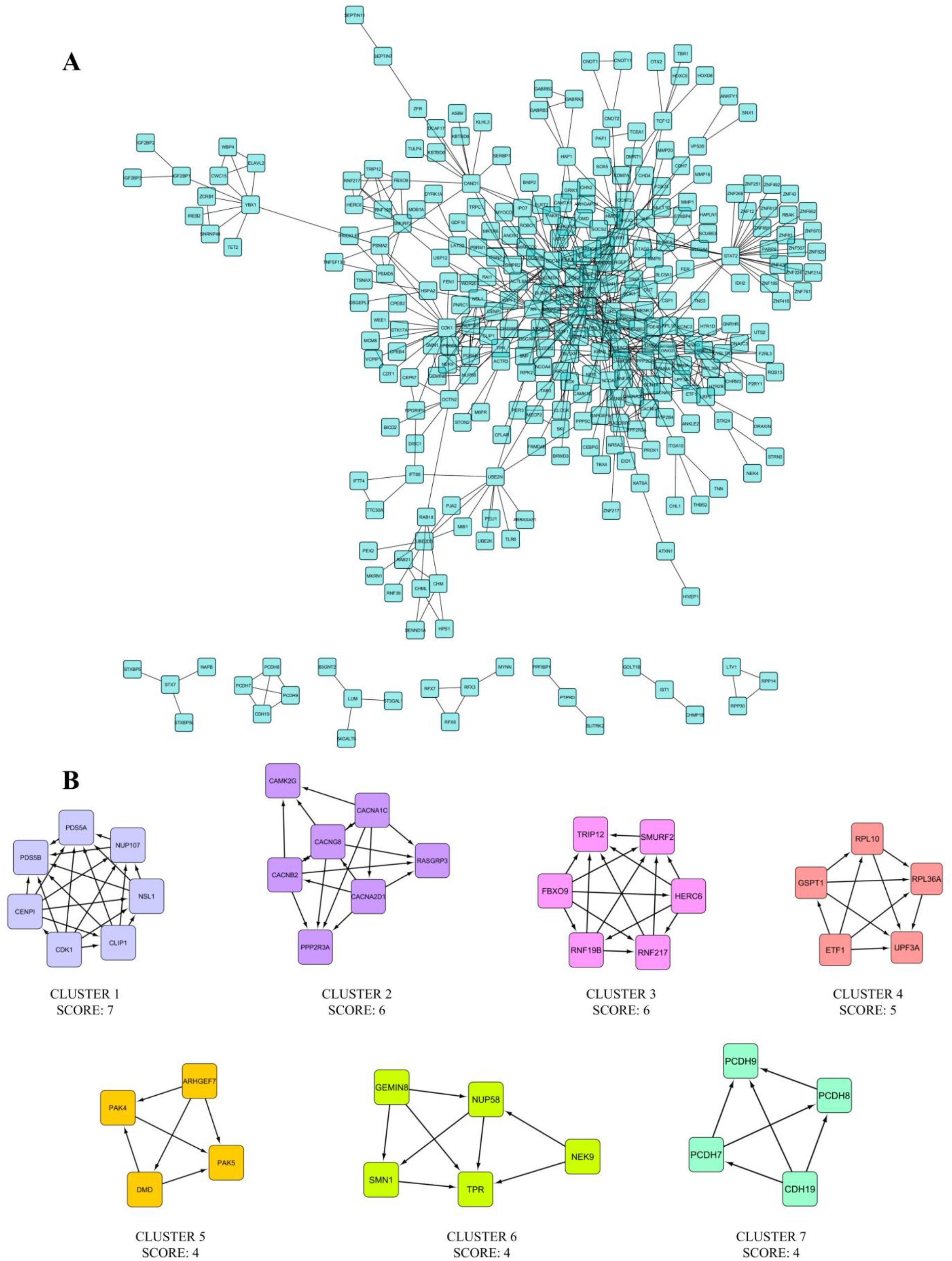
Construction and analysis of target genes from the interaction network. **(A)** PPI network of the target genes of SARS-CoV-2 miRNAs constructed using ReactomeFI plugin in Cytoscape. The node represents the genes, and the edge represents the interaction among the products of the genes. **(B)** Modules in the PPI network with most significant interactions belong majorly to the MAPK signalling pathway generated by MCODE analysis (see Figure 6).

**Figure 6.**
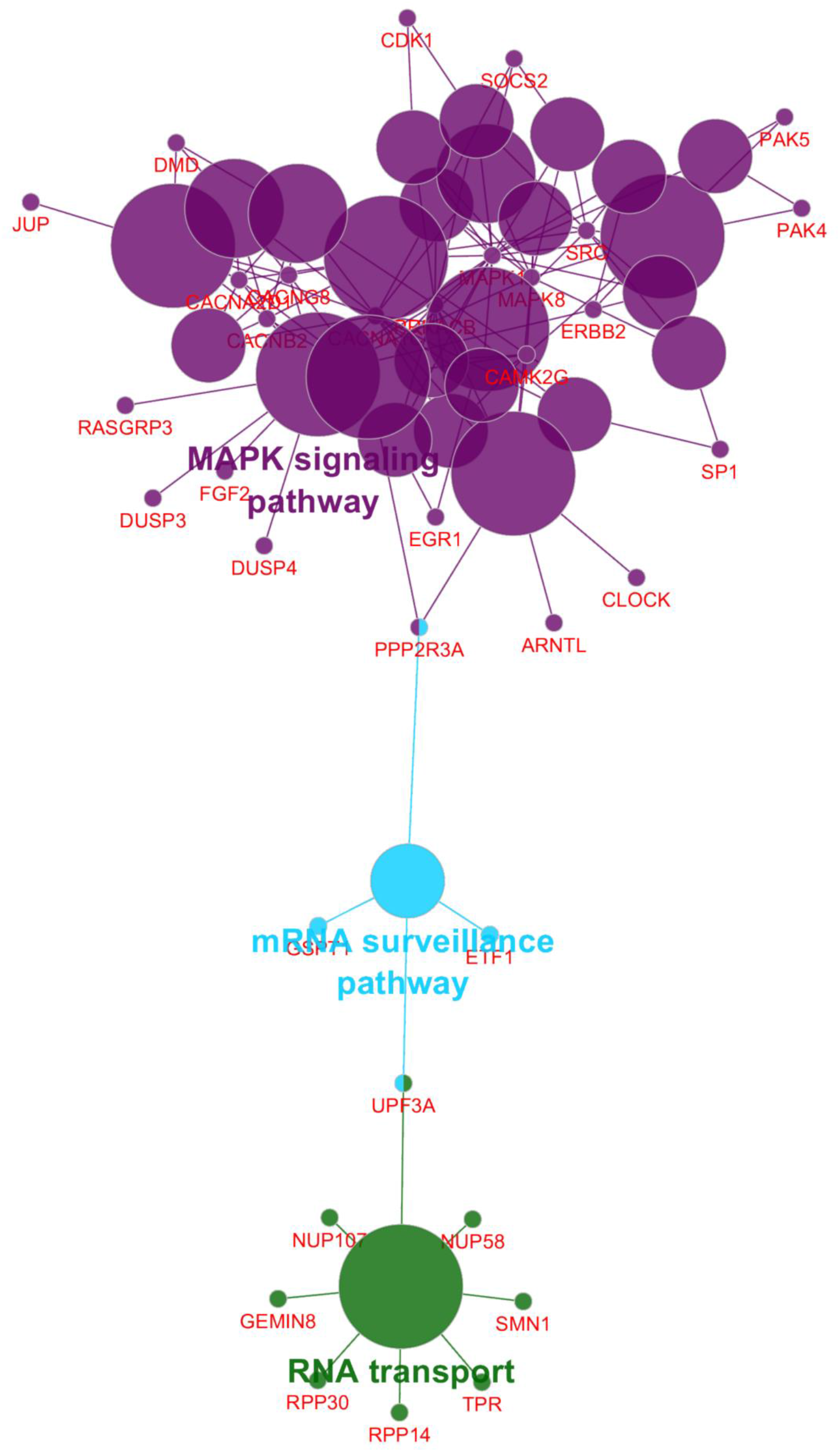
Analysis of modules reveal significant signalling pathways. Visualization of KEGG pathway enrichment profiles from the clustered genes of significant modules using ClueGO/CluePedia plugin of Cytoscape software.

**Table 2.** List of KEGG pathways enriched for the target mRNAs in the modules.

| KEGG ID | KEGG Term | Count | p-Value | Genes |
| --- | --- | --- | --- | --- |
| hsa03013 | RNA transport | 8 | 1.5E-05 | GEMIN8, NUP107, NUP58, RPP14, RPP30, SMN1, TPR, UPF3A |
| hsa03015 | mRNA surveillance pathway | 4 | 2.6E-03 | ETF1, GSPT1, PPP2R3A, UPF3A |
| hsa04010 | MAPK signalling pathway | 12 | 2.1E-07 | CACNA1C, CACNA2D1, CACNB2, CACNG8, DUSP3, DUSP4, ERBB2, FGF2, MAPK1, MAPK8, PRKACB, RASGRP3 |

## Discussion and Conclusions

The SARS-CoV-2 attaches to and enters the host cell by binding of its spike (S) protein to the host cell membrane protein, angiotensin-converting enzyme 2 receptors (36). After a SARS-CoV-2 virion attaches to its target cell, the cell’s trans-membrane Serine Protease 2 (TMPRSS2) cleaves the S protein to expose a fusion peptide (37). The virion then releases its RNA into the cell, taking over the cellular translation machinery to generate multiple copies of the virus that subsequently egress to infect more cells. The RNA genome of the virus harbours significant UTRs both at the 5’ and 3’ ends of its open-reading frames that give rise to a range of RNA secondary structures with unknown functions. A recent study has demonstrated the conservation of these structures across many betacoronaviruses (38). RNA secondary structures are primarily involved in enzymatic reactions and are also processed to generate miRNA that regulates host gene silencing. Studies have shown that viruses exploit the cellular machinery to encode miRNA and like eukaryotic miRNAs, the viral miRNAs are generated using the Drosha and/or Dicer proteins of the host (39).

Primarily, DNA viruses such as Herpesviruses, Papillomavirus, and Polyomavirus encode miRNAs (40–42). Epstein–Barr virus, a Herpesviridae family member, expresses 4 mature miRNAs transcribed from its BART and BHRF1 regions as it switches from the latent to the lytic phase of its infection cycle (43). Kaposi’s sarcoma virus, another herpesvirus, expresses over 25 matured miRNAs transcribed from the region between the kaposin and ORF71 of its genome (Cai et al., 2005). These miRNAs are associated with the latency phase of infection and are reported to modulate various genes in the host immune system and cell cycle. Human Cytomegalovirus genome encodes for over 25 miRNAs, which have been shown to interfere with host immunity, thus increasing the virulence of the virus (45). HBV-miR-3, encoded by the Hepatitis B virus down-regulates the expression of HbcAg, thus inhibiting its replication cycle (46). Adenovirus encodes for the mature miRNA-mivaRNAII-138, which when expressed in HeLa cells modulated core cellular functions such as growth, RNA metabolism, DNA repair, and apoptosis (47). Another report shows that mivaRNAII-138 repressed the expression of CUL4, thus positively modulating the JNK signalling (48). However, recent studies have demonstrated RNA viruses, such as HIV-1, Influenza virus, and Ebola virus to encode viral miRNAs (49–51). Also, the external incorporation of pre-miRNA sequence into the viral RNA has been reported to generate functional miRNAs by cytoplasmic RNA viruses. For example, the incorporation of pri-miR-124 the Sindbis virus genomes resulted in the production of pre-miR-124 and miR-124 in a DGCR8-independent, exportin-5-independent, and Dicer-dependent manner (39). Also, the flavivirus tick-borne encephalitis virus was reported to express an Epstein–Barr virus miRNA through the integration of its miR-BART2 precursor in the 3’-UTR of its genome (52). These examples of miRNA production from cytoplasmic RNA viruses demonstrate that pre-miRNA stem-loops can be processed in the cytoplasm by Dicer-dependent and Drosha independent non-canonical cytoplasmic mechanisms.

Recent studies on the effect of SARS-CoV-2 on host metabolism and immunity have provided interesting correlations to our study. Our results suggest that SARS-CoV-2 miRNAs target a range of genes encoding for TGF-β signalling inhibitors such as DCN, INHBC, SMAD-7, SMURF, and MAPK1. Notably, the serum level of TGF-β1 is elevated during the early phase of SARS-CoVs infection (53). These result in rapid and massive edema and fibrosis of lungs that remodels and ultimately blocks the airways leading to functional failure of the lungs and death of the patients. The increased levels of TGF-β along with dysregulated signalling are possibly achieved by targeting these aforementioned negative regulators through viral miRNAs. Also, various evidence has shown that hyperglycaemia, regardless of the past medical history of diabetes has emerged as an important risk factor for severe illness and death from COVID-19 (28). Our results suggest that the SARS-CoV-2 miRNAs target the gene encoding SLC2A4, a glucose transporter responsible for the uptake of extracellular glucose into the cytoplasm. Also, the gene for RHOQ, a molecule involved in insulin-mediated glucose is a predicted target for SARS-CoV-2 miRNAs. Notably, the gene for KRAS, a positive regulator of another glucose transporter GLUT4 is amongst the SARS-CoV-2 miRNAs targets. The down-regulation of these molecules is associated with hyperglycaemia and severe diabetes.

T-cell receptor zeta or CD247 plays an important role in coupling antigen recognition to several intracellular signal-transduction pathways. Impaired expression of CD247 results in impaired immune response and is associated with immunosuppression in T-cells (54). Our predictions list the gene encoding CD247 as a target for SARS-CoV-2 miRNAs, and the transcriptomics analysis by Xiong *et al*. demonstrated a down-regulated expression of CD247 in PBMCs of SARS-CoV-2 infected patients (21). TLR8, a key player of the anti-viral innate immunity, is an endosome localized sensor of RNA degradation end products and actively recognizes viral and bacterial pathogens. Further, it activates IRF5 and induces the production of IFNβ, IL-12p70, and TNF (55). Of note, TLR8 is also reported to sense cytoplasmic microRNAs of both host and pathogen origins (56). Rhinovirus, the causative agent of the common cold is known to repress the expression of TLR8 mRNA (57).

This study presents a list of potential viral miRNAs from the SARS-CoV-2 genome and predicts a list of putative gene targets from the human host. Moreover, the study draws a link between genes involved in the SARS-CoV-2 infection-associated symptoms such as lung damage, diabetes, and cardiovascular diseases and their possible regulation via SARS-CoV-2 miRNAs. Identifying the presence of these viral miRNAs in COVID-19 patient samples will further enhance our understanding of the disease prognosis at a cellular and molecular level, thus aiding the on-going hunt for COVID-19 therapeutics.

## Methods

### Resources Table

| RESOURCE | SOURCE | IDENTIFIER |
| --- | --- | --- |
| **Deposited Data** |  |  |
| Genome sequence of SARS-CoV-2 (Isolate Wuhan Hu-1) | Wu F. *et al.*, 2020, *Nature* | NC_045512.2<br>(Nuccore, NCBI) |
| Transcriptional response to SARS-CoV-2 infection (A459 and A459-ACE2 cell lines infected with SARS-CoV2, post mortem lung samples of COVID-19 patients) | Blanco M. *et al.*, 2020; *Cell* | GSE147507<br>(GEO, NCBI) |
| The transcriptome of patients infected with SARS-CoV-2 (Peripheral Blood Mononuclear Cells and Broncho -Alveolar Lavage Fluid) | Xiong Y. *et al.*, 2020; *Emerg Microbes Infect* | CRA002390<br>(Genome Sequence Archive, BIG Data Center) |
| The BALF transcriptome data for Pneumonia patients (Broncho -Alveolar Lavage Fluid) | Zhou Z. *et al.*, 2020; *Cell Host Microbe* | PRJCA002273<br>(BioProject, BIG Data Center) |
| **Software and Algorithms** |  |  |
| VMir | Grundhoff A. *et al.*, 2006; *RNA* | Version 2.3 |
| MFold | http://unafold.rna.albany.edu/?q=MFold | Version 3.6 |
| Mature-Bayes | http://mirna.imbb.forth.gr/MatureBayes.html | N/A |
| miRDB | http://mirdb.org | N/A |
| Cytoscape | https://cytoscape.org | Version 3.8 |
| ClueGO | http://apps.cytoscape.org/apps/cluego | Version 2.5.7 |
| CluePedia | http://apps.cytoscape.org/apps/cluepedia | Version 1.5.7 |
| CytoKEGG | http://apps.cytoscape.org/apps/cytokegg | Version 1.0.1 |
| KEGGParser | http://apps.cytoscape.org/apps/cykeggparser | Version 1.2.9 |
| ReactomeFI | http://apps.cytoscape.org/apps/reactomefiplugin | Version 7.2.3 |
| MCODE | http://apps.cytoscape.org/apps/mcode | Version 1.6.1 |
| PaintOmics | http://www.paintomics.org/ | Version 3 |
| R | https://cran.r-project.org/ | Version 4.0.1 |
| R Studio | https://rstudio.com/ | Version 1.3.959 |

### Precursor miRNAs (pre-miRNAs) prediction

The SARS-CoV-2 genome sequence was downloaded from the NCBI with accession number NC_045512.2. VMir software (10), which consists of two programs VMir analyser and VMir viewer, was used with default settings (window size: 500 nucleotides, step size: 1 nucleotide, conformation: linear, orientation: both, hairpin size: any, minimum VMir score: 115, minimum hairpin size:70, minimum Window Count:35) to identify putative pre-miRNA hairpin structures in the SARS-CoV-2 genome.

### Secondary structure validation

The VMir-derived pre-miRNA sequences were submitted to the MFold (11) web-server with default parameters for secondary structure prediction and MFE (ΔG in kcal/mol) calculation. The MFEI was also calculated as described in the equation:

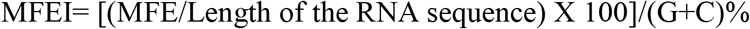

The potential pre-miRNA secondary structures were assessed using the following criteria:

i. MFE ≤ −18 kcal/mol
ii. (G+C)% between 30–70 %;
iii. MFEI ≤ −0.85 kcal/mol

### Mature miRNAs Prediction

The MatureBayes tool was used at default parameters to predict mature miRNAs (13). MatureBayes incorporates the Naïve Bayes classifier for identifying the putative mature miRNA molecules based on the sequence and structure of the miRNA precursors.

### Potential Target genes prediction

miRDB (15), a web server was used for predicting the target genes in the human genome for the mature miRNAs. The server uses the MirTarget algorithm, which is based on a 7-mer seeding approach to predict miRNAs targets in the human gene’s 3’ UTRs. The algorithm scans the 3’ UTR of human genes for hybridization with miRNAs sequence. Target genes with miRDB score ≥ 80 were selected as a predicted target with a score ≥ 80 is most likely to exist in a physiological scenario and not require any other supporting evidence.

### Gene Ontology and pathway enrichment analyses of the target mRNAs

Gene ontology is a bioinformatics tool used for investigating functional relationships between gene products and predicting three key aspects: biological process, cellular component, and molecular function. KEGG is a database used for analysing enriched pathways of the selected genes to further understand gene functions. To conduct GO and KEGG pathway analysis for the target mRNAs, the ClueGO (26), CluePedia (58), CytoKEGG, and KEGGParser (59) plugins on Cytoscape (27) were used. GO terms and KEGG pathways with p-value < 0.05 were considered statistically significant. Genes were mapped onto the KEGG pathways using the PaintOmics3 (60) using default parameters. Software packages R and R Studio were used to visualize the top GO terms and KEGG pathways.

### PPI Network construction

Using Cytoscape plugin, ReactomeFI (61), PPI network was constructed (*P* < 0.05, FWER Correction). Another Cytoscape plug-in, MCODE (31) was used to identify the interconnected clusters/ modules from the PPI network (degree cut-off = 2, max. depth = 100, k-core = 2, and node score cut-off = 0.2). ClueGO was used to identify the significant GO Terms enriched with the gene clusters retrieved in MCODE.

## Authors’ contributions

SV and AD conceptualized the study, performed the computational analysis and analysed the data. SV, AD, NK and BKB wrote the manuscript.

## Acknowledgements

The authors thank Mr. Krushna Chandra Murmu from Institute of Life Sciences, Bhubaneswar, for his suggestions and help in generating Figure 3 using the R package.

