## Supplementary Information for "Computational prediction of SARS-CoV-2 encoded miRNAs and their putative host targets"

**Sheet_1**

**List of potential stem-loop structures in SARS-CoV-2 genome as predicted by VMir.**

| Rank | Name | Start | Apex | Size | Score | Window Count (Absolute) |
|---|---|---|---|---|---|---|
| **Direct Orientation** | | | | | | |
| 1 | MD13 | 2801 | 2864 | 125 | 243.8 | 61 |
| 2 | MD62 | 11234 | 11286 | 101 | 211.4 | 49 |
| 4 | MD136 | 27666 | 27721 | 104 | 205.6 | 119 |
| 5 | MD108 | 21131 | 21184 | 110 | 204.7 | 210 |
| 9 | MD132 | 26743 | 26801 | 119 | 188.9 | 252 |
| 19 | MD56 | 9797 | 9858 | 128 | 179.1 | 59 |
| 26 | MD139 | 28196 | 28233 | 72 | 170.4 | 133 |
| 28 | MD16 | 2934 | 2974 | 76 | 169.9 | 71 |
| 43 | MD103 | 20002 | 20042 | 80 | 159.3 | 403 |
| 46 | MD6 | 1489 | 1531 | 86 | 156.7 | 171 |
| 51 | MD17 | 2981 | 3047 | 131 | 152.8 | 38 |
| 87 | MD4 | 651 | 692 | 75 | 140.3 | 46 |
| 95 | MD7 | 1810 | 1872 | 121 | 137.4 | 58 |

| | | | | | | |
|---|---|---|---|---|---|---|
| 116 | MD140 | 28217 | 28252 | 72 | 133.8 | 62 |
| 122 | MD55 | 9712 | 9758 | 96 | 132.5 | 49 |
| 135 | MD70 | 13171 | 13219 | 93 | 130.2 | 131 |
| 164 | MD95 | 18782 | 18820 | 79 | 124.7 | 184 |
| 173 | MD121 | 24086 | 24135 | 99 | 123.1 | 45 |
| 176 | MD96 | 19046 | 19086 | 75 | 123.1 | 179 |
| 196 | MD19 | 3197 | 3236 | 76 | 120.4 | 49 |
| 200 | MD86 | 17048 | 17083 | 73 | 119.8 | 428 |
| 223 | MD75 | 14534 | 14600 | 137 | 117 | 51 |
| 228 | MD50 | 8824 | 8870 | 94 | 115.8 | 79 |
| 234 | MD129 | 25598 | 25642 | 89 | 115.6 | 354 |
| **Reverse Orientation** | | | | | | |
| 6 | MR61 | 19088 | 19132 | 88 | 197.8 | 271 |
| 10 | MR72 | 23563 | 23636 | 148 | 188.8 | 286 |
| 11 | MR11 | 3775 | 3844 | 136 | 185.1 | 116 |
| 12 | MR94 | 29532 | 29582 | 94 | 184.6 | 271 |

| | | | | | | |
|---|---|---|---|---|---|---|
| 15 | MR43 | 14973 | 15028 | 109 | 183.9 | 226 |
| 27 | MR14 | 4160 | 4206 | 89 | 170 | 241 |
| 34 | MR35 | 11734 | 11792 | 111 | 164.2 | 37 |
| 52 | MR5 | 1603 | 1652 | 89 | 152.7 | 118 |
| 53 | MR57 | 18089 | 18132 | 101 | 152.7 | 139 |
| 94 | MR8 | 2804 | 2864 | 122 | 137.4 | 38 |
| 107 | MR58 | 18474 | 18508 | 72 | 134.9 | 237 |
| 117 | MR16 | 4506 | 4540 | 72 | 133.8 | 311 |
| 120 | MR34 | 10010 | 10048 | 82 | 132.7 | 245 |
| 133 | MR7 | 2534 | 2578 | 90 | 130.4 | 75 |
| 146 | MR79 | 24766 | 24808 | 75 | 127.9 | 59 |
| 150 | MR65 | 21528 | 21576 | 99 | 127.4 | 83 |
| 180 | MR60 | 19016 | 19049 | 70 | 122.5 | 72 |
| 187 | MR51 | 16450 | 16482 | 75 | 121 | 363 |
| 190 | MR80 | 25687 | 25734 | 96 | 120.6 | 75 |
| 198 | MR64 | 21507 | 21544 | 70 | 120.3 | 35 |

| | | | | | | |
|---|---|---|---|---|---|---|
| 206 | MR41 | 14500 | 14542 | 84 | 119.2 | 94 |
| 218 | MR84 | 26840 | 26894 | 108 | 117.6 | 94 |

**Sheet_2**

**List of stable stem-loop structures based on MFE.**

| **Rank** | **Name** | **Start** | **Apex** | **Size** | **Score** | **Window Count (Absolute)** | **MFE (kcal/mol )** |
|---|---|---|---|---|---|---|---|
| **Direct Orientation** | | | | | | | |
| 1 | MD13 | 2801 | 2864 | 125 | 243.8 | 61 | -40.7 |
| 2 | MD62 | 11234 | 11286 | 101 | 211.4 | 49 | -36.7 |
| 4 | MD136 | 27666 | 27721 | 104 | 205.6 | 119 | -25.3 |
| 5 | MD108 | 21131 | 21184 | 110 | 204.7 | 210 | -38.6 |
| 9 | MD132 | 26743 | 26801 | 119 | 188.9 | 252 | -39.9 |
| 19 | MD56 | 9797 | 9858 | 128 | 179.1 | 59 | -35 |
| 26 | MD139 | 28196 | 28233 | 72 | 170.4 | 133 | -18.2 |
| 28 | MD16 | 2934 | 2974 | 76 | 169.9 | 71 | -24.1 |
| 43 | MD103 | 20002 | 20042 | 80 | 159.3 | 403 | -21.3 |
| 46 | MD6 | 1489 | 1531 | 86 | 156.7 | 171 | -26.1 |
| 51 | MD17 | 2981 | 3047 | 131 | 152.8 | 38 | -40.8 |
| 87 | MD4 | 651 | 692 | 75 | 140.3 | 46 | -28.1 |
| 95 | MD7 | 1810 | 1872 | 121 | 137.4 | 58 | -34.6 |
| 122 | MD55 | 9712 | 9758 | 96 | 132.5 | 49 | -24.7 |
| 135 | MD70 | 13171 | 13219 | 93 | 130.2 | 131 | -30.5 |
| 164 | MD95 | 18782 | 18820 | 79 | 124.7 | 184 | -20.8 |
| 173 | MD121 | 24086 | 24135 | 99 | 123.1 | 45 | -24.4 |
| 176 | MD96 | 19046 | 19086 | 75 | 123.1 | 179 | -20.9 |
| 196 | MD19 | 3197 | 3236 | 76 | 120.4 | 49 | -20.1 |
| 200 | MD86 | 17048 | 17083 | 73 | 119.8 | 428 | -30.1 |
| 223 | MD75 | 14534 | 14600 | 137 | 117 | 51 | -34.5 |
| 228 | MD50 | 8824 | 8870 | 94 | 115.8 | 79 | -37.3 |

| | | | | | | | |
|---|---|---|---|---|---|---|---|
| 234 | MD129 | 25598 | 25642 | 89 | 115.6 | 354 | -26 |
| **Reverse Orientation** | | | | | | | |
| 6 | MR61 | 19088 | 19132 | 88 | 197.8 | 271 | -23.7 |
| 10 | MR72 | 23563 | 23636 | 148 | 188.8 | 286 | -44 |
| 11 | MR11 | 3775 | 3844 | 136 | 185.1 | 116 | -27.1 |
| 12 | MR94 | 29532 | 29582 | 94 | 184.6 | 271 | -28.8 |
| 15 | MR43 | 14973 | 15028 | 109 | 183.9 | 226 | -28.8 |
| 27 | MR14 | 4160 | 4206 | 89 | 170 | 241 | -26.6 |
| 34 | MR35 | 11734 | 11792 | 111 | 164.2 | 37 | -34.4 |
| 52 | MR5 | 1603 | 1652 | 89 | 152.7 | 118 | -25.9 |
| 53 | MR57 | 18089 | 18132 | 101 | 152.7 | 139 | -32.5 |
| 94 | MR8 | 2804 | 2864 | 122 | 137.4 | 38 | -32.5 |
| 117 | MR16 | 4506 | 4540 | 72 | 133.8 | 311 | -19.6 |
| 120 | MR34 | 10010 | 10048 | 82 | 132.7 | 245 | -21.7 |
| 133 | MR7 | 2534 | 2578 | 90 | 130.4 | 75 | -26.7 |
| 146 | MR79 | 24766 | 24808 | 75 | 127.9 | 59 | -25.3 |
| 187 | MR51 | 16450 | 16482 | 75 | 121 | 363 | -20.2 |
| 190 | MR80 | 25687 | 25734 | 96 | 120.6 | 75 | -20.5 |
| 206 | MR41 | 14500 | 14542 | 84 | 119.2 | 94 | -19.8 |
| 218 | MR84 | 26840 | 26894 | 108 | 117.6 | 94 | -31.2 |

**Sheet_3**

**List of stem-loop structures differentiated from other non-coding RNAs based on MFEI.**

| Name | Orientation | Start* | Apex | Size | Score | Window Count (Absolute) | MFE (kcal/mol ) | G+C % | MFEI (kcal/mol ) | Sequence |
|---|---|---|---|---|---|---|---|---|---|---|
| MD50 | Direct | 8824 | 8870 | 94 | 115.8 | 79 | -37.3 | 45 | -0.8818 | GAUUGCUGCAGUCAUAACAAGAGAAGUGGGUUUUGUCGUGCCUGGUUUGCCUGGCACGAUAUUACGCACAACUAAUGGUGACUUUUUGCAUUUC |
| MD86 | Direct | 17048 | 17083 | 73 | 119.8 | 428 | -30.1 | 45 | -0.9163 | AGGUUGGUAUGCAAAAGUAUUCUACACUCCAGGGACCACCUGGUACUGGUAAGAGUCAUUUUGCUAUUGGCCU |
| MD62 | Direct | 11234 | 11286 | 101 | 211.4 | 49 | -36.7 | 39 | -0.9318 | GCUAGUUGGGUGAUGCGUAUUAUGACAUGGUUGGAUAUGGUUGAUACUAGUUUGUCUGGUUUUAAGCUAAAAGACUGUGUUAUGUAUGCAUCAGCUGUAGU |
| MR5 | Reverse | 1603 | 1652 | 89 | 152.7 | 118 | -25.9 | 31 | -0.9387 | UAAUGGCGAUCUCUUCAUUAAGUUUAAAGUCACCAACAAUAUUGAUGUUGACUUUCUCUUUUUGGAGUAUUUCAAGAAGGUUGUCAUUA |

* start position w.r.t to the first nucleotide of the SARS-CoV-2 genome

**Sheet_4**

**List of putative targets of each SARS-CoV-2 miRNA as  predited by miRDB.**

| miRNA Name | Target Rank | Target Score | Gene Symbol | Gene Description |
|---|---|---|---|---|
| MD50-5p | 1 | 97 | GPNMB | glycoprotein nmb |
| | 2 | 95 | CAMK1D | calcium/calmodulin dependent protein kinase ID |
| | 3 | 94 | IST1 | IST1, ESCRT-III associated factor |
| | 4 | 94 | TMEM120A | transmembrane protein 120A |
| | 5 | 93 | LGI3 | leucine rich repeat LGI family member 3 |
| | 6 | 93 | ACSBG1 | acyl-CoA synthetase bubblegum family member 1 |
| | 7 | 92 | KCTD20 | potassium channel tetramerization domain containing 20 |
| | 8 | 92 | CCDC69 | coiled-coil domain containing 69 |
| | 9 | 92 | FHL2 | four and a half LIM domains 2 |
| | 10 | 92 | GNL3 | G protein nucleolar 3 |
| | 11 | 89 | TMEM127 | transmembrane protein 127 |
| | 12 | 89 | RAB5B | RAB5B, member RAS oncogene family |
| | 13 | 89 | TRIM9 | tripartite motif containing 9 |
| | 14 | 89 | DCX | doublecortin |

| | | | | |
|---|---|---|---|---|
| | 15 | 89 | ZNF83 | zinc finger protein 83 |
| | 16 | 88 | FRMD8 | FERM domain containing 8 |
| | 17 | 88 | ELAVL4 | ELAV like RNA binding protein 4 |
| | 18 | 88 | PPM1M | protein phosphatase, Mg2+/Mn2+ dependent 1M |
| | 19 | 87 | NAA40 | N(alpha)-acetyltransferase 40, NatD catalytic subunit |
| | 20 | 87 | F2RL3 | F2R like thrombin or trypsin receptor 3 |
| | 21 | 87 | ABRAXAS1 | abraxas 1, BRCA1 A complex subunit |
| | 22 | 87 | NYX | nyctalopin |
| | 23 | 87 | ADAMTS6 | ADAM metallopeptidase with thrombospondin type 1 motif 6 |
| | 24 | 87 | TFCP2 | transcription factor CP2 |
| | 25 | 86 | DLX3 | distal-less homeobox 3 |
| | 26 | 86 | CCR2 | C-C motif chemokine receptor 2 |
| | 27 | 86 | SV2A | synaptic vesicle glycoprotein 2A |
| | 28 | 86 | IGF2BP3 | insulin like growth factor 2 mRNA binding protein 3 |
| | 29 | 86 | IGF2BP1 | insulin like growth factor 2 mRNA binding protein 1 |
| | 30 | 85 | CPNE8 | copine 8 |

| | | | | |
|---|---|---|---|---|
| | 31 | 85 | NEK9 | NIMA related kinase 9 |
| | 32 | 85 | TNFSF4 | TNF superfamily member 4 |
| | 33 | 85 | SAMD9 | sterile alpha motif domain containing 9 |
| | 34 | 85 | DMP1 | dentin matrix acidic phosphoprotein 1 |
| | 35 | 84 | RPL10 | ribosomal protein L10 |
| | 36 | 84 | XPNPEP3 | X-prolyl aminopeptidase 3 |
| | 37 | 84 | WDR4 | WD repeat domain 4 |
| | 38 | 84 | DUSP3 | dual specificity phosphatase 3 |
| | 39 | 84 | FZD8 | frizzled class receptor 8 |
| | 40 | 83 | DRAXIN | dorsal inhibitory axon guidance protein |
| | 41 | 83 | WDR93 | WD repeat domain 93 |
| | 42 | 83 | GOLGA6A | golgin A6 family member A |
| | 43 | 83 | GOLGA6B | golgin A6 family member B |
| | 44 | 83 | OTX2 | orthodenticle homeobox 2 |
| | 45 | 83 | YPEL4 | yippee like 4 |
| | 46 | 83 | TTC28 | tetratricopeptide repeat domain 28 |

| | | | | |
|---|---|---|---|---|
| | 47 | 82 | TKFC | triokinase and FMN cyclase |
| | 48 | 82 | GOLGA6D | golgin A6 family member D |
| | 49 | 82 | WT1 | Wilms tumor 1 |
| | 50 | 82 | BPTF | bromodomain PHD finger transcription factor |
| | 51 | 82 | ANKS3 | ankyrin repeat and sterile alpha motif domain containing 3 |
| | 52 | 82 | SCUBE3 | signal peptide, CUB domain and EGF like domain containing 3 |
| | 53 | 82 | PPP1R16B | protein phosphatase 1 regulatory subunit 16B |
| | 54 | 81 | BTNL3 | butyrophilin like 3 |
| | 55 | 81 | SLC25A38 | solute carrier family 25 member 38 |
| | 56 | 81 | CNOT11 | CCR4-NOT transcription complex subunit 11 |
| | 57 | 81 | ZNF43 | zinc finger protein 43 |
| | 58 | 81 | CARD14 | caspase recruitment domain family member 14 |
| | 59 | 81 | KCNQ5 | potassium voltage-gated channel subfamily Q member 5 |
| | 60 | 81 | RGMB | repulsive guidance molecule BMP co-receptor b |
| | 61 | 81 | DISC1 | DISC1 scaffold protein |
| | 62 | 81 | PLEKHA6 | pleckstrin homology domain containing A6 |

| | | | | |
|---|---|---|---|---|
| | 63 | 80 | EEF2KMT | eukaryotic elongation factor 2 lysine methyltransferase |
| | 64 | 80 | FZD4 | frizzled class receptor 4 |
| | 65 | 80 | MECP2 | methyl-CpG binding protein 2 |
| | 66 | 80 | BAZ2A | bromodomain adjacent to zinc finger domain 2A |
| | 67 | 80 | GSPT1 | G1 to S phase transition 1 |
| | 68 | 80 | SRSF7 | serine and arginine rich splicing factor 7 |
| | 69 | 80 | NGEF | neuronal guanine nucleotide exchange factor |
| | 70 | 80 | SCN2B | sodium voltage-gated channel beta subunit 2 |
| | 71 | 80 | GOLGA6C | golgin A6 family member C |
| | 72 | 80 | GJC1 | gap junction protein gamma 1 |
| | 73 | 80 | CACNG8 | calcium voltage-gated channel auxiliary subunit gamma 8 |
| MD50-3p | 1 | 91 | ZFAND5 | zinc finger AN1-type containing 5 |
| | 2 | 85 | PRICKLE2 | prickle planar cell polarity protein 2 |
| MD86-5p | 1 | 100 | TNPO1 | transportin 1 |

| | | | | |
|---|---|---|---|---|
| | 2 | 100 | HIPK3 | homeodomain interacting protein kinase 3 |
| | 3 | 100 | SLC30A5 | solute carrier family 30 member 5 |
| | 4 | 100 | LCOR | ligand dependent nuclear receptor corepressor |
| | 5 | 100 | KBTBD8 | kelch repeat and BTB domain containing 8 |
| | 6 | 100 | HELZ | helicase with zinc finger |
| | 7 | 100 | RFX7 | regulatory factor X7 |
| | 8 | 99 | SCAI | suppressor of cancer cell invasion |
| | 9 | 99 | TRIO | trio Rho guanine nucleotide exchange factor |
| | 10 | 99 | FRMD4B | FERM domain containing 4B |
| | 11 | 99 | TENT4B | terminal nucleotidyltransferase 4B |
| | 12 | 99 | BRWD3 | bromodomain and WD repeat domain containing 3 |
| | 13 | 99 | NUP58 | nucleoporin 58 |
| | 14 | 99 | DACH1 | dachshund family transcription factor 1 |
| | 15 | 99 | RIMS2 | regulating synaptic membrane exocytosis 2 |
| | 16 | 99 | USF3 | upstream transcription factor family member 3 |
| | 17 | 99 | CAND1 | cullin associated and neddylation dissociated 1 |

| | | | | |
|---|---|---|---|---|
| | 18 | 99 | ARHGEF7 | Rho guanine nucleotide exchange factor 7 |
| | 19 | 99 | PELI1 | pellino E3 ubiquitin protein ligase 1 |
| | 20 | 98 | LRRC8B | leucine rich repeat containing 8 VRAC subunit B |
| | 21 | 98 | CPNE4 | copine 4 |
| | 22 | 98 | SP8 | Sp8 transcription factor |
| | 23 | 98 | IGF2BP2 | insulin like growth factor 2 mRNA binding protein 2 |
| | 24 | 98 | TBX4 | T-box 4 |
| | 25 | 98 | RRM2B | ribonucleotide reductase regulatory TP53 inducible subunit M2B |
| | 26 | 98 | B3GNT2 | UDP-GlcNAc:betaGal beta-1,3-N-acetylglucosaminyltransferase 2 |
| | 27 | 98 | SMURF2 | SMAD specific E3 ubiquitin protein ligase 2 |
| | 28 | 98 | TAOK1 | TAO kinase 1 |
| | 29 | 98 | CCNT2 | cyclin T2 |
| | 30 | 98 | OLFML3 | olfactomedin like 3 |
| | 31 | 98 | HAP1 | huntingtin associated protein 1 |
| | 32 | 98 | TSHZ2 | teashirt zinc finger homeobox 2 |
| | 33 | 98 | ASXL2 | ASXL transcriptional regulator 2 |

| | | | | |
|---|---|---|---|---|
| | 34 | 98 | PPP3R1 | protein phosphatase 3 regulatory subunit B, alpha |
| | 35 | 98 | UBR3 | ubiquitin protein ligase E3 component n-recognin 3 |
| | 36 | 98 | MYNN | myoneurin |
| | 37 | 97 | C11orf96 | chromosome 11 open reading frame 96 |
| | 38 | 97 | SLC30A4 | solute carrier family 30 member 4 |
| | 39 | 97 | ZNF761 | zinc finger protein 761 |
| | 40 | 97 | SENP2 | SUMO specific peptidase 2 |
| | 41 | 97 | BNC2 | basonuclin 2 |
| | 42 | 97 | TRDN | triadin |
| | 43 | 97 | TNFRSF19 | TNF receptor superfamily member 19 |
| | 44 | 97 | GPATCH2L | G-patch domain containing 2 like |
| | 45 | 97 | ELOC | elongin C |
| | 46 | 97 | TSNAX | translin associated factor X |
| | 47 | 97 | CDK2AP1 | cyclin dependent kinase 2 associated protein 1 |
| | 48 | 97 | DPP10 | dipeptidyl peptidase like 10 |
| | 49 | 97 | TARDBP | TAR DNA binding protein |

| | | | | |
|---|---|---|---|---|
| | 50 | 97 | SEC14L1 | SEC14 like lipid binding 1 |
| | 51 | 97 | MFSD14A | major facilitator superfamily domain containing 14A |
| | 52 | 97 | RAB2A | RAB2A, member RAS oncogene family |
| | 53 | 97 | EGR1 | early growth response 1 |
| | 54 | 97 | RHAG | Rh associated glycoprotein |
| | 55 | 97 | VWC2 | von Willebrand factor C domain containing 2 |
| | 56 | 96 | ANO4 | anoctamin 4 |
| | 57 | 96 | CREBRF | CREB3 regulatory factor |
| | 58 | 96 | NKAIN2 | sodium/potassium transporting ATPase interacting 2 |
| | 59 | 96 | PRKAR1A | protein kinase cAMP-dependent type I regulatory subunit alpha |
| | 60 | 96 | MGAT2 | mannosyl (alpha-1,6-)-glycoprotein beta-1,2-N-acetylglucosaminyltransferase |
| | 61 | 96 | GNB4 | G protein subunit beta 4 |
| | 62 | 96 | B3GNT5 | UDP-GlcNAc:betaGal beta-1,3-N-acetylglucosaminyltransferase 5 |
| | 63 | 96 | RFX3 | regulatory factor X3 |
| | 64 | 96 | TRA2B | transformer 2 beta homolog |
| | 65 | 96 | RBM46 | RNA binding motif protein 46 |

| | | | | |
|---|---|---|---|---|
| | 66 | 96 | TRPS1 | transcriptional repressor GATA binding 1 |
| | 67 | 96 | SGK1 | serum/glucocorticoid regulated kinase 1 |
| | 68 | 96 | SLC12A1 | solute carrier family 12 member 1 |
| | 69 | 96 | ROBO1 | roundabout guidance receptor 1 |
| | 70 | 96 | MYBL1 | MYB proto-oncogene like 1 |
| | 71 | 96 | FUT4 | fucosyltransferase 4 |
| | 72 | 96 | TRPC1 | transient receptor potential cation channel subfamily C member 1 |
| | 73 | 96 | TCF12 | transcription factor 12 |
| | 74 | 96 | LUM | lumican |
| | 75 | 96 | FRS2 | fibroblast growth factor receptor substrate 2 |
| | 76 | 96 | PJA2 | praja ring finger ubiquitin ligase 2 |
| | 77 | 96 | ANKRD44 | ankyrin repeat domain 44 |
| | 78 | 96 | CD247 | CD247 molecule |
| | 79 | 96 | RASSF6 | Ras association domain family member 6 |
| | 80 | 96 | IVNS1ABP | influenza virus NS1A binding protein |
| | 81 | 96 | ACTL6A | actin like 6A |

| | | | | |
|---|---|---|---|---|
| | 82 | 95 | DPM1 | dolichyl-phosphate mannosyltransferase subunit 1, catalytic |
| | 83 | 95 | SNX16 | sorting nexin 16 |
| | 84 | 95 | MAMDC2 | MAM domain containing 2 |
| | 85 | 95 | CTNND1 | catenin delta 1 |
| | 86 | 95 | USP19 | ubiquitin specific peptidase 19 |
| | 87 | 95 | PTGR2 | prostaglandin reductase 2 |
| | 88 | 95 | ZNF813 | zinc finger protein 813 |
| | 89 | 95 | DMD | dystrophin |
| | 90 | 95 | TMEFF2 | transmembrane protein with EGF like and two follistatin like domains 2 |
| | 91 | 95 | GABRB3 | gamma-aminobutyric acid type A receptor beta3 subunit |
| | 92 | 95 | ZBTB11 | zinc finger and BTB domain containing 11 |
| | 93 | 95 | SBNO1 | strawberry notch homolog 1 |
| | 94 | 95 | KCNMA1 | potassium calcium-activated channel subfamily M alpha 1 |
| | 95 | 95 | ZBTB18 | zinc finger and BTB domain containing 18 |
| | 96 | 95 | SLF2 | SMC5-SMC6 complex localization factor 2 |
| | 97 | 95 | KMT5A | lysine methyltransferase 5A |

| | | | | |
|---|---|---|---|---|
| | 98 | 95 | CSMD3 | CUB and Sushi multiple domains 3 |
| | 99 | 95 | EEA1 | early endosome antigen 1 |
| | 100 | 95 | PTPRM | protein tyrosine phosphatase, receptor type M |
| | 101 | 94 | NEXN | nexilin F-actin binding protein |
| | 102 | 94 | VPS35 | VPS35, retromer complex component |
| | 103 | 94 | SCML4 | Scm polycomb group protein like 4 |
| | 104 | 94 | MKNK1 | MAP kinase interacting serine/threonine kinase 1 |
| | 105 | 94 | VSNL1 | visinin like 1 |
| | 106 | 94 | ID3 | inhibitor of DNA binding 3, HLH protein |
| | 107 | 94 | ARMC1 | armadillo repeat containing 1 |
| | 108 | 94 | LRP6 | LDL receptor related protein 6 |
| | 109 | 94 | CACNA1C | calcium voltage-gated channel subunit alpha1 C |
| | 110 | 94 | ZNF24 | zinc finger protein 24 |
| | 111 | 94 | LIMCH1 | LIM and calponin homology domains 1 |
| | 112 | 94 | PARP9 | poly(ADP-ribose) polymerase family member 9 |
| | 113 | 94 | RXFP1 | relaxin family peptide receptor 1 |

| | | | | |
|---|---|---|---|---|
| | 114 | 94 | SMAD5 | SMAD family member 5 |
| | 115 | 94 | PBRM1 | polybromo 1 |
| | 116 | 94 | EIF4ENIF1 | eukaryotic translation initiation factor 4E nuclear import factor 1 |
| | 117 | 94 | TRIM10 | tripartite motif containing 10 |
| | 118 | 94 | PDCD6IP | programmed cell death 6 interacting protein |
| | 119 | 94 | RCN2 | reticulocalbin 2 |
| | 120 | 94 | ZNF532 | zinc finger protein 532 |
| | 121 | 93 | TRAM1L1 | translocation associated membrane protein 1 like 1 |
| | 122 | 93 | CEP55 | centrosomal protein 55 |
| | 123 | 93 | GABRB2 | gamma-aminobutyric acid type A receptor beta2 subunit |
| | 124 | 93 | DCUN1D4 | defective in cullin neddylation 1 domain containing 4 |
| | 125 | 93 | SPIN1 | spindlin 1 |
| | 126 | 93 | KLF4 | Kruppel like factor 4 |
| | 127 | 93 | KCNC2 | potassium voltage-gated channel subfamily C member 2 |
| | 128 | 93 | PSMA2 | proteasome subunit alpha 2 |
| | 129 | 93 | EDEM1 | ER degradation enhancing alpha-mannosidase like protein 1 |

| | | | | |
|---|---|---|---|---|
| | 130 | 93 | FGF14 | fibroblast growth factor 14 |
| | 131 | 93 | CLIP1 | CAP-Gly domain containing linker protein 1 |
| | 132 | 93 | DUSP6 | dual specificity phosphatase 6 |
| | 133 | 93 | SOCS2 | suppressor of cytokine signaling 2 |
| | 134 | 93 | RHOXF2 | Rhox homeobox family member 2 |
| | 135 | 93 | RHOXF2B | Rhox homeobox family member 2B |
| | 136 | 93 | CDK14 | cyclin dependent kinase 14 |
| | 137 | 93 | PDS5A | PDS5 cohesin associated factor A |
| | 138 | 93 | SCN8A | sodium voltage-gated channel alpha subunit 8 |
| | 139 | 93 | BRWD1 | bromodomain and WD repeat domain containing 1 |
| | 140 | 93 | PTCH1 | patched 1 |
| | 141 | 93 | RDX | radixin |
| | 142 | 93 | VPS13B | vacuolar protein sorting 13 homolog B |
| | 143 | 93 | NECAB3 | N-terminal EF-hand calcium binding protein 3 |
| | 144 | 93 | STXBP5L | syntaxin binding protein 5 like |
| | 145 | 92 | KLHL3 | kelch like family member 3 |

| | | | | |
|---|---|---|---|---|
| | 146 | 92 | AMD1 | adenosylmethionine decarboxylase 1 |
| | 147 | 92 | PRPF39 | pre-mRNA processing factor 39 |
| | 148 | 92 | GXYLT1 | glucoside xylosyltransferase 1 |
| | 149 | 92 | TCEA1 | transcription elongation factor A1 |
| | 150 | 92 | PFDN4 | prefoldin subunit 4 |
| | 151 | 92 | CNNM4 | cyclin and CBS domain divalent metal cation transport mediator 4 |
| | 152 | 92 | IMPAD1 | inositol monophosphatase domain containing 1 |
| | 153 | 92 | SLC9B2 | solute carrier family 9 member B2 |
| | 154 | 92 | CDH7 | cadherin 7 |
| | 155 | 92 | HOOK1 | hook microtubule tethering protein 1 |
| | 156 | 92 | STC1 | stanniocalcin 1 |
| | 157 | 92 | KIAA1841 | KIAA1841 |
| | 158 | 92 | LILRB4 | leukocyte immunoglobulin like receptor B4 |
| | 159 | 92 | GPX7 | glutathione peroxidase 7 |
| | 160 | 92 | SORBS2 | sorbin and SH3 domain containing 2 |
| | 161 | 92 | GATA6 | GATA binding protein 6 |

| | | | | |
|---|---|---|---|---|
| | 162 | 92 | TFAP2C | transcription factor AP-2 gamma |
| | 163 | 92 | PUM1 | pumilio RNA binding family member 1 |
| | 164 | 92 | ATXN7L3 | ataxin 7 like 3 |
| | 165 | 92 | ITSN2 | intersectin 2 |
| | 166 | 92 | KLF7 | Kruppel like factor 7 |
| | 167 | 92 | CLOCK | clock circadian regulator |
| | 168 | 92 | ANKRD13A | ankyrin repeat domain 13A |
| | 169 | 92 | MAPK8 | mitogen-activated protein kinase 8 |
| | 170 | 92 | BCAT1 | branched chain amino acid transaminase 1 |
| | 171 | 92 | KITLG | KIT ligand |
| | 172 | 92 | NRXN1 | neurexin 1 |
| | 173 | 91 | L3MBTL3 | L3MBTL3, histone methyl-lysine binding protein |
| | 174 | 91 | PCDH7 | protocadherin 7 |
| | 175 | 91 | PHF6 | PHD finger protein 6 |
| | 176 | 91 | SLC5A3 | solute carrier family 5 member 3 |
| | 177 | 91 | FOXJ3 | forkhead box J3 |

| | | | | |
|---|---|---|---|---|
| | 178 | 91 | TCEAL1 | transcription elongation factor A like 1 |
| | 179 | 91 | CELF4 | CUGBP Elav-like family member 4 |
| | 180 | 91 | PIAS2 | protein inhibitor of activated STAT 2 |
| | 181 | 91 | PRKCI | protein kinase C iota |
| | 182 | 91 | CCR2 | C-C motif chemokine receptor 2 |
| | 183 | 91 | BNIP2 | BCL2 interacting protein 2 |
| | 184 | 91 | SPTY2D1 | SPT2 chromatin protein domain containing 1 |
| | 185 | 91 | SLC9C1 | solute carrier family 9 member C1 |
| | 186 | 91 | HNRNPLL | heterogeneous nuclear ribonucleoprotein L like |
| | 187 | 91 | STUM | stum, mechanosensory transduction mediator homolog |
| | 188 | 91 | IL6ST | interleukin 6 signal transducer |
| | 189 | 91 | GRIK1 | glutamate ionotropic receptor kainate type subunit 1 |
| | 190 | 91 | FOXJ2 | forkhead box J2 |
| | 191 | 91 | ENTPD1 | ectonucleoside triphosphate diphosphohydrolase 1 |
| | 192 | 91 | AGGF1 | angiogenic factor with G-patch and FHA domains 1 |
| | 193 | 91 | IL1A | interleukin 1 alpha |

| | | | | |
|---|---|---|---|---|
| | 194 | 91 | CLRN1 | clarin 1 |
| | 195 | 91 | C20orf194 | chromosome 20 open reading frame 194 |
| | 196 | 90 | HACE1 | HECT domain and ankyrin repeat containing E3 ubiquitin protein ligase 1 |
| | 197 | 90 | RBM26 | RNA binding motif protein 26 |
| | 198 | 90 | TNN | tenascin N |
| | 199 | 90 | DYRK1A | dual specificity tyrosine phosphorylation regulated kinase 1A |
| | 200 | 90 | MAEA | macrophage erythroblast attacher |
| | 201 | 90 | C2orf76 | chromosome 2 open reading frame 76 |
| | 202 | 90 | ESPN | espin |
| | 203 | 90 | C16orf74 | chromosome 16 open reading frame 74 |
| | 204 | 90 | CHFR | checkpoint with forkhead and ring finger domains |
| | 205 | 90 | INTS6L | integrator complex subunit 6 like |
| | 206 | 90 | KPNA4 | karyopherin subunit alpha 4 |
| | 207 | 90 | NANP | N-acetylneuraminic acid phosphatase |
| | 208 | 90 | PPP2R2D | protein phosphatase 2 regulatory subunit Bdelta |
| | 209 | 90 | SEPSECS | Sep (O-phosphoserine) tRNA:Sec (selenocysteine) tRNA synthase |

| | | | | |
|---|---|---|---|---|
| | 210 | 90 | BMPR2 | bone morphogenetic protein receptor type 2 |
| | 211 | 90 | NHLH2 | nescient helix-loop-helix 2 |
| | 212 | 90 | PPP2R3A | protein phosphatase 2 regulatory subunit B"alpha |
| | 213 | 90 | PRSS23 | serine protease 23 |
| | 214 | 90 | MINDY2 | MINDY lysine 48 deubiquitinase 2 |
| | 215 | 90 | MFSD8 | major facilitator superfamily domain containing 8 |
| | 216 | 90 | PHF14 | PHD finger protein 14 |
| | 217 | 90 | PCSK5 | proprotein convertase subtilisin/kexin type 5 |
| | 218 | 90 | RCOR3 | REST corepressor 3 |
| | 219 | 90 | SRC | SRC proto-oncogene, non-receptor tyrosine kinase |
| | 220 | 90 | MTCL1 | microtubule crosslinking factor 1 |
| | 221 | 90 | DDHD2 | DDHD domain containing 2 |
| | 222 | 90 | RBPMS2 | RNA binding protein, mRNA processing factor 2 |
| | 223 | 90 | GHITM | growth hormone inducible transmembrane protein |
| | 224 | 89 | RNF212 | ring finger protein 212 |
| | 225 | 89 | SMAP1 | small ArfGAP 1 |

| | | | | |
|---|---|---|---|---|
| | 226 | 89 | CNTN5 | contactin 5 |
| | 227 | 89 | GEMIN8 | gem nuclear organelle associated protein 8 |
| | 228 | 89 | RAMAC | RNA guanine-7 methyltransferase activating subunit |
| | 229 | 89 | MARCHF2 | membrane associated ring-CH-type finger 2 |
| | 230 | 89 | TBL1XR1 | transducin beta like 1 X-linked receptor 1 |
| | 231 | 89 | BMF | Bcl2 modifying factor |
| | 232 | 89 | RSBN1 | round spermatid basic protein 1 |
| | 233 | 89 | ANTXR2 | ANTXR cell adhesion molecule 2 |
| | 234 | 89 | TSG101 | tumor susceptibility 101 |
| | 235 | 89 | G0S2 | G0/G1 switch 2 |
| | 236 | 89 | BMT2 | base methyltransferase of 25S rRNA 2 homolog |
| | 237 | 89 | CTH | cystathionine gamma-lyase |
| | 238 | 89 | YTHDF3 | YTH N6-methyladenosine RNA binding protein 3 |
| | 239 | 89 | PLPP4 | phospholipid phosphatase 4 |
| | 240 | 89 | HARBI1 | harbinger transposase derived 1 |
| | 241 | 89 | RBMS2 | RNA binding motif single stranded interacting protein 2 |

| | | | | |
|---|---|---|---|---|
| | 242 | 89 | TFRC | transferrin receptor |
| | 243 | 89 | GHSR | growth hormone secretagogue receptor |
| | 244 | 89 | DUSP4 | dual specificity phosphatase 4 |
| | 245 | 89 | SLC16A7 | solute carrier family 16 member 7 |
| | 246 | 89 | PPP1R1C | protein phosphatase 1 regulatory inhibitor subunit 1C |
| | 247 | 89 | RELCH | RAB11 binding and LisH domain, coiled-coil and HEAT repeat containing |
| | 248 | 89 | CPEB2 | cytoplasmic polyadenylation element binding protein 2 |
| | 249 | 89 | MOCS2 | molybdenum cofactor synthesis 2 |
| | 250 | 89 | SLC15A2 | solute carrier family 15 member 2 |
| | 251 | 89 | VCPIP1 | valosin containing protein interacting protein 1 |
| | 252 | 88 | C5orf24 | chromosome 5 open reading frame 24 |
| | 253 | 88 | IFT74 | intraflagellar transport 74 |
| | 254 | 88 | NEDD4L | neural precursor cell expressed, developmentally down-regulated 4-like, E3 ubiquitin protein ligase |
| | 255 | 88 | ITGAM | integrin subunit alpha M |
| | 256 | 88 | HOXC5 | homeobox C5 |
| | 257 | 88 | ATP2B4 | ATPase plasma membrane Ca2+ transporting 4 |

| | | | | |
|---|---|---|---|---|
| | 258 | 88 | CNMD | chondromodulin |
| | 259 | 88 | KCNB2 | potassium voltage-gated channel subfamily B member 2 |
| | 260 | 88 | LEPR | leptin receptor |
| | 261 | 88 | SYT14 | synaptotagmin 14 |
| | 262 | 88 | SUZ12 | SUZ12, polycomb repressive complex 2 subunit |
| | 263 | 88 | EEF1AKMT2 | EEF1A lysine methyltransferase 2 |
| | 264 | 88 | TMEM183A | transmembrane protein 183A |
| | 265 | 88 | CTTNBP2 | cortactin binding protein 2 |
| | 266 | 88 | TRNT1 | tRNA nucleotidyl transferase 1 |
| | 267 | 88 | PNN | pinin, desmosome associated protein |
| | 268 | 88 | RGPD4 | RANBP2-like and GRIP domain containing 4 |
| | 269 | 88 | DSCAM | DS cell adhesion molecule |
| | 270 | 88 | FREM2 | FRAS1 related extracellular matrix protein 2 |
| | 271 | 88 | SNN | stannin |
| | 272 | 88 | DLX5 | distal-less homeobox 5 |
| | 273 | 88 | GABRA5 | gamma-aminobutyric acid type A receptor alpha5 subunit |

| | | | | |
|---|---|---|---|---|
| | 274 | 87 | MRC1 | mannose receptor C-type 1 |
| | 275 | 87 | USP12 | ubiquitin specific peptidase 12 |
| | 276 | 87 | VAPB | VAMP associated protein B and C |
| | 277 | 87 | CACNB2 | calcium voltage-gated channel auxiliary subunit beta 2 |
| | 278 | 87 | ZNF662 | zinc finger protein 662 |
| | 279 | 87 | BTBD7 | BTB domain containing 7 |
| | 280 | 87 | PAX4 | paired box 4 |
| | 281 | 87 | UBL3 | ubiquitin like 3 |
| | 282 | 87 | METTL8 | methyltransferase like 8 |
| | 283 | 87 | GYPA | glycophorin A (MNS blood group) |
| | 284 | 87 | HAS2 | hyaluronan synthase 2 |
| | 285 | 87 | TRIM55 | tripartite motif containing 55 |
| | 286 | 87 | SMARCD2 | SWI/SNF related, matrix associated, actin dependent regulator of chromatin, subfamily d, member 2 |
| | 287 | 87 | ZBTB20 | zinc finger and BTB domain containing 20 |
| | 288 | 87 | MPP2 | membrane palmitoylated protein 2 |
| | 289 | 87 | UQCRB | ubiquinol-cytochrome c reductase binding protein |

| | | | | |
|---|---|---|---|---|
| | 290 | 87 | ARPP19 | cAMP regulated phosphoprotein 19 |
| | 291 | 87 | RBAK | RB associated KRAB zinc finger |
| | 292 | 87 | UNC13C | unc-13 homolog C |
| | 293 | 87 | PGM2 | phosphoglucomutase 2 |
| | 294 | 86 | NCK2 | NCK adaptor protein 2 |
| | 295 | 86 | RPP14 | ribonuclease P/MRP subunit p14 |
| | 296 | 86 | CNOT2 | CCR4-NOT transcription complex subunit 2 |
| | 297 | 86 | NSUN4 | NOP2/Sun RNA methyltransferase family member 4 |
| | 298 | 86 | GNAQ | G protein subunit alpha q |
| | 299 | 86 | C5orf51 | chromosome 5 open reading frame 51 |
| | 300 | 86 | RNF219 | ring finger protein 219 |
| | 301 | 86 | PRUNE1 | prune exopolyphosphatase 1 |
| | 302 | 86 | ZEB2 | zinc finger E-box binding homeobox 2 |
| | 303 | 86 | TMEM187 | transmembrane protein 187 |
| | 304 | 86 | CFLAR | CASP8 and FADD like apoptosis regulator |
| | 305 | 86 | RAPGEF5 | Rap guanine nucleotide exchange factor 5 |

| | | | | |
|---|---|---|---|---|
| | 306 | 86 | FNDC3B | fibronectin type III domain containing 3B |
| | 307 | 86 | PLEKHA1 | pleckstrin homology domain containing A1 |
| | 308 | 86 | LPCAT2 | lysophosphatidylcholine acyltransferase 2 |
| | 309 | 86 | NAPB | NSF attachment protein beta |
| | 310 | 86 | MEGF10 | multiple EGF like domains 10 |
| | 311 | 86 | MCM8 | minichromosome maintenance 8 homologous recombination repair factor |
| | 312 | 86 | DCN | decorin |
| | 313 | 86 | ZNF224 | zinc finger protein 224 |
| | 314 | 86 | STXBP5 | syntaxin binding protein 5 |
| | 315 | 86 | C18orf63 | chromosome 18 open reading frame 63 |
| | 316 | 86 | RCE1 | Ras converting CAAX endopeptidase 1 |
| | 317 | 86 | KIAA1211 | KIAA1211 |
| | 318 | 86 | ZNF567 | zinc finger protein 567 |
| | 319 | 86 | TECRL | trans-2,3-enoyl-CoA reductase like |
| | 320 | 86 | KCNQ5 | potassium voltage-gated channel subfamily Q member 5 |
| | 321 | 86 | M6PR | mannose-6-phosphate receptor, cation dependent |

| | | | | |
|---|---|---|---|---|
| | 322 | 85 | NUP107 | nucleoporin 107 |
| | 323 | 85 | C5orf63 | chromosome 5 open reading frame 63 |
| | 324 | 85 | PYHIN1 | pyrin and HIN domain family member 1 |
| | 325 | 85 | GRIA1 | glutamate ionotropic receptor AMPA type subunit 1 |
| | 326 | 85 | TMEM196 | transmembrane protein 196 |
| | 327 | 85 | PTAR1 | protein prenyltransferase alpha subunit repeat containing 1 |
| | 328 | 85 | PRDM2 | PR/SET domain 2 |
| | 329 | 85 | UBXN2B | UBX domain protein 2B |
| | 330 | 85 | PHC3 | polyhomeotic homolog 3 |
| | 331 | 85 | CYP2E1 | cytochrome P450 family 2 subfamily E member 1 |
| | 332 | 85 | VMA21 | VMA21, vacuolar ATPase assembly factor |
| | 333 | 85 | KCNT2 | potassium sodium-activated channel subfamily T member 2 |
| | 334 | 85 | ANKDD1A | ankyrin repeat and death domain containing 1A |
| | 335 | 85 | ACSL4 | acyl-CoA synthetase long chain family member 4 |
| | 336 | 85 | CYSLTR2 | cysteinyl leukotriene receptor 2 |
| | 337 | 85 | RMND5A | required for meiotic nuclear division 5 homolog A |

| | | | | |
|---|---|---|---|---|
| | 338 | 85 | CDK6 | cyclin dependent kinase 6 |
| | 339 | 85 | ONECUT2 | one cut homeobox 2 |
| | 340 | 84 | TMEM178B | transmembrane protein 178B |
| | 341 | 84 | THAP5 | THAP domain containing 5 |
| | 342 | 84 | MOSMO | modulator of smoothened |
| | 343 | 84 | ACER3 | alkaline ceramidase 3 |
| | 344 | 84 | EMP2 | epithelial membrane protein 2 |
| | 345 | 84 | CDC42 | cell division cycle 42 |
| | 346 | 84 | FYTTD1 | forty-two-three domain containing 1 |
| | 347 | 84 | MEF2C | myocyte enhancer factor 2C |
| | 348 | 84 | RHOQ | ras homolog family member Q |
| | 349 | 84 | CHML | CHM like, Rab escort protein 2 |
| | 350 | 84 | PCMTD1 | protein-L-isoaspartate (D-aspartate) O-methyltransferase domain containing 1 |
| | 351 | 84 | PAX6 | paired box 6 |
| | 352 | 84 | ST7L | suppression of tumorigenicity 7 like |
| | 353 | 84 | CCL7 | C-C motif chemokine ligand 7 |

| | | | | |
|---|---|---|---|---|
| | 354 | 84 | ATP8A1 | ATPase phospholipid transporting 8A1 |
| | 355 | 84 | LRRTM3 | leucine rich repeat transmembrane neuronal 3 |
| | 356 | 84 | CHL1 | cell adhesion molecule L1 like |
| | 357 | 84 | ZNF660 | zinc finger protein 660 |
| | 358 | 84 | SLC35A5 | solute carrier family 35 member A5 |
| | 359 | 84 | CHMP1B | charged multivesicular body protein 1B |
| | 360 | 84 | MIGA1 | mitoguardin 1 |
| | 361 | 84 | SLC40A1 | solute carrier family 40 member 1 |
| | 362 | 84 | NEGR1 | neuronal growth regulator 1 |
| | 363 | 84 | TPST2 | tyrosylprotein sulfotransferase 2 |
| | 364 | 84 | SRPK2 | SRSF protein kinase 2 |
| | 365 | 84 | PRMT8 | protein arginine methyltransferase 8 |
| | 366 | 84 | SMN2 | survival of motor neuron 2, centromeric |
| | 367 | 83 | MACROD2 | MACRO domain containing 2 |
| | 368 | 83 | MTPN | myotrophin |
| | 369 | 83 | ANKFY1 | ankyrin repeat and FYVE domain containing 1 |

| | | | | |
|---|---|---|---|---|
| | 370 | 83 | TMEM68 | transmembrane protein 68 |
| | 371 | 83 | SMN1 | survival of motor neuron 1, telomeric |
| | 372 | 83 | KCNK1 | potassium two pore domain channel subfamily K member 1 |
| | 373 | 83 | LGR4 | leucine rich repeat containing G protein-coupled receptor 4 |
| | 374 | 83 | UBQLN2 | ubiquilin 2 |
| | 375 | 83 | ZNF827 | zinc finger protein 827 |
| | 376 | 83 | SLC39A10 | solute carrier family 39 member 10 |
| | 377 | 83 | SPOCK1 | SPARC (osteonectin), cwcv and kazal like domains proteoglycan 1 |
| | 378 | 83 | RAB7A | RAB7A, member RAS oncogene family |
| | 379 | 83 | EIF3H | eukaryotic translation initiation factor 3 subunit H |
| | 380 | 83 | ZNF492 | zinc finger protein 492 |
| | 381 | 83 | RPGRIP1L | RPGRIP1 like |
| | 382 | 83 | TRIM58 | tripartite motif containing 58 |
| | 383 | 83 | MKX | mohawk homeobox |
| | 384 | 83 | UBE2N | ubiquitin conjugating enzyme E2 N |
| | 385 | 83 | OCLN | occludin |

| | | | | |
|---|---|---|---|---|
| | 386 | 83 | CHURC1 | churchill domain containing 1 |
| | 387 | 83 | MARCKS | myristoylated alanine rich protein kinase C substrate |
| | 388 | 83 | CENPI | centromere protein I |
| | 389 | 83 | AHNAK | AHNAK nucleoprotein |
| | 390 | 83 | CPM | carboxypeptidase M |
| | 391 | 83 | SCAF11 | SR-related CTD associated factor 11 |
| | 392 | 83 | IDH2 | isocitrate dehydrogenase (NADP(+)) 2, mitochondrial |
| | 393 | 83 | MDGA2 | MAM domain containing glycosylphosphatidylinositol anchor 2 |
| | 394 | 83 | LEMD3 | LEM domain containing 3 |
| | 395 | 83 | LATS2 | large tumor suppressor kinase 2 |
| | 396 | 83 | WIPF1 | WAS/WASL interacting protein family member 1 |
| | 397 | 82 | SLC38A2 | solute carrier family 38 member 2 |
| | 398 | 82 | OLR1 | oxidized low density lipoprotein receptor 1 |
| | 399 | 82 | LRIG2 | leucine rich repeats and immunoglobulin like domains 2 |
| | 400 | 82 | ZNF418 | zinc finger protein 418 |
| | 401 | 82 | TIAM1 | T cell lymphoma invasion and metastasis 1 |

| | | | | |
|---|---|---|---|---|
| | 402 | 82 | RIMS1 | regulating synaptic membrane exocytosis 1 |
| | 403 | 82 | ZNF678 | zinc finger protein 678 |
| | 404 | 82 | SPOPL | speckle type BTB/POZ protein like |
| | 405 | 82 | CAMK2G | calcium/calmodulin dependent protein kinase II gamma |
| | 406 | 82 | CCNG2 | cyclin G2 |
| | 407 | 82 | SNRNP48 | small nuclear ribonucleoprotein U11/U12 subunit 48 |
| | 408 | 82 | ZNF195 | zinc finger protein 195 |
| | 409 | 82 | CACNA2D1 | calcium voltage-gated channel auxiliary subunit alpha2delta 1 |
| | 410 | 82 | SEC24A | SEC24 homolog A, COPII coat complex component |
| | 411 | 82 | HERC6 | HECT and RLD domain containing E3 ubiquitin protein ligase family member 6 |
| | 412 | 82 | OPRM1 | opioid receptor mu 1 |
| | 413 | 82 | ASB9 | ankyrin repeat and SOCS box containing 9 |
| | 414 | 82 | SDC2 | syndecan 2 |
| | 415 | 82 | SERBP1 | SERPINE1 mRNA binding protein 1 |
| | 416 | 82 | DUT | deoxyuridine triphosphatase |
| | 417 | 82 | HDAC9 | histone deacetylase 9 |

| | | | | |
|---|---|---|---|---|
| | 418 | 82 | ITM2C | integral membrane protein 2C |
| | 419 | 82 | TRIM66 | tripartite motif containing 66 |
| | 420 | 82 | LOC388813 | uncharacterized protein ENSP00000383407-like |
| | 421 | 81 | NBR1 | NBR1, autophagy cargo receptor |
| | 422 | 81 | GPR65 | G protein-coupled receptor 65 |
| | 423 | 81 | ZNF148 | zinc finger protein 148 |
| | 424 | 81 | SEPTIN11 | septin 11 |
| | 425 | 81 | DMRT1 | doublesex and mab-3 related transcription factor 1 |
| | 426 | 81 | HORMAD2 | HORMA domain containing 2 |
| | 427 | 81 | ZNF441 | zinc finger protein 441 |
| | 428 | 81 | PRDM1 | PR/SET domain 1 |
| | 429 | 81 | BCL11A | BCL11A, BAF complex component |
| | 430 | 81 | HMG20A | high mobility group 20A |
| | 431 | 81 | C8orf88 | chromosome 8 open reading frame 88 |
| | 432 | 81 | FBXO9 | F-box protein 9 |
| | 433 | 81 | PIGK | phosphatidylinositol glycan anchor biosynthesis class K |

| | | | | |
|---|---|---|---|---|
| | 434 | 81 | PDK3 | pyruvate dehydrogenase kinase 3 |
| | 435 | 81 | ATF1 | activating transcription factor 1 |
| | 436 | 81 | TCF3 | transcription factor 3 |
| | 437 | 81 | C12orf40 | chromosome 12 open reading frame 40 |
| | 438 | 81 | ZNF831 | zinc finger protein 831 |
| | 439 | 81 | CLMN | calmin |
| | 440 | 81 | ATXN7L3B | ataxin 7 like 3B |
| | 441 | 81 | KDM3A | lysine demethylase 3A |
| | 442 | 81 | HERC1 | HECT and RLD domain containing E3 ubiquitin protein ligase family member 1 |
| | 443 | 81 | CA3 | carbonic anhydrase 3 |
| | 444 | 81 | CDK1 | cyclin dependent kinase 1 |
| | 445 | 81 | RAB18 | RAB18, member RAS oncogene family |
| | 446 | 80 | AQP4 | aquaporin 4 |
| | 447 | 80 | ASAP2 | ArfGAP with SH3 domain, ankyrin repeat and PH domain 2 |
| | 448 | 80 | PPM1B | protein phosphatase, Mg2+/Mn2+ dependent 1B |
| | 449 | 80 | RGS10 | regulator of G protein signaling 10 |

| | | | | |
|---|---|---|---|---|
| | 450 | 80 | ARL14EP | ADP ribosylation factor like GTPase 14 effector protein |
| | 451 | 80 | KCNJ13 | potassium voltage-gated channel subfamily J member 13 |
| | 452 | 80 | HERC4 | HECT and RLD domain containing E3 ubiquitin protein ligase 4 |
| | 453 | 80 | TULP4 | tubby like protein 4 |
| | 454 | 80 | KCTD8 | potassium channel tetramerization domain containing 8 |
| | 455 | 80 | TES | testin LIM domain protein |
| | 456 | 80 | CCDC80 | coiled-coil domain containing 80 |
| | 457 | 80 | PPHLN1 | periphilin 1 |
| | 458 | 80 | CYP26B1 | cytochrome P450 family 26 subfamily B member 1 |
| | 459 | 80 | NDFIP2 | Nedd4 family interacting protein 2 |
| | 460 | 80 | EXD2 | exonuclease 3'-5' domain containing 2 |
| | 461 | 80 | DNMT3A | DNA methyltransferase 3 alpha |
| | 462 | 80 | HNRNPH3 | heterogeneous nuclear ribonucleoprotein H3 |
| | 463 | 80 | MED12L | mediator complex subunit 12 like |
| | 464 | 80 | LRP1B | LDL receptor related protein 1B |
| | 465 | 80 | G3BP1 | G3BP stress granule assembly factor 1 |

| | | | | |
|---|---|---|---|---|
| | 466 | 80 | MAP1LC3B | microtubule associated protein 1 light chain 3 beta |
| | 467 | 80 | RPL36A | ribosomal protein L36a |
| | 468 | 80 | PDE1C | phosphodiesterase 1C |
| | 469 | 80 | OSGEPL1 | O-sialoglycoprotein endopeptidase like 1 |
| | 470 | 80 | MRTFB | myocardin related transcription factor B |
| | 471 | 80 | FOXN3 | forkhead box N3 |
| | 472 | 80 | SV2B | synaptic vesicle glycoprotein 2B |
| MD86-3p | 1 | 99 | CNTNAP5 | contactin associated protein like 5 |
| | 2 | 99 | ZNF430 | zinc finger protein 430 |
| | 3 | 98 | SNX10 | sorting nexin 10 |
| | 4 | 97 | CLDND1 | claudin domain containing 1 |
| | 5 | 96 | MTX2 | metaxin 2 |
| | 6 | 95 | SKI | SKI proto-oncogene |
| | 7 | 95 | KRAS | KRAS proto-oncogene, GTPase |
| | 8 | 95 | TPGS2 | tubulin polyglutamylase complex subunit 2 |

| | | | | |
|---|---|---|---|---|
| | 9 | 95 | SP1 | Sp1 transcription factor |
| | 10 | 95 | P2RY12 | purinergic receptor P2Y12 |
| | 11 | 95 | DNAJC3 | DnaJ heat shock protein family (Hsp40) member C3 |
| | 12 | 94 | MARK1 | microtubule affinity regulating kinase 1 |
| | 13 | 94 | SERINC1 | serine incorporator 1 |
| | 14 | 94 | SNX1 | sorting nexin 1 |
| | 15 | 94 | EHBP1 | EH domain binding protein 1 |
| | 16 | 93 | C6orf62 | chromosome 6 open reading frame 62 |
| | 17 | 93 | RNF19B | ring finger protein 19B |
| | 18 | 93 | EYA3 | EYA transcriptional coactivator and phosphatase 3 |
| | 19 | 93 | TSPAN12 | tetraspanin 12 |
| | 20 | 92 | PAK5 | p21 (RAC1) activated kinase 5 |
| | 21 | 92 | ENOSF1 | enolase superfamily member 1 |
| | 22 | 92 | LIN7A | lin-7 homolog A, crumbs cell polarity complex component |
| | 23 | 91 | ABCC1 | ATP binding cassette subfamily C member 1 |
| | 24 | 91 | INPP5A | inositol polyphosphate-5-phosphatase A |

| | | | | |
|---|---|---|---|---|
| | 25 | 91 | NEXMIF | neurite extension and migration factor |
| | 26 | 91 | MRPL48 | mitochondrial ribosomal protein L48 |
| | 27 | 91 | DCBLD2 | discoidin, CUB and LCCL domain containing 2 |
| | 28 | 91 | TAPT1 | transmembrane anterior posterior transformation 1 |
| | 29 | 90 | VPS13B | vacuolar protein sorting 13 homolog B |
| | 30 | 90 | CHRNA3 | cholinergic receptor nicotinic alpha 3 subunit |
| | 31 | 90 | RBFOX3 | RNA binding fox-1 homolog 3 |
| | 32 | 90 | C1orf112 | chromosome 1 open reading frame 112 |
| | 33 | 90 | PNRC1 | proline rich nuclear receptor coactivator 1 |
| | 34 | 90 | IMP4 | IMP4, U3 small nucleolar ribonucleoprotein |
| | 35 | 89 | CYS1 | cystin 1 |
| | 36 | 89 | PCDH9 | protocadherin 9 |
| | 37 | 89 | NELL2 | neural EGFL like 2 |
| | 38 | 89 | NCOA2 | nuclear receptor coactivator 2 |
| | 39 | 88 | CHDH | choline dehydrogenase |
| | 40 | 88 | TLR8 | toll like receptor 8 |

| | | | | |
|---|---|---|---|---|
| | 41 | 88 | RASGRP3 | RAS guanyl releasing protein 3 |
| | 42 | 88 | TRIP4 | thyroid hormone receptor interactor 4 |
| | 43 | 88 | RASSF8 | Ras association domain family member 8 |
| | 44 | 88 | RPP30 | ribonuclease P/MRP subunit p30 |
| | 45 | 87 | ZNF202 | zinc finger protein 202 |
| | 46 | 87 | ZFAND1 | zinc finger AN1-type containing 1 |
| | 47 | 87 | CADM1 | cell adhesion molecule 1 |
| | 48 | 87 | GALNT3 | polypeptide N-acetylgalactosaminyltransferase 3 |
| | 49 | 87 | WEE1 | WEE1 G2 checkpoint kinase |
| | 50 | 87 | MKRN2 | makorin ring finger protein 2 |
| | 51 | 87 | CHN2 | chimerin 2 |
| | 52 | 87 | ST3GAL1 | ST3 beta-galactoside alpha-2,3-sialyltransferase 1 |
| | 53 | 87 | C2CD6 | C2 calcium dependent domain containing 6 |
| | 54 | 87 | DCAF17 | DDB1 and CUL4 associated factor 17 |
| | 55 | 87 | ANKLE2 | ankyrin repeat and LEM domain containing 2 |
| | 56 | 87 | DIPK2A | divergent protein kinase domain 2A |

| | | | | |
|---|---|---|---|---|
| | 57 | 86 | DCTN2 | dynactin subunit 2 |
| | 58 | 86 | PPM1E | protein phosphatase, Mg2+/Mn2+ dependent 1E |
| | 59 | 86 | DNAJB9 | DnaJ heat shock protein family (Hsp40) member B9 |
| | 60 | 86 | DMXL2 | Dmx like 2 |
| | 61 | 86 | DOCK3 | dedicator of cytokinesis 3 |
| | 62 | 85 | TVP23A | trans-golgi network vesicle protein 23 homolog A |
| | 63 | 85 | DPYD | dihydropyrimidine dehydrogenase |
| | 64 | 85 | USP49 | ubiquitin specific peptidase 49 |
| | 65 | 85 | GABPA | GA binding protein transcription factor subunit alpha |
| | 66 | 85 | ADSS | adenylosuccinate synthase |
| | 67 | 85 | DHTKD1 | dehydrogenase E1 and transketolase domain containing 1 |
| | 68 | 85 | POU3F2 | POU class 3 homeobox 2 |
| | 69 | 84 | LOXL4 | lysyl oxidase like 4 |
| | 70 | 84 | SLC6A17 | solute carrier family 6 member 17 |
| | 71 | 84 | PER3 | period circadian regulator 3 |
| | 72 | 84 | SLC30A4 | solute carrier family 30 member 4 |

| | | | | |
|---|---|---|---|---|
| | 73 | 84 | ATXN1L | ataxin 1 like |
| | 74 | 84 | FPR3 | formyl peptide receptor 3 |
| | 75 | 84 | GLIPR1 | GLI pathogenesis related 1 |
| | 76 | 84 | GTF3C4 | general transcription factor IIIC subunit 4 |
| | 77 | 84 | PDE6A | phosphodiesterase 6A |
| | 78 | 83 | CPNE5 | copine 5 |
| | 79 | 83 | TTC30A | tetratricopeptide repeat domain 30A |
| | 80 | 83 | SPON1 | spondin 1 |
| | 81 | 83 | DUSP4 | dual specificity phosphatase 4 |
| | 82 | 83 | NUDT5 | nudix hydrolase 5 |
| | 83 | 83 | FRMPD4 | FERM and PDZ domain containing 4 |
| | 84 | 83 | ACTR3 | ARP3 actin related protein 3 homolog |
| | 85 | 82 | TDRKH | tudor and KH domain containing |
| | 86 | 82 | XK | X-linked Kx blood group |
| | 87 | 82 | FGF2 | fibroblast growth factor 2 |
| | 88 | 82 | CHM | CHM, Rab escort protein 1 |

| | | | | |
|---|---|---|---|---|
| | 89 | 82 | CDYL | chromodomain Y like |
| | 90 | 82 | SC5D | sterol-C5-desaturase |
| | 91 | 82 | USF3 | upstream transcription factor family member 3 |
| | 92 | 82 | NKIRAS2 | NFKB inhibitor interacting Ras like 2 |
| | 93 | 81 | GNPNAT1 | glucosamine-phosphate N-acetyltransferase 1 |
| | 94 | 81 | CEP68 | centrosomal protein 68 |
| | 95 | 81 | RIOX1 | ribosomal oxygenase 1 |
| | 96 | 81 | RIPOR2 | RHO family interacting cell polarization regulator 2 |
| | 97 | 81 | MMP20 | matrix metallopeptidase 20 |
| | 98 | 81 | ZNF280D | zinc finger protein 280D |
| | 99 | 80 | SLC35B4 | solute carrier family 35 member B4 |
| | 100 | 80 | SLC30A5 | solute carrier family 30 member 5 |
| | 101 | 80 | STAT2 | signal transducer and activator of transcription 2 |
| | 102 | 80 | FERMT2 | fermitin family member 2 |
| | 103 | 80 | STYX | serine/threonine/tyrosine interacting protein |

| | | | | |
|---|---|---|---|---|
| MD62-5p | 1 | 96 | FAM104A | family with sequence similarity 104 member A |
| | 2 | 94 | GRAP2 | GRB2 related adaptor protein 2 |
| | 3 | 94 | TMEM167B | transmembrane protein 167B |
| | 4 | 93 | NEUROD1 | neuronal differentiation 1 |
| | 5 | 93 | ABCA1 | ATP binding cassette subfamily A member 1 |
| | 6 | 92 | NCOA4 | nuclear receptor coactivator 4 |
| | 7 | 92 | C17orf102 | chromosome 17 open reading frame 102 |
| | 8 | 92 | ACTL6A | actin like 6A |
| | 9 | 91 | PCGF5 | polycomb group ring finger 5 |
| | 10 | 91 | LTV1 | LTV1 ribosome biogenesis factor |
| | 11 | 91 | LIFR | LIF receptor alpha |
| | 12 | 90 | RAB10 | RAB10, member RAS oncogene family |
| | 13 | 90 | WDR20 | WD repeat domain 20 |
| | 14 | 90 | THAP2 | THAP domain containing 2 |
| | 15 | 89 | PARPBP | PARP1 binding protein |
| | 16 | 88 | STK17A | serine/threonine kinase 17a |

| | | | | |
|---|---|---|---|---|
| | 17 | 87 | PAX6 | paired box 6 |
| | 18 | 87 | FNIP1 | folliculin interacting protein 1 |
| | 19 | 87 | SPTLC1 | serine palmitoyltransferase long chain base subunit 1 |
| | 20 | 87 | ERBB4 | erb-b2 receptor tyrosine kinase 4 |
| | 21 | 86 | PRMT3 | protein arginine methyltransferase 3 |
| | 22 | 86 | KIF27 | kinesin family member 27 |
| | 23 | 86 | RNF38 | ring finger protein 38 |
| | 24 | 85 | C5orf34 | chromosome 5 open reading frame 34 |
| | 25 | 85 | GTF2A1 | general transcription factor IIA subunit 1 |
| | 26 | 84 | CKMT1A | creatine kinase, mitochondrial 1A |
| | 27 | 84 | DGKH | diacylglycerol kinase eta |
| | 28 | 84 | PAPSS1 | 3'-phosphoadenosine 5'-phosphosulfate synthase 1 |
| | 29 | 84 | POLR3G | RNA polymerase III subunit G |
| | 30 | 84 | UPF3A | UPF3A, regulator of nonsense mediated mRNA decay |
| | 31 | 84 | CRADD | CASP2 and RIPK1 domain containing adaptor with death domain |
| | 32 | 83 | TNS3 | tensin 3 |

| | | | | |
|---|---|---|---|---|
| | 33 | 83 | ZCRB1 | zinc finger CCHC-type and RNA binding motif containing 1 |
| | 34 | 83 | MMP16 | matrix metallopeptidase 16 |
| | 35 | 83 | SBSPON | somatomedin B and thrombospondin type 1 domain containing |
| | 36 | 81 | TRIP12 | thyroid hormone receptor interactor 12 |
| | 37 | 81 | SLFN13 | schlafen family member 13 |
| | 38 | 81 | CYFIP2 | cytoplasmic FMR1 interacting protein 2 |
| | 39 | 80 | TMEM206 | transmembrane protein 206 |
| | 40 | 80 | NBEA | neurobeachin |
| MD62-3p | 1 | 98 | GNRHR | gonadotropin releasing hormone receptor |
| | 2 | 98 | HIVEP1 | human immunodeficiency virus type I enhancer binding protein 1 |
| | 3 | 97 | RNF11 | ring finger protein 11 |
| | 4 | 97 | CPEB4 | cytoplasmic polyadenylation element binding protein 4 |
| | 5 | 97 | TBL1XR1 | transducin beta like 1 X-linked receptor 1 |
| | 6 | 96 | NANP | N-acetylneuraminic acid phosphatase |
| | 7 | 96 | PDHX | pyruvate dehydrogenase complex component X |

| | | | | |
|---|---|---|---|---|
| | 8 | 96 | SRGAP2B | SLIT-ROBO Rho GTPase activating protein 2B |
| | 9 | 96 | KLF6 | Kruppel like factor 6 |
| | 10 | 95 | TENM1 | teneurin transmembrane protein 1 |
| | 11 | 94 | NEGR1 | neuronal growth regulator 1 |
| | 12 | 94 | C14orf39 | chromosome 14 open reading frame 39 |
| | 13 | 94 | MCTP1 | multiple C2 and transmembrane domain containing 1 |
| | 14 | 94 | LCORL | ligand dependent nuclear receptor corepressor like |
| | 15 | 94 | ERBB2 | erb-b2 receptor tyrosine kinase 2 |
| | 16 | 94 | GDF10 | growth differentiation factor 10 |
| | 17 | 93 | SMCO1 | single-pass membrane protein with coiled-coil domains 1 |
| | 18 | 93 | PIGC | phosphatidylinositol glycan anchor biosynthesis class C |
| | 19 | 93 | ZNF214 | zinc finger protein 214 |
| | 20 | 92 | PAK4 | p21 (RAC1) activated kinase 4 |
| | 21 | 92 | TNFSF13B | TNF superfamily member 13b |
| | 22 | 92 | SMYD4 | SET and MYND domain containing 4 |
| | 23 | 91 | TMEM168 | transmembrane protein 168 |

| | | | | |
|---|---|---|---|---|
| | 24 | 91 | ZNF529 | zinc finger protein 529 |
| | 25 | 91 | LRP1 | LDL receptor related protein 1 |
| | 26 | 91 | GPR85 | G protein-coupled receptor 85 |
| | 27 | 91 | CCDC126 | coiled-coil domain containing 126 |
| | 28 | 91 | NEBL | nebulette |
| | 29 | 90 | CPE | carboxypeptidase E |
| | 30 | 90 | ARL5A | ADP ribosylation factor like GTPase 5A |
| | 31 | 90 | PMP22 | peripheral myelin protein 22 |
| | 32 | 90 | STEAP4 | STEAP4 metalloreductase |
| | 33 | 89 | TASP1 | taspase 1 |
| | 34 | 89 | NTRK2 | neurotrophic receptor tyrosine kinase 2 |
| | 35 | 89 | PTPRD | protein tyrosine phosphatase, receptor type D |
| | 36 | 89 | HDC | histidine decarboxylase |
| | 37 | 88 | KIAA1217 | KIAA1217 |
| | 38 | 88 | NR4A1 | nuclear receptor subfamily 4 group A member 1 |
| | 39 | 88 | ZNF750 | zinc finger protein 750 |

| | | | | |
|---|---|---|---|---|
| | 40 | 88 | RAB21 | RAB21, member RAS oncogene family |
| | 41 | 88 | ARRDC3 | arrestin domain containing 3 |
| | 42 | 88 | MOB1A | MOB kinase activator 1A |
| | 43 | 88 | C2CD5 | C2 calcium dependent domain containing 5 |
| | 44 | 88 | MMRN1 | multimerin 1 |
| | 45 | 88 | ZFR | zinc finger RNA binding protein |
| | 46 | 88 | CYYR1 | cysteine and tyrosine rich 1 |
| | 47 | 88 | FLG2 | filaggrin family member 2 |
| | 48 | 88 | IPO9 | importin 9 |
| | 49 | 87 | ROR1 | receptor tyrosine kinase like orphan receptor 1 |
| | 50 | 87 | FNIP1 | folliculin interacting protein 1 |
| | 51 | 87 | ANGPTL6 | angiopoietin like 6 |
| | 52 | 87 | HS3ST3B1 | heparan sulfate-glucosamine 3-sulfotransferase 3B1 |
| | 53 | 87 | C1orf43 | chromosome 1 open reading frame 43 |
| | 54 | 87 | POF1B | POF1B, actin binding protein |
| | 55 | 87 | KIAA0355 | KIAA0355 |

| | | | | |
|---|---|---|---|---|
| | 56 | 87 | USP54 | ubiquitin specific peptidase 54 |
| | 57 | 86 | JUP | junction plakoglobin |
| | 58 | 86 | PRR18 | proline rich 18 |
| | 59 | 86 | VPS50 | VPS50, EARP/GARPII complex subunit |
| | 60 | 86 | ARPP19 | cAMP regulated phosphoprotein 19 |
| | 61 | 86 | CALHM5 | calcium homeostasis modulator family member 5 |
| | 62 | 86 | PLPP4 | phospholipid phosphatase 4 |
| | 63 | 86 | PRMT2 | protein arginine methyltransferase 2 |
| | 64 | 85 | NSL1 | NSL1, MIS12 kinetochore complex component |
| | 65 | 85 | HPS1 | HPS1, biogenesis of lysosomal organelles complex 3 subunit 1 |
| | 66 | 85 | MBL2 | mannose binding lectin 2 |
| | 67 | 85 | GAB1 | GRB2 associated binding protein 1 |
| | 68 | 85 | LOC100144595 | uncharacterized LOC100144595 |
| | 69 | 85 | URI1 | URI1, prefoldin like chaperone |
| | 70 | 85 | NUFIP1 | nuclear FMR1 interacting protein 1 |
| | 71 | 85 | TIGD6 | tigger transposable element derived 6 |

| | | | | |
|---|---|---|---|---|
| | 72 | 84 | UTS2 | urotensin 2 |
| | 73 | 84 | PRG4 | proteoglycan 4 |
| | 74 | 84 | SLC9B1 | solute carrier family 9 member B1 |
| | 75 | 84 | ZNF365 | zinc finger protein 365 |
| | 76 | 84 | EARS2 | glutamyl-tRNA synthetase 2, mitochondrial |
| | 77 | 84 | FOXN3 | forkhead box N3 |
| | 78 | 84 | SEL1L | SEL1L, ERAD E3 ligase adaptor subunit |
| | 79 | 84 | MKRN1 | makorin ring finger protein 1 |
| | 80 | 83 | EIF3A | eukaryotic translation initiation factor 3 subunit A |
| | 81 | 83 | DOCK4 | dedicator of cytokinesis 4 |
| | 82 | 83 | CDKN1B | cyclin dependent kinase inhibitor 1B |
| | 83 | 83 | ATP11B | ATPase phospholipid transporting 11B (putative) |
| | 84 | 83 | RASSF9 | Ras association domain family member 9 |
| | 85 | 83 | SRGAP2C | SLIT-ROBO Rho GTPase activating protein 2C |
| | 86 | 83 | CLDN11 | claudin 11 |
| | 87 | 83 | GREB1 | growth regulating estrogen receptor binding 1 |

| | | | | |
|---|---|---|---|---|
| | 88 | 83 | ANKLE2 | ankyrin repeat and LEM domain containing 2 |
| | 89 | 82 | C11orf87 | chromosome 11 open reading frame 87 |
| | 90 | 82 | UBAP1 | ubiquitin associated protein 1 |
| | 91 | 82 | IFT88 | intraflagellar transport 88 |
| | 92 | 82 | ATP9B | ATPase phospholipid transporting 9B (putative) |
| | 93 | 82 | JADE3 | jade family PHD finger 3 |
| | 94 | 82 | TRMT9B | tRNA methyltransferase 9B (putative) |
| | 95 | 82 | TMEM132B | transmembrane protein 132B |
| | 96 | 82 | MAP1B | microtubule associated protein 1B |
| | 97 | 81 | CNTNAP2 | contactin associated protein like 2 |
| | 98 | 81 | SESTD1 | SEC14 and spectrin domain containing 1 |
| | 99 | 81 | ITPRID1 | ITPR interacting domain containing 1 |
| | 100 | 81 | STOX1 | storkhead box 1 |
| | 101 | 81 | NRXN1 | neurexin 1 |
| | 102 | 81 | SLC16A6 | solute carrier family 16 member 6 |
| | 103 | 81 | RNF39 | ring finger protein 39 |

| | | | | |
|---|---|---|---|---|
| | 104 | 81 | TFAP2C | transcription factor AP-2 gamma |
| | 105 | 80 | MAPK1 | mitogen-activated protein kinase 1 |
| | 106 | 80 | FAM13C | family with sequence similarity 13 member C |
| | 107 | 80 | PCNX3 | pecanex 3 |
| | 108 | 80 | PDE4D | phosphodiesterase 4D |
| | 109 | 80 | MTF2 | metal response element binding transcription factor 2 |
| | 110 | 80 | ATXN1 | ataxin 1 |
| | 111 | 80 | REL | REL proto-oncogene, NF-kB subunit |
| | 112 | 80 | SCOC | short coiled-coil protein |
| | 113 | 80 | ANKRD1 | ankyrin repeat domain 1 |
| MR5-5p | 1 | 98 | HNRNPK | heterogeneous nuclear ribonucleoprotein K |
| | 2 | 97 | MIB1 | mindbomb E3 ubiquitin protein ligase 1 |
| | 3 | 95 | PCF11 | PCF11, cleavage and polyadenylation factor subunit |
| | 4 | 95 | SMAD7 | SMAD family member 7 |
| | 5 | 95 | RPS6KA3 | ribosomal protein S6 kinase A3 |

| | | | | |
|---|---|---|---|---|
| | 6 | 95 | PTPN4 | protein tyrosine phosphatase, non-receptor type 4 |
| | 7 | 95 | C15orf41 | chromosome 15 open reading frame 41 |
| | 8 | 94 | FERMT3 | fermitin family member 3 |
| | 9 | 94 | ZNF891 | zinc finger protein 891 |
| | 10 | 94 | SECISBP2L | SECIS binding protein 2 like |
| | 11 | 94 | SOAT1 | sterol O-acyltransferase 1 |
| | 12 | 93 | VDAC1 | voltage dependent anion channel 1 |
| | 13 | 93 | ASCC1 | activating signal cointegrator 1 complex subunit 1 |
| | 14 | 93 | BRINP1 | BMP/retinoic acid inducible neural specific 1 |
| | 15 | 93 | SEPTIN7 | septin 7 |
| | 16 | 93 | TAF5L | TATA-box binding protein associated factor 5 like |
| | 17 | 93 | MMP9 | matrix metallopeptidase 9 |
| | 18 | 93 | STRN3 | striatin 3 |
| | 19 | 93 | ALKBH1 | alkB homolog 1, histone H2A dioxygenase |
| | 20 | 92 | PDS5B | PDS5 cohesin associated factor B |
| | 21 | 92 | PRKACB | protein kinase cAMP-activated catalytic subunit beta |

| | | | | |
|---|---|---|---|---|
| | 22 | 92 | KDM7A | lysine demethylase 7A |
| | 23 | 92 | GASK1A | golgi associated kinase 1A |
| | 24 | 92 | SNX15 | sorting nexin 15 |
| | 25 | 91 | DICER1 | dicer 1, ribonuclease III |
| | 26 | 91 | SMIM13 | small integral membrane protein 13 |
| | 27 | 91 | MXI1 | MAX interactor 1, dimerization protein |
| | 28 | 91 | IKZF2 | IKAROS family zinc finger 2 |
| | 29 | 91 | ZNF670 | zinc finger protein 670 |
| | 30 | 91 | ODR4 | odr-4 GPCR localization factor homolog |
| | 31 | 91 | TGS1 | trimethylguanosine synthase 1 |
| | 32 | 90 | AR | androgen receptor |
| | 33 | 90 | PPP2CA | protein phosphatase 2 catalytic subunit alpha |
| | 34 | 90 | GK5 | glycerol kinase 5 |
| | 35 | 90 | SGMS1 | sphingomyelin synthase 1 |
| | 36 | 90 | LIN7C | lin-7 homolog C, crumbs cell polarity complex component |
| | 37 | 90 | DENND1A | DENN domain containing 1A |

| | | | | |
|---|---|---|---|---|
| | 38 | 90 | CUL3 | cullin 3 |
| | 39 | 90 | C1orf94 | chromosome 1 open reading frame 94 |
| | 40 | 90 | ITCH | itchy E3 ubiquitin protein ligase |
| | 41 | 89 | ST8SIA2 | ST8 alpha-N-acetyl-neuraminide alpha-2,8-sialyltransferase 2 |
| | 42 | 89 | TBR1 | T-box, brain 1 |
| | 43 | 89 | MAGEA4 | MAGE family member A4 |
| | 44 | 89 | IFIT5 | interferon induced protein with tetratricopeptide repeats 5 |
| | 45 | 89 | HOXD8 | homeobox D8 |
| | 46 | 89 | KCNK9 | potassium two pore domain channel subfamily K member 9 |
| | 47 | 89 | HMOX1 | heme oxygenase 1 |
| | 48 | 89 | FAM243A | family with sequence similarity 243 member A |
| | 49 | 89 | FBXO33 | F-box protein 33 |
| | 50 | 88 | ARPP21 | cAMP regulated phosphoprotein 21 |
| | 51 | 88 | RFK | riboflavin kinase |
| | 52 | 88 | MTF2 | metal response element binding transcription factor 2 |
| | 53 | 88 | CAMTA1 | calmodulin binding transcription activator 1 |

| | | | | |
|---|---|---|---|---|
| | 54 | 88 | BEND4 | BEN domain containing 4 |
| | 55 | 88 | ATP6V1B2 | ATPase H+ transporting V1 subunit B2 |
| | 56 | 88 | SERPINH1 | serpin family H member 1 |
| | 57 | 88 | SLC39A10 | solute carrier family 39 member 10 |
| | 58 | 87 | HSPA2 | heat shock protein family A (Hsp70) member 2 |
| | 59 | 87 | DISC1 | DISC1 scaffold protein |
| | 60 | 87 | STX7 | syntaxin 7 |
| | 61 | 87 | CWC15 | CWC15 spliceosome associated protein homolog |
| | 62 | 87 | ARHGAP30 | Rho GTPase activating protein 30 |
| | 63 | 87 | FAM19A4 | family with sequence similarity 19 member A4, C-C motif chemokine like |
| | 64 | 87 | DAZL | deleted in azoospermia like |
| | 65 | 87 | MATN4 | matrilin 4 |
| | 66 | 86 | IRX2 | iroquois homeobox 2 |
| | 67 | 86 | ZNF217 | zinc finger protein 217 |
| | 68 | 86 | MINDY2 | MINDY lysine 48 deubiquitinase 2 |
| | 69 | 86 | CHRNA5 | cholinergic receptor nicotinic alpha 5 subunit |

| | | | | |
|---|---|---|---|---|
| | 70 | 86 | LRRC47 | leucine rich repeat containing 47 |
| | 71 | 86 | BTG3 | BTG anti-proliferation factor 3 |
| | 72 | 86 | LTA4H | leukotriene A4 hydrolase |
| | 73 | 86 | NAPB | NSF attachment protein beta |
| | 74 | 86 | ATAD2 | ATPase family, AAA domain containing 2 |
| | 75 | 85 | PDIA6 | protein disulfide isomerase family A member 6 |
| | 76 | 85 | RUFY3 | RUN and FYVE domain containing 3 |
| | 77 | 85 | PCDH8 | protocadherin 8 |
| | 78 | 85 | CCDC177 | coiled-coil domain containing 177 |
| | 79 | 85 | ZNF521 | zinc finger protein 521 |
| | 80 | 85 | RAB21 | RAB21, member RAS oncogene family |
| | 81 | 85 | PRICKLE1 | prickle planar cell polarity protein 1 |
| | 82 | 85 | FBXO9 | F-box protein 9 |
| | 83 | 85 | PGM2L1 | phosphoglucomutase 2 like 1 |
| | 84 | 84 | ZFP42 | ZFP42 zinc finger protein |
| | 85 | 84 | NACA | nascent polypeptide associated complex subunit alpha |

| | | | | |
|---|---|---|---|---|
| | 86 | 84 | C2CD6 | C2 calcium dependent domain containing 6 |
| | 87 | 84 | CBX8 | chromobox 8 |
| | 88 | 84 | SAXO1 | stabilizer of axonemal microtubules 1 |
| | 89 | 84 | RTKN2 | rhotekin 2 |
| | 90 | 84 | SLCO5A1 | solute carrier organic anion transporter family member 5A1 |
| | 91 | 84 | USP3 | ubiquitin specific peptidase 3 |
| | 92 | 84 | VSX1 | visual system homeobox 1 |
| | 93 | 84 | TSPAN3 | tetraspanin 3 |
| | 94 | 84 | CNTN3 | contactin 3 |
| | 95 | 84 | PPP6C | protein phosphatase 6 catalytic subunit |
| | 96 | 84 | SUCLA2 | succinate-CoA ligase ADP-forming beta subunit |
| | 97 | 84 | ARFGEF1 | ADP ribosylation factor guanine nucleotide exchange factor 1 |
| | 98 | 84 | TRMT13 | tRNA methyltransferase 13 homolog |
| | 99 | 83 | TOR1AIP2 | torsin 1A interacting protein 2 |
| | 100 | 83 | ABT1 | activator of basal transcription 1 |
| | 101 | 83 | JARID2 | jumonji and AT-rich interaction domain containing 2 |

| | | | | |
|---|---|---|---|---|
| | 102 | 83 | ZFP36L1 | ZFP36 ring finger protein like 1 |
| | 103 | 83 | ETS1 | ETS proto-oncogene 1, transcription factor |
| | 104 | 83 | ENPP3 | ectonucleotide pyrophosphatase/phosphodiesterase 3 |
| | 105 | 83 | ABCC9 | ATP binding cassette subfamily C member 9 |
| | 106 | 83 | CHD4 | chromodomain helicase DNA binding protein 4 |
| | 107 | 83 | SEC22B | SEC22 homolog B, vesicle trafficking protein (gene/pseudogene) |
| | 108 | 83 | TMTC1 | transmembrane and tetratricopeptide repeat containing 1 |
| | 109 | 83 | B3GALNT1 | beta-1,3-N-acetylgalactosaminyltransferase 1 (globoside blood group) |
| | 110 | 82 | RBM26 | RNA binding motif protein 26 |
| | 111 | 82 | ELAVL2 | ELAV like RNA binding protein 2 |
| | 112 | 82 | ALG13 | ALG13, UDP-N-acetylglucosaminyltransferase subunit |
| | 113 | 82 | ENOSF1 | enolase superfamily member 1 |
| | 114 | 82 | HTR1D | 5-hydroxytryptamine receptor 1D |
| | 115 | 82 | YPEL2 | yippee like 2 |
| | 116 | 82 | CLMN | calmin |
| | 117 | 82 | HIPK1 | homeodomain interacting protein kinase 1 |

| | | | | |
|---|---|---|---|---|
| | 118 | 81 | MATN3 | matrilin 3 |
| | 119 | 81 | ANKRD44 | ankyrin repeat domain 44 |
| | 120 | 81 | SENP8 | SUMO peptidase family member, NEDD8 specific |
| | 121 | 81 | BTN2A1 | butyrophilin subfamily 2 member A1 |
| | 122 | 81 | RHOQ | ras homolog family member Q |
| | 123 | 81 | YPEL5 | yippee like 5 |
| | 124 | 81 | ZSCAN12 | zinc finger and SCAN domain containing 12 |
| | 125 | 81 | RANBP17 | RAN binding protein 17 |
| | 126 | 81 | MGST1 | microsomal glutathione S-transferase 1 |
| | 127 | 81 | PURA | purine rich element binding protein A |
| | 128 | 81 | ZNF12 | zinc finger protein 12 |
| | 129 | 81 | SERBP1 | SERPINE1 mRNA binding protein 1 |
| | 130 | 81 | MTHFS | methenyltetrahydrofolate synthetase |
| | 131 | 81 | ST20-MTHFS | ST20-MTHFS readthrough |
| | 132 | 80 | NAMPT | nicotinamide phosphoribosyltransferase |
| | 133 | 80 | KAT6A | lysine acetyltransferase 6A |

| | | | | |
|---|---|---|---|---|
| | 134 | 80 | CNOT1 | CCR4-NOT transcription complex subunit 1 |
| | 135 | 80 | DCLK2 | doublecortin like kinase 2 |
| | 136 | 80 | ITPRID2 | ITPR interacting domain containing 2 |
| | 137 | 80 | XYLT1 | xylosyltransferase 1 |
| | 138 | 80 | THBS2 | thrombospondin 2 |
| | 139 | 80 | AFF2 | AF4/FMR2 family member 2 |
| | 140 | 80 | IREB2 | iron responsive element binding protein 2 |
| | 141 | 80 | ARPIN | actin related protein 2/3 complex inhibitor |
| | 142 | 80 | PIK3CB | phosphatidylinositol-4,5-bisphosphate 3-kinase catalytic subunit beta |
| | 143 | 80 | PRKAR2B | protein kinase cAMP-dependent type II regulatory subunit beta |
| | 144 | 80 | FSHB | follicle stimulating hormone subunit beta |
| | 145 | 80 | UBE2K | ubiquitin conjugating enzyme E2 K |
| MR3-3p | 1 | 100 | LIG4 | DNA ligase 4 |
| | 2 | 99 | PARS2 | prolyl-tRNA synthetase 2, mitochondrial |
| | 3 | 99 | HAPLN1 | hyaluronan and proteoglycan link protein 1 |

| | | | | |
|---|---|---|---|---|
| | 4 | 99 | CNTNAP2 | contactin associated protein like 2 |
| | 5 | 97 | OPA1 | OPA1, mitochondrial dynamin like GTPase |
| | 6 | 97 | BIRC6 | baculoviral IAP repeat containing 6 |
| | 7 | 97 | PGM3 | phosphoglucomutase 3 |
| | 8 | 96 | MAFF | MAF bZIP transcription factor F |
| | 9 | 96 | ZBTB34 | zinc finger and BTB domain containing 34 |
| | 10 | 96 | TECTB | tectorin beta |
| | 11 | 96 | SEMA6D | semaphorin 6D |
| | 12 | 96 | SUCNR1 | succinate receptor 1 |
| | 13 | 96 | PAWR | pro-apoptotic WT1 regulator |
| | 14 | 96 | YBX1 | Y-box binding protein 1 |
| | 15 | 96 | SLITRK2 | SLIT and NTRK like family member 2 |
| | 16 | 96 | MAP3K2 | mitogen-activated protein kinase kinase kinase 2 |
| | 17 | 96 | A1CF | APOBEC1 complementation factor |
| | 18 | 95 | MCCC2 | methylcrotonoyl-CoA carboxylase 2 |
| | 19 | 95 | CBLN2 | cerebellin 2 precursor |

| | | | | |
|---|---|---|---|---|
| | 20 | 95 | C1orf162 | chromosome 1 open reading frame 162 |
| | 21 | 95 | RGS13 | regulator of G protein signaling 13 |
| | 22 | 95 | ZFP42 | ZFP42 zinc finger protein |
| | 23 | 95 | MLLT10 | MLLT10, histone lysine methyltransferase DOT1L cofactor |
| | 24 | 94 | BRD1 | bromodomain containing 1 |
| | 25 | 94 | GPX8 | glutathione peroxidase 8 (putative) |
| | 26 | 94 | DNER | delta/notch like EGF repeat containing |
| | 27 | 94 | CMTR2 | cap methyltransferase 2 |
| | 28 | 94 | CNKSR1 | connector enhancer of kinase suppressor of Ras 1 |
| | 29 | 94 | HHIP | hedgehog interacting protein |
| | 30 | 94 | SLC2A4 | solute carrier family 2 member 4 |
| | 31 | 94 | PPP4R2 | protein phosphatase 4 regulatory subunit 2 |
| | 32 | 94 | AP1G1 | adaptor related protein complex 1 subunit gamma 1 |
| | 33 | 93 | MAP1B | microtubule associated protein 1B |
| | 34 | 93 | STK24 | serine/threonine kinase 24 |
| | 35 | 93 | RFX6 | regulatory factor X6 |

| | | | | |
|---|---|---|---|---|
| | 36 | 93 | PPP5C | protein phosphatase 5 catalytic subunit |
| | 37 | 93 | IPO7 | importin 7 |
| | 38 | 93 | C2CD2L | C2CD2 like |
| | 39 | 93 | UBFD1 | ubiquitin family domain containing 1 |
| | 40 | 92 | LILRB4 | leukocyte immunoglobulin like receptor B4 |
| | 41 | 92 | TIAL1 | TIA1 cytotoxic granule associated RNA binding protein like 1 |
| | 42 | 92 | SMC5 | structural maintenance of chromosomes 5 |
| | 43 | 92 | GOLT1B | golgi transport 1B |
| | 44 | 92 | FER | FER tyrosine kinase |
| | 45 | 92 | CNOT1 | CCR4-NOT transcription complex subunit 1 |
| | 46 | 92 | FSD1L | fibronectin type III and SPRY domain containing 1 like |
| | 47 | 92 | DKC1 | dyskerin pseudouridine synthase 1 |
| | 48 | 92 | PPFIBP1 | PPFIA binding protein 1 |
| | 49 | 92 | C5orf24 | chromosome 5 open reading frame 24 |
| | 50 | 92 | PROX1 | prospero homeobox 1 |
| | 51 | 92 | KCNT2 | potassium sodium-activated channel subfamily T member 2 |

| | | | | |
|---|---|---|---|---|
| | 52 | 92 | FEN1 | flap structure-specific endonuclease 1 |
| | 53 | 92 | PLCXD1 | phosphatidylinositol specific phospholipase C X domain containing 1 |
| | 54 | 91 | SLAMF9 | SLAM family member 9 |
| | 55 | 91 | PKHD1 | PKHD1, fibrocystin/polyductin |
| | 56 | 91 | COBLL1 | cordon-bleu WH2 repeat protein like 1 |
| | 57 | 91 | DRAM2 | DNA damage regulated autophagy modulator 2 |
| | 58 | 91 | PACSIN1 | protein kinase C and casein kinase substrate in neurons 1 |
| | 59 | 91 | HERPUD2 | HERPUD family member 2 |
| | 60 | 91 | SEC62 | SEC62 homolog, preprotein translocation factor |
| | 61 | 91 | UNC80 | unc-80 homolog, NALCN channel complex subunit |
| | 62 | 91 | P2RY1 | purinergic receptor P2Y1 |
| | 63 | 91 | CSF1 | colony stimulating factor 1 |
| | 64 | 91 | NMT2 | N-myristoyltransferase 2 |
| | 65 | 91 | RAI1 | retinoic acid induced 1 |
| | 66 | 91 | BLOC1S6 | biogenesis of lysosomal organelles complex 1 subunit 6 |
| | 67 | 91 | ERCC6L2 | ERCC excision repair 6 like 2 |

| | | | | |
|---|---|---|---|---|
| | 68 | 91 | C11orf96 | chromosome 11 open reading frame 96 |
| | 69 | 90 | PPP1CB | protein phosphatase 1 catalytic subunit beta |
| | 70 | 90 | POU2F1 | POU class 2 homeobox 1 |
| | 71 | 90 | HTR2B | 5-hydroxytryptamine receptor 2B |
| | 72 | 90 | TAB3 | TGF-beta activated kinase 1 (MAP3K7) binding protein 3 |
| | 73 | 90 | FLRT2 | fibronectin leucine rich transmembrane protein 2 |
| | 74 | 90 | B3GALNT1 | beta-1,3-N-acetylgalactosaminyltransferase 1 (globoside blood group) |
| | 75 | 90 | INSM1 | INSM transcriptional repressor 1 |
| | 76 | 89 | ZNF268 | zinc finger protein 268 |
| | 77 | 89 | RNF169 | ring finger protein 169 |
| | 78 | 89 | RRM1 | ribonucleotide reductase catalytic subunit M1 |
| | 79 | 89 | MYO1D | myosin ID |
| | 80 | 89 | NEUROG1 | neurogenin 1 |
| | 81 | 89 | CHRM3 | cholinergic receptor muscarinic 3 |
| | 82 | 89 | CLCA2 | chloride channel accessory 2 |
| | 83 | 89 | CCDC127 | coiled-coil domain containing 127 |

| | | | | |
|---|---|---|---|---|
| | 84 | 89 | HNF1A | HNF1 homeobox A |
| | 85 | 89 | NEFL | neurofilament light |
| | 86 | 89 | RIMS2 | regulating synaptic membrane exocytosis 2 |
| | 87 | 89 | CPEB2 | cytoplasmic polyadenylation element binding protein 2 |
| | 88 | 89 | CABP5 | calcium binding protein 5 |
| | 89 | 89 | PTER | phosphotriesterase related |
| | 90 | 89 | CCDC170 | coiled-coil domain containing 170 |
| | 91 | 89 | RIC3 | RIC3 acetylcholine receptor chaperone |
| | 92 | 89 | KCNQ3 | potassium voltage-gated channel subfamily Q member 3 |
| | 93 | 89 | SEL1L3 | SEL1L family member 3 |
| | 94 | 89 | KCNN3 | potassium calcium-activated channel subfamily N member 3 |
| | 95 | 88 | INHBC | inhibin subunit beta C |
| | 96 | 88 | ONECUT2 | one cut homeobox 2 |
| | 97 | 88 | SP100 | SP100 nuclear antigen |
| | 98 | 88 | MMP1 | matrix metallopeptidase 1 |
| | 99 | 88 | SLC5A1 | solute carrier family 5 member 1 |

| | | | | |
|---|---|---|---|---|
| | 100 | 88 | ZBTB8B | zinc finger and BTB domain containing 8B |
| | 101 | 88 | FOXJ3 | forkhead box J3 |
| | 102 | 88 | FAM135A | family with sequence similarity 135 member A |
| | 103 | 88 | CEP57 | centrosomal protein 57 |
| | 104 | 88 | ZFPM2 | zinc finger protein, FOG family member 2 |
| | 105 | 88 | CLASRP | CLK4 associating serine/arginine rich protein |
| | 106 | 88 | EDIL3 | EGF like repeats and discoidin domains 3 |
| | 107 | 87 | ZNF251 | zinc finger protein 251 |
| | 108 | 87 | CHCHD2 | coiled-coil-helix-coiled-coil-helix domain containing 2 |
| | 109 | 87 | ZNF567 | zinc finger protein 567 |
| | 110 | 87 | MIA2 | MIA SH3 domain ER export factor 2 |
| | 111 | 87 | TLL1 | tolloid like 1 |
| | 112 | 87 | SLC14A1 | solute carrier family 14 member 1 (Kidd blood group) |
| | 113 | 87 | KIF3A | kinesin family member 3A |
| | 114 | 87 | TMEM231 | transmembrane protein 231 |
| | 115 | 87 | NPPB | natriuretic peptide B |

| | | | | |
|---|---|---|---|---|
| | 116 | 87 | WBP1L | WW domain binding protein 1 like |
| | 117 | 87 | LIAS | lipoic acid synthetase |
| | 118 | 87 | B4GALT6 | beta-1,4-galactosyltransferase 6 |
| | 119 | 87 | NEK4 | NIMA related kinase 4 |
| | 120 | 87 | MARCHF3 | membrane associated ring-CH-type finger 3 |
| | 121 | 87 | ARHGAP5 | Rho GTPase activating protein 5 |
| | 122 | 87 | GKN2 | gastrokine 2 |
| | 123 | 87 | WEE1 | WEE1 G2 checkpoint kinase |
| | 124 | 87 | STXBP4 | syntaxin binding protein 4 |
| | 125 | 86 | MBL2 | mannose binding lectin 2 |
| | 126 | 86 | MYOCD | myocardin |
| | 127 | 86 | KIRREL2 | kirre like nephrin family adhesion molecule 2 |
| | 128 | 86 | PSMD5 | proteasome 26S subunit, non-ATPase 5 |
| | 129 | 86 | SCN1A | sodium voltage-gated channel alpha subunit 1 |
| | 130 | 86 | PDS5A | PDS5 cohesin associated factor A |
| | 131 | 86 | SMC2 | structural maintenance of chromosomes 2 |

| | | | | |
|---|---|---|---|---|
| | 132 | 86 | PEX2 | peroxisomal biogenesis factor 2 |
| | 133 | 86 | TTC13 | tetratricopeptide repeat domain 13 |
| | 134 | 86 | ELF5 | E74 like ETS transcription factor 5 |
| | 135 | 86 | CDH19 | cadherin 19 |
| | 136 | 86 | SLC35G3 | solute carrier family 35 member G3 |
| | 137 | 86 | DDTL | D-dopachrome tautomerase like |
| | 138 | 85 | TMEM209 | transmembrane protein 209 |
| | 139 | 85 | DHX57 | DExH-box helicase 57 |
| | 140 | 85 | LRRC8C | leucine rich repeat containing 8 VRAC subunit C |
| | 141 | 85 | FZD2 | frizzled class receptor 2 |
| | 142 | 85 | MCEMP1 | mast cell expressed membrane protein 1 |
| | 143 | 85 | CALHM4 | calcium homeostasis modulator family member 4 |
| | 144 | 85 | USP25 | ubiquitin specific peptidase 25 |
| | 145 | 85 | ARCN1 | archain 1 |
| | 146 | 85 | WBP4 | WW domain binding protein 4 |
| | 147 | 85 | ALG10B | ALG10B, alpha-1,2-glucosyltransferase |

| | | | | |
|---|---|---|---|---|
| | 148 | 85 | CDT1 | chromatin licensing and DNA replication factor 1 |
| | 149 | 85 | EBF3 | EBF transcription factor 3 |
| | 150 | 85 | ANKDD1B | ankyrin repeat and death domain containing 1B |
| | 151 | 85 | GPD2 | glycerol-3-phosphate dehydrogenase 2 |
| | 152 | 84 | PURA | purine rich element binding protein A |
| | 153 | 84 | OGFOD3 | 2-oxoglutarate and iron dependent oxygenase domain containing 3 |
| | 154 | 84 | SOX5 | SRY-box 5 |
| | 155 | 84 | CD93 | CD93 molecule |
| | 156 | 84 | LONP1 | lon peptidase 1, mitochondrial |
| | 157 | 84 | C9orf85 | chromosome 9 open reading frame 85 |
| | 158 | 84 | TPR | translocated promoter region, nuclear basket protein |
| | 159 | 84 | STON2 | stonin 2 |
| | 160 | 84 | SLC8B1 | solute carrier family 8 member B1 |
| | 161 | 84 | EID1 | EP300 interacting inhibitor of differentiation 1 |
| | 162 | 84 | NR5A2 | nuclear receptor subfamily 5 group A member 2 |
| | 163 | 84 | DPP10 | dipeptidyl peptidase like 10 |

| | | | | |
|---|---|---|---|---|
| | 164 | 84 | DCN | decorin |
| | 165 | 84 | C1QTNF3 | C1q and TNF related 3 |
| | 166 | 84 | FGD4 | FYVE, RhoGEF and PH domain containing 4 |
| | 167 | 84 | PDK3 | pyruvate dehydrogenase kinase 3 |
| | 168 | 83 | IGSF10 | immunoglobulin superfamily member 10 |
| | 169 | 83 | C5orf51 | chromosome 5 open reading frame 51 |
| | 170 | 83 | ZDHHC21 | zinc finger DHHC-type containing 21 |
| | 171 | 83 | SSH2 | slingshot protein phosphatase 2 |
| | 172 | 83 | UTP3 | UTP3, small subunit processome component |
| | 173 | 83 | THSD4 | thrombospondin type 1 domain containing 4 |
| | 174 | 83 | SPATA6L | spermatogenesis associated 6 like |
| | 175 | 83 | CNKSR2 | connector enhancer of kinase suppressor of Ras 2 |
| | 176 | 83 | IL19 | interleukin 19 |
| | 177 | 83 | RIPK2 | receptor interacting serine/threonine kinase 2 |
| | 178 | 83 | KBTBD6 | kelch repeat and BTB domain containing 6 |
| | 179 | 83 | ATP2B4 | ATPase plasma membrane Ca2+ transporting 4 |

| | | | | |
|---|---|---|---|---|
| | 180 | 83 | PGRMC2 | progesterone receptor membrane component 2 |
| | 181 | 83 | DCBLD2 | discoidin, CUB and LCCL domain containing 2 |
| | 182 | 83 | RNF217 | ring finger protein 217 |
| | 183 | 83 | MAPK15 | mitogen-activated protein kinase 15 |
| | 184 | 82 | ITGA10 | integrin subunit alpha 10 |
| | 185 | 82 | ANOS1 | anosmin 1 |
| | 186 | 82 | TMEM178B | transmembrane protein 178B |
| | 187 | 82 | BICD2 | BICD cargo adaptor 2 |
| | 188 | 82 | GRIA3 | glutamate ionotropic receptor AMPA type subunit 3 |
| | 189 | 82 | RBM27 | RNA binding motif protein 27 |
| | 190 | 82 | FAM13C | family with sequence similarity 13 member C |
| | 191 | 82 | LACTB | lactamase beta |
| | 192 | 82 | CEBPG | CCAAT enhancer binding protein gamma |
| | 193 | 82 | ATP8A1 | ATPase phospholipid transporting 8A1 |
| | 194 | 82 | POF1B | POF1B, actin binding protein |
| | 195 | 82 | GNG3 | G protein subunit gamma 3 |

| | | | | |
|---|---|---|---|---|
| | 196 | 82 | C1orf115 | chromosome 1 open reading frame 115 |
| | 197 | 82 | MKNK2 | MAP kinase interacting serine/threonine kinase 2 |
| | 198 | 82 | KRTAP9-8 | keratin associated protein 9-8 |
| | 199 | 82 | KRTAP9-2 | keratin associated protein 9-2 |
| | 200 | 82 | DISC1 | DISC1 scaffold protein |
| | 201 | 82 | YWHAG | tyrosine 3-monooxygenase/tryptophan 5-monooxygenase activation protein gamma |
| | 202 | 82 | CDYL2 | chromodomain Y like 2 |
| | 203 | 82 | C12orf65 | chromosome 12 open reading frame 65 |
| | 204 | 81 | MORF4L1 | mortality factor 4 like 1 |
| | 205 | 81 | DOK1 | docking protein 1 |
| | 206 | 81 | ETF1 | eukaryotic translation termination factor 1 |
| | 207 | 81 | L3HYPDH | trans-L-3-hydroxyproline dehydratase |
| | 208 | 81 | UBE2D3 | ubiquitin conjugating enzyme E2 D3 |
| | 209 | 81 | IL36B | interleukin 36 beta |
| | 210 | 81 | ASPH | aspartate beta-hydroxylase |
| | 211 | 81 | TKTL2 | transketolase like 2 |

| | | | | |
|---|---|---|---|---|
| | 212 | 81 | PAF1 | PAF1 homolog, Paf1/RNA polymerase II complex component |
| | 213 | 81 | CBFB | core-binding factor subunit beta |
| | 214 | 81 | RNF38 | ring finger protein 38 |
| | 215 | 81 | FAP | fibroblast activation protein alpha |
| | 216 | 81 | PGK1 | phosphoglycerate kinase 1 |
| | 217 | 81 | SAR1A | secretion associated Ras related GTPase 1A |
| | 218 | 81 | ADAMTS18 | ADAM metallopeptidase with thrombospondin type 1 motif 18 |
| | 219 | 81 | ARNTL | aryl hydrocarbon receptor nuclear translocator like |
| | 220 | 81 | ENTPD1 | ectonucleoside triphosphate diphosphohydrolase 1 |
| | 221 | 80 | TET2 | tet methylcytosine dioxygenase 2 |
| | 222 | 80 | RNF139 | ring finger protein 139 |
| | 223 | 80 | KIAA1324L | KIAA1324 like |
| | 224 | 80 | PDLIM3 | PDZ and LIM domain 3 |
| | 225 | 80 | NBEA | neurobeachin |
| | 226 | 80 | PHACTR2 | phosphatase and actin regulator 2 |
| | 227 | 80 | CERKL | ceramide kinase like |

| | | | | |
|---|---|---|---|---|
| | 228 | 80 | ZNF697 | zinc finger protein 697 |
| | 229 | 80 | ACSL1 | acyl-CoA synthetase long chain family member 1 |
| | 230 | 80 | RWDD3 | RWD domain containing 3 |
| | 231 | 80 | CPNE8 | copine 8 |
| | 232 | 80 | RAB3GAP2 | RAB3 GTPase activating non-catalytic protein subunit 2 |
| | 233 | 80 | FAM111A | family with sequence similarity 111 member A |
| | 234 | 80 | LRRC10 | leucine rich repeat containing 10 |
| | 235 | 80 | CLPSL2 | colipase like 2 |
| | 236 | 80 | USP6 | ubiquitin specific peptidase 6 |
| | 237 | 80 | G6PC | glucose-6-phosphatase catalytic subunit |
| | 238 | 80 | ZRANB3 | zinc finger RANBP2-type containing 3 |
| | 239 | 80 | FNDC3B | fibronectin type III domain containing 3B |
| | 240 | 80 | RAG2 | recombination activating 2 |

**Sheet_5**

**List of target mRNAs downregulated in various transcriptomic studies.**

| Xiong *et. al.* | | Blanco-Melo *et. al.* | | | Zhuo Zhou *et. al.* |
|---|---|---|---|---|---|
| *Emerg Microbes Infect.* 2020 | | *Cell*, 2020 | | | *Cell Host Microbe*, 2020 |
| COVID-19 patients-BALF | COVID-19 patients-PBL | COVID-19 patients-Post-mortem Lung | SARS-Cov-2 infected A549 | SARS-Cov-2 infected A549-ACE-2 | COVID-19 patients-BALF |
| 325 | 316 | 2408 | 77 | 96 | 739 |
| 15 | 19 | 126 | 6 | 4 | 56 |
| ACER3 | ANGPTL6 | ABT1 | CBLN2 | ASB9 | ABCA1 |
| ARHGAP30 | CCDC80 | ACTL6A | CNTNAP2 | FLRT2 | AHNAK |
| C2CD2L | CD247 | AHNAK | FEN1 | SERPINH1 | ANKRD44 |
| ITGAM | DRAXIN | ANKDD1A | NR4A1 | SLC40A1 | ARHGEF7 |
| LRP1 | ENPP3 | AR | OLFML3 | | ATP2B4 |
| MRC1 | G0S2 | ARHGEF7 | PACSIN1 | | ATP9B |
| PCNX3 | IKZF2 | ARNTL | | | BAZ2A |
| PDIA6 | KIF3A | ASAP2 | | | BCL11A |
| PIGK | MYBL1 | ATXN7L3 | | | C20orf194 |

| | | | | | |
|---|---|---|---|---|---|
| SEL1L | PER3 | BNC2 | | | C2CD2L |
| SMAD7 | PLEKHA1 | C20orf194 | | | CHFR |
| SMARCD2 | PRSS23 | CACNA1C | | | CPM |
| TMEM127 | PTCH1 | CEP68 | | | DMXL2 |
| TSPAN3 | SLC30A4 | CHFR | | | DNMT3A |
| USP19 | SPON1 | CHL1 | | | DOCK3 |
| | TSHZ2 | CLDN11 | | | EIF4ENIF1 |
| | ZBTB20 | CPE | | | GPNMB |
| | ZNF365 | DDHD2 | | | HERC1 |
| | ZNF831 | DHX57 | | | KCNMA1 |
| | | DOK1 | | | KLHL3 |
| | | EGR1 | | | LPCAT2 |
| | | ENOSF1 | | | LRP1 |
| | | ERBB2 | | | LRRC8C |
| | | F2RL3 | | | LTA4H |
| | | FAM135A | | | MKNK1 |
| | | FBXO9 | | | MRC1 |

| | | | | | |
|---|---|---|---|---|---|
| | | FER | | | OLR1 |
| | | FOXJ2 | | | PCNX3 |
| | | FZD4 | | | PIAS2 |
| | | FZD8 | | | POU2F1 |
| | | GJC1 | | | PPHLN1 |
| | | GKN2 | | | PPM1M |
| | | GNL3 | | | PTPRM |
| | | HMOX1 | | | RBM26 |
| | | HSPA2 | | | RELCH |
| | | ID3 | | | SCN2B |
| | | IDH2 | | | SCN8A |
| | | IFT74 | | | SCUBE3 |
| | | INPP5A | | | SEL1L |
| | | ITGA10 | | | SKI |
| | | ITM2C | | | SLC39A10 |
| | | KCNN3 | | | SLC8B1 |
| | | KCNQ3 | | | SNX1 |

| | | | | | |
|---|---|---|---|---|---|
| | | KCTD20 | | | SOAT1 |
| | | KDM3A | | | SSH2 |
| | | KIAA0355 | | | ST3GAL1 |
| | | KLHL3 | | | TFRC |
| | | L3MBTL3 | | | THSD4 |
| | | LEPR | | | TIAM1 |
| | | LONP1 | | | TNS3 |
| | | LRIG2 | | | TRIM66 |
| | | LRP6 | | | TRIO |
| | | LRRC47 | | | TTC28 |
| | | MACROD2 | | | UNC80 |
| | | MAMDC2 | | | VPS13B |
| | | MAPK8 | | | ZNF891 |
| | | MARK1 | | | |
| | | MECP2 | | | |
| | | METTL8 | | | |
| | | MIB1 | | | |

| | | |
|---|---|---|
| | | MLLT10 |
| | | MMP1 |
| | | MRPL48 |
| | | MXI1 |
| | | MYO1D |
| | | NANP |
| | | NEBL |
| | | NEK4 |
| | | NMT2 |
| | | NR4A1 |
| | | NR5A2 |
| | | NSUN4 |
| | | NTRK2 |
| | | OCLN |
| | | PAWR |
| | | PER3 |
| | | PEX2 |

| | | |
|---|---|---|
| | | PIAS2 |
| | | PPM1M |
| | | PPP1R16B |
| | | PPP3R1 |
| | | PRICKLE2 |
| | | PRMT2 |
| | | PRPF39 |
| | | PTPRD |
| | | RAI1 |
| | | RBM26 |
| | | RCN2 |
| | | RDX |
| | | RNF219 |
| | | RTKN2 |
| | | RUFY3 |
| | | SBNO1 |
| | | SEMA6D |

| | | |
|---|---|---|
| | | SEPSECS |
| | | SGK1 |
| | | SGMS1 |
| | | SLC16A6 |
| | | SMAD7 |
| | | SMARCD2 |
| | | SSH2 |
| | | STC1 |
| | | STON2 |
| | | STYX |
| | | TARDBP |
| | | TBX4 |
| | | THAP2 |
| | | THAP5 |
| | | TIAM1 |
| | | TIGD6 |
| | | TLL1 |

| | | |
|---|---|---|
| | | TMEM231<br>TMTC1<br>TRPS1<br>USP19<br>VAPB<br>WEE1<br>XPNPEP3<br>XYLT1<br>ZBTB18<br>ZBTB34<br>ZNF12<br>ZNF521<br>ZNF697<br>ZNF761<br>ZNF83 |

**Sheet_6**

**List of GO term and KEGG pathways enriched for the target mRNAs.**

| GO ID | GO Term | Count | PValue | Genes (Entrez ID) |
|---|---|---|---|---|
| **Biological Processes** | | | | |
| GO:0000122 | Negative regulation of transcription from RNA polymerase II promoter | 77 | 1E-06 | 4208, 7022, 96459, 1761, 4204, 3720, 23512, 7753, 1108, 7707, 10795, 53335, 7490, 27063, 6927, 6929, 128553, 57332, 8325, 23405, 3234, 6497, 54816, 153222, 54623, 5074, 639, 55758, 5078, 79718, 3096, 4848, 10472, 23741, 7175, 8725, 342371, 8452, 9960, 9314, 3642, 23414, 8863, 64750, 1602, 1958, 7227, 2627, 10363, 5080, 1788, 387893, 9678, 23019, 2274, 3399, 23327, 4092, 5828, 4904, 5966, 22823, 2551, 163081, 29966, 5629, 9734, 7764, 5727, 9839, 10499, 7323, 6672, 93649, 26137, 2186, 9338 |
| GO:0045893 | Positive regulation of transcription, DNA-templated | 57 | 1E-05 | 4208, 4040, 4204, 1316, 29028, 5991, 7490, 6927, 6929, 55818, 8322, 2247, 7994, 4760, 55810, 2066, 222546, 79718, 9063, 444, 4603, 55366, 406, 2113, 9314, 5594, 55617, 1958, 8031, 2627, 1749, 9575, 5080, 5015, 2274, 6787, 367, 4090, 9325, 6714, 3275, 56970, 5629, 8609, 1112, 285704, 6667, 5727, 80853, 10743, 2494, 6672, 93649, 2535, 91612, 2186, 10746 |
| GO:0045944 | Positive regulation of transcription from RNA polymerase II promoter | 92 | 2E-05 | 4208, 7022, 4040, 1761, 7707, 10795, 53335, 8767, 29028, 5991, 7490, 27063, 5534, 6927, 6929, 55818, 1054, 59343, 2247, 4762, 8325, 4760, 10135, 3234, 6497, 55810, 3190, 10716, 22807, 54623, 57496, 222546, 79718, 79755, 8028, 3552, 54790, 9063, 4603, 5028, 23261, 6197, 9480, 6938, 80173, 406, 84333, 9314, 2113, 23414, 253738, 5454, 1958, 865, 7227, 1747, 2627, 9575, 5080, 55252, 5015, 367, 8440, 4808, 4092, 4904, 116931, 5966, 659, 22823, 2551, 163081, 29966, 5629, 30061, 2488, 6667, 9839, 10499, 10743, 23764, 23433, 466, 3164, 2494, 4060, 93649, 26747, 1634, 905, 2001, 51142 |
| GO:0006351 | Transcription, DNA-templated | 153 | 4E-04 | 4208, 147686, 221833, 4204, 23512, 7753, 3720, 1108, 6773, 10795, 285349, 1316, 29028, 90987, 5536, 51008, 9753, 389114, 6497, 54816, 10716, 22807, 7748, 126017, 30813, 22887, 204851, 219736, 55758, 79718, 8028, 55193, 79697, 10472, 54796, 342371, 8725, 4601, 406, 6603, 84333, 5451, 253738, 8863, 84295, 403341, 728116, 8031, 7799, 93474, 80264, 9575, 388561, 387893, 144108, 23019, 2274, 1500, 57615, 3222, 4090, 152485, 3399, 10178, 4808, 4092, 9425, 55205, 101060200, 56970, 5629, 1112, 27107, 80853, 5813, 10499, 10933, 6672, 26137, 2186, 7024, 7594, 51535, 53335, 126068, 25925, 5991, 10957, 57531, 168451, 6929, 55818, 128553, 57332, 4762, 7994, 9496, 86, 132625, 153222, 2066, 5074, 639, 2064, 57496, 5078, 5411, 339500, 6310, 55892, 96764, 4848, 9063, 23741, 3642, 23414, 5594, 7572, 55769, |

| | | | | |
|---|---|---|---|---|
| | | | | 10363, 90874, 84458, 5080, 84456, 55252, 57786, 286205, 11176, 7767, 367, 256380, 116931, 9325, 9329, 163081, 2140, 57711, 8609, 9734, 7764, 9839, 7761, 54014, 3164, 2494, 6917, 55571, 91612, 7559, 4666, 905, 51142, 9338 |
| GO:0000278 | Mitotic cell cycle | 10 | 6E-04 | 10298, 57144, 8428, 6787, 8452, 9960, 6240, 55844, 2241, 1788 |
| GO:0006366 | Transcription from RNA polymerase II promoter | 50 | 9E-04 | 4208, 7022, 1761, 865, 1958, 7227, 7753, 7707, 1747, 1749, 2627, 9575, 254251, 5080, 7490, 6927, 1054, 5015, 4760, 2957, 3234, 6660, 29777, 3190, 55810, 6921, 4904, 5966, 659, 2551, 128611, 222546, 3096, 79755, 27097, 4603, 23261, 23764, 6938, 9480, 466, 23435, 6917, 93649, 9314, 2113, 905, 2001, 5454, 253738 |
| GO:0003309 | Type B pancreatic cell differentiation | 5 | 1E-03 | 222546, 3642, 2627, 5991, 5080 |
| GO:0030182 | Neuron differentiation | 15 | 2E-03 | 4208, 204851, 5629, 9734, 1788, 136319, 8322, 4762, 8325, 3399, 2535, 4915, 81618, 5789, 5454 |
| GO:0032869 | Cellular response to insulin stimulus | 13 | 2E-03 | 677, 132864, 6714, 10890, 9734, 6517, 9847, 6667, 2241, 2693, 23433, 7532, 5584 |
| GO:0045765 | Regulation of angiogenesis | 8 | 2E-03 | 3162, 2064, 2013, 2247, 6672, 2113, 23554, 10457 |
| GO:0017148 | Negative regulation of translation | 11 | 2E-03 | 5813, 11236, 10643, 23019, 10644, 10642, 3658, 56853, 56478, 7490, 4848 |
| GO:0031018 | Endocrine pancreas development | 7 | 4E-03 | 6927, 9480, 5078, 222546, 4760, 3642, 5991 |
| GO:0034220 | Ion transmembrane transport | 24 | 4E-03 | 285175, 2562, 133308, 2561, 10345, 6557, 23200, 2892, 2890, 361, 10396, 444, 2897, 493, 254228, 121601, 2558, 23327, 6446, 9635, 10052, 221301, 526, 150159 |
| GO:0050790 | Regulation of catalytic activity | 11 | 6E-03 | 10527, 5523, 26051, 57332, 22999, 367, 23141, 6446, 55844, 9839, 151987 |
| GO:0060119 | Inner ear receptor cell development | 4 | 6E-03 | 2562, 2561, 2535, 2558 |
| GO:0007399 | Nervous system development | 29 | 8E-03 | 4208, 5067, 1749, 8128, 2897, 9771, 4762, 2247, 1131, 10178, 1826, 4131, 1641, 1859, 2066, 1136, 6091, 818, 8874, 647135, 340533, 9839, 6334, 2259, 5813, 6327, 6695, 6607, 6606 |
| GO:0060045 | Positive regulation of cardiac muscle cell proliferation | 6 | 9E-03 | 4208, 2066, 2247, 2627, 23414, 983 |
| GO:1901843 | Positive regulation of high voltage-gated calcium channel | 3 | 1E-02 | 2259, 781, 783 |

| | | | | |
|---|---|---|---|---|
| | activity | | | |
| GO:0006479 | Protein methylation | 6 | 1E-02 | 115294, 3275, 2935, 10196, 2107, 56341 |
| GO:0051301 | Cell division | 33 | 1E-02 | 55844, 55743, 57132, 983, 387893, 10296, 6787, 9798, 10776, 7251, 23137, 23244, 219736, 26524, 10592, 23141, 25936, 7465, 55752, 3981, 1021, 91754, 7175, 5500, 5218, 57448, 23047, 905, 23331, 151242, 64282, 998, 901 |
| GO:0006355 | Regulation of transcription, DNA-templated | 112 | 1E-02 | 4208, 147686, 221833, 23512, 1108, 10795, 285349, 153572, 29028, 90987, 7490, 51008, 9753, 2957, 389114, 126017, 7748, 56341, 30813, 677, 204851, 56995, 55193, 4603, 54796, 4601, 406, 253738, 84295, 403341, 728116, 93474, 7799, 80264, 9575, 388561, 144108, 1500, 1993, 57615, 152485, 4090, 10114, 4092, 9425, 55205, 101060200, 27097, 27107, 55206, 6667, 10933, 10499, 152006, 23435, 2186, 283078, 84528, 7594, 51535, 10196, 126068, 25925, 5991, 10957, 57531, 168451, 6929, 7994, 4760, 6660, 9496, 132625, 153222, 1756, 128611, 5411, 5078, 339500, 55892, 96764, 727940, 4848, 57122, 1747, 55769, 90874, 10363, 84458, 5080, 84456, 57786, 55252, 286205, 11176, 5015, 5515, 7767, 367, 256380, 64864, 4904, 9325, 163081, 2140, 57711, 7764, 7761, 2494, 55571, 7559, 4666 |
| GO:0043524 | Negative regulation of neuron apoptotic process | 16 | 1E-02 | 2562, 4208, 2561, 4204, 4035, 3845, 4747, 3981, 3162, 80315, 2558, 5584, 374946, 4915, 10752, 8846 |
| GO:0070848 | Response to growth factor | 5 | 1E-02 | 5028, 4988, 4060, 2627, 1848 |
| GO:0007067 | Mitotic nuclear division | 25 | 1E-02 | 989, 55844, 55743, 5536, 983, 10540, 387893, 55165, 80124, 6787, 10776, 23137, 219736, 26524, 23141, 25936, 7465, 55193, 91754, 7175, 6249, 57448, 23331, 64282, 901 |
| GO:0016573 | Histone acetylation | 7 | 1E-02 | 9329, 10499, 6927, 9425, 4204, 7994, 9575 |
| GO:0021542 | Dentate gyrus development | 5 | 2E-02 | 4208, 1021, 5629, 4760, 11127 |
| GO:0016567 | Protein ubiquitination | 33 | 2E-02 | 115123, 25893, 57534, 8925, 89890, 5080, 57531, 254225, 8835, 26268, 80067, 80124, 7334, 23327, 51257, 26249, 130507, 55832, 11236, 5897, 114088, 56995, 140462, 84541, 152006, 9867, 8452, 7323, 254170, 57448, 23609, 83737, 998 |
| GO:0017157 | Regulation of exocytosis | 6 | 2E-02 | 9699, 9001, 134957, 9515, 203190, 23011 |
| GO:0001570 | Vasculogenesis | 9 | 2E-02 | 3037, 677, 55109, 8322, 93649, 23414, 10052, 4915, 7490 |

| | | | | |
|---|---|---|---|---|
| GO:0050796 | Regulation of insulin secretion | 10 | 2E-02 | 6927, 4082, 3747, 222546, 406, 4760, 775, 80024, 5991, 9575 |
| GO:0034969 | Histone arginine methylation | 3 | 2E-02 | 3275, 10196, 56341 |
| GO:0003310 | Pancreatic A cell differentiation | 3 | 2E-02 | 222546, 3642, 2627 |
| GO:0045022 | Early endosome to late endosome transport | 5 | 2E-02 | 2013, 8411, 64089, 51361, 7879 |
| GO:1903861 | Positive regulation of dendrite extension | 5 | 2E-02 | 9699, 22999, 23327, 57699, 50618 |
| GO:0042787 | Protein ubiquitination involved in ubiquitin-dependent protein catabolic process | 17 | 2E-02 | 26091, 127544, 64750, 154214, 57531, 983, 10296, 55008, 8452, 23327, 339745, 9320, 6921, 26994, 83737, 26249, 64795 |
| GO:0045597 | Positive regulation of cell differentiation | 7 | 2E-02 | 6197, 10911, 5078, 10795, 367, 4760, 3642 |
| GO:0030855 | Epithelial cell differentiation | 10 | 2E-02 | 3189, 9317, 2494, 5230, 2535, 3399, 7416, 1749, 7490, 983 |
| GO:2001237 | Negative regulation of extrinsic apoptotic signaling pathway | 7 | 2E-02 | 57144, 6714, 8837, 367, 29949, 57448, 259230 |
| GO:0014070 | Response to organic cyclic compound | 8 | 3E-02 | 2180, 466, 3747, 4060, 10135, 1848, 983, 54790 |
| GO:0010628 | Positive regulation of gene expression | 25 | 3E-02 | 4208, 1435, 5599, 22999, 3845, 5080, 983, 9699, 367, 5962, 259230, 55737, 5074, 2064, 639, 219736, 6733, 3552, 2488, 5291, 1021, 9314, 4915, 998, 56603 |
| GO:0043433 | Negative regulation of sequence-specific DNA binding transcription factor activity | 9 | 3E-02 | 3162, 1054, 7292, 5629, 6672, 3399, 5727, 4092, 9063 |
| GO:0045732 | Positive regulation of protein catabolic process | 9 | 3E-02 | 5523, 10795, 6642, 23327, 1027, 9554, 7879, 140462, 153222 |
| GO:0045668 | Negative regulation of osteoblast differentiation | 7 | 3E-02 | 1021, 2662, 3399, 4077, 6497, 5727, 79697 |
| GO:0045444 | Fat cell differentiation | 10 | 3E-02 | 26268, 2662, 3164, 59343, 114899, 84230, 9314, 83862, 79689, 79828 |
| GO:0006396 | RNA processing | 12 | 3E-02 | 1996, 51095, 90957, 55109, 3189, 5939, 26747, 23405, 6310, 1736, 3190, 83857 |

| | | | | |
|---|---|---|---|---|
| GO:0030326 | Embryonic limb morphogenesis | 7 | 3E-02 | 6927, 221833, 1749, 9496, 6497, 5727, 56603 |
| GO:0098656 | Anion transmembrane transport | 6 | 3E-02 | 4363, 19, 84230, 23507, 7416, 10060 |
| GO:0007281 | Germ cell development | 6 | 3E-02 | 639, 7073, 1618, 4090, 56853, 7490 |
| GO:0007268 | Chemical synaptic transmission | 23 | 3E-02 | 1138, 9378, 2561, 783, 4040, 2892, 2890, 5376, 3352, 2897, 3357, 6327, 4988, 10911, 3786, 2558, 9001, 5100, 440279, 56479, 10052, 5789, 5594 |
| GO:0001889 | Liver development | 10 | 3E-02 | 64841, 6927, 23705, 9480, 1054, 3720, 5629, 3845, 2627, 23322 |
| GO:0048025 | Negative regulation of mRNA splicing, via spliceosome | 5 | 3E-02 | 1859, 6432, 6434, 56853, 3190 |
| GO:0006469 | Negative regulation of protein kinase activity | 12 | 3E-02 | 8835, 4040, 5573, 1634, 2776, 22843, 7532, 347731, 5611, 60506, 151242, 23768 |
| GO:0018105 | Peptidyl-serine phosphorylation | 14 | 4E-02 | 1859, 166614, 6714, 5599, 5165, 26524, 983, 8569, 2872, 6446, 5584, 10114, 1641, 5594 |
| GO:0030509 | BMP signaling pathway | 10 | 4E-02 | 659, 64750, 2662, 1958, 7323, 4090, 1749, 285704, 6497, 4092 |
| GO:0045892 | Negative regulation of transcription, DNA-templated | 41 | 4E-02 | 7572, 64750, 1602, 147686, 4204, 3720, 7707, 7292, 2627, 5991, 9575, 1027, 7490, 387893, 57786, 2274, 7994, 7767, 144165, 3399, 54816, 7251, 6714, 29966, 3275, 5629, 6310, 9734, 1112, 7764, 79697, 5813, 10472, 23741, 55366, 406, 6672, 9314, 7559, 5451, 23414 |
| GO:0006470 | Protein dephosphorylation | 14 | 4E-02 | 6815, 818, 5537, 5536, 5534, 5523, 5495, 5515, 5500, 85464, 5797, 132160, 5775, 5789 |
| GO:0051145 | Smooth muscle cell differentiation | 4 | 4E-02 | 4208, 57496, 93649, 2627 |
| GO:0072657 | Protein localization to membrane | 4 | 4E-02 | 1363, 7074, 10979, 29993 |
| GO:0060644 | Mammary gland epithelial cell differentiation | 4 | 4E-02 | 2066, 2247, 5727, 2001 |
| GO:0001954 | Positive regulation of cell-matrix adhesion | 5 | 4E-02 | 1756, 1435, 1021, 2013, 27185 |

| | | | | |
|---|---|---|---|---|
| GO:0090316 | Positive regulation of intracellular protein transport | 5 | 4E-02 | 444, 7175, 22902, 10178, 998 |
| GO:0014066 | Regulation of phosphatidylinositol 3-kinase signaling | 10 | 4E-02 | 2066, 5291, 205428, 10818, 2064, 4254, 2549, 26051, 2247, 5594 |
| GO:0007185 | Transmembrane receptor protein tyrosine phosphatase signaling pathway | 3 | 5E-02 | 10818, 7204, 5789 |
| GO:0014033 | Neural crest cell differentiation | 3 | 5E-02 | 4208, 4040, 3357 |
| GO:0051271 | Negative regulation of cellular component movement | 3 | 5E-02 | 6672, 5314, 1027 |
| GO:0032743 | Positive regulation of interleukin-2 production | 4 | 5E-02 | 7292, 5144, 8767, 729230 |
| GO:0007507 | Heart development | 18 | 5E-02 | 4208, 2066, 677, 2064, 341640, 7227, 6091, 775, 9734, 2776, 7490, 27295, 3357, 2549, 80173, 59343, 3399, 5125 |
| **Celullar Component** | | | | |
| GO:0005634 | Nucleus | 385 | 6E-07 | 23512, 6773, 1316, 29028, 7490, 27063, 8738, 11098, 339745, 6497, 10716, 22807, 84083, 6781, 219736, 204851, 2013, 222546, 79718, 8028, 26007, 8725, 406, 5218, 58497, 8863, 84295, 7227, 27185, 8031, 79828, 63901, 388561, 55218, 387893, 7334, 3399, 1845, 6744, 1846, 8428, 55205, 101060200, 340533, 80853, 10743, 2001, 64795, 989, 7594, 10957, 983, 7257, 2247, 3234, 9496, 2107, 128611, 57496, 79755, 96764, 2259, 23741, 6197, 9480, 8452, 2113, 63941, 1602, 7572, 140733, 55156, 865, 10979, 2872, 5203, 5015, 8813, 23774, 9320, 205717, 9325, 3275, 2140, 8609, 7465, 2241, 23764, 466, 2237, 10643, 10644, 10642, 7559, 10785, 51142, 8470, 9338, 132864, 221833, 10795, 9753, 10296, 9931, 54816, 126017, 677, 81620, 56853, 1796, 79697, 83666, 80173, 1618, 29978, 93474, 2627, 7799, 80264, 9575, 9361, 1788, 376132, 9771, 55008, 1500, 2274, 283254, 79026, 23327, 10178, 29777, 6001, 9169, 9635, 6003, 4808, 1641, 9425, 29761, 29966, 388698, 6815, 56970, 23435, 11129, 83737, 2186, 1761, 53335, 126068, 6927, 55818, 6929, 128553, 23405, 9001, 55810, 3190, 132625, 55719, 2066, 1756, 29887, 1859, 2064, 639, 3306, 6733, 339500, 727940, 1854, 1758, 9402, 54960, 3189, 6938, 9960, 3642, 23414, 1747, 9263, 9678, 11176, 132160, 4904, 55832, 6714, 81789, 11164, 1736, 9839, 83640, 3162, 3164, 54014, 6917, 80012, 4208, 147686, 4204, 3720, 7753, 92906, 153572, 55743, 3728, 144165, |

| | | | | |
|---|---|---|---|---|
| | | | | 389114, 84675, 7748, 56341, 23137, 55758, 3096, 3093, 51091, 54790, 4603, 10472, 23261, 7073, 54796, 7175, 342371, 4601, 7074, 1491, 6607, 6606, 253738, 728116, 55727, 3842, 23019, 57615, 152485, 4090, 4092, 151987, 5897, 5629, 10592, 136319, 10499, 57604, 124359, 55783, 54665, 23177, 254065, 80315, 4762, 7994, 4760, 23231, 86, 160851, 55892, 4257, 4848, 9063, 9702, 5500, 8099, 4048, 55769, 10363, 84458, 84456, 11102, 57786, 286205, 5515, 7767, 64864, 259230, 23244, 2551, 9734, 23142, 3981, 7761, 2491, 285498, 2494, 10556, 93649, 4338, 23047, 4666, 1108, 84142, 285349, 90987, 5536, 51008, 10927, 57162, 30813, 22887, 84946, 5939, 84333, 10146, 5451, 26354, 5454, 57050, 64924, 7416, 150280, 26994, 10114, 84515, 26999, 5966, 57144, 26051, 26135, 1112, 8661, 6667, 10933, 5813, 152006, 26747, 6672, 26137, 26136, 283078, 84528, 51503, 7022, 7024, 5991, 25925, 254251, 57531, 5683, 168451, 8569, 1054, 57332, 65110, 25862, 6660, 56478, 5584, 5074, 90957, 5078, 26524, 6310, 25936, 22843, 6446, 375748, 5594, 64750, 5599, 1958, 90874, 1027, 5080, 254225, 57118, 367, 256380, 22823, 6432, 26091, 5862, 163081, 6434, 146713, 25950, 1363, 57711, 137886, 5291, 1021, 91754, 55571, 905 |
| GO:0005737 | Cytoplasm | 366 | 7E-06 | 253943, 4208, 6773, 9860, 159, 56243, 3728, 7490, 27063, 64399, 8738, 22902, 2957, 85464, 339745, 6497, 84675, 84076, 256356, 166614, 6781, 219736, 204851, 100037417, 2013, 6091, 8028, 23507, 3093, 55752, 28511, 50618, 51091, 23261, 9867, 164, 7073, 54796, 7175, 10625, 8725, 4601, 406, 25782, 79092, 5218, 1491, 440279, 8411, 6607, 58497, 6606, 29907, 55654, 399818, 8863, 3845, 23299, 3842, 5611, 63901, 252983, 79828, 55218, 387893, 23019, 7334, 919, 83715, 3399, 4090, 8440, 114294, 729230, 128338, 1848, 6744, 1845, 4092, 151987, 8428, 55109, 51585, 114088, 196483, 10592, 5629, 59338, 30061, 653464, 10499, 27295, 57604, 347731, 121512, 10746, 2001, 64282, 64795, 989, 257397, 10950, 55783, 202018, 51535, 983, 10588, 166336, 23177, 253017, 8835, 4254, 8837, 2247, 80315, 4760, 51271, 154007, 6397, 2107, 160851, 7251, 153222, 51663, 7782, 96764, 4848, 4747, 23741, 6197, 9702, 4976, 55327, 9217, 5314, 9314, 2113, 85439, 56603, 63941, 55156, 1602, 23255, 4048, 23259, 10979, 9098, 23150, 11102, 2872, 1806, 286205, 3357, 5203, 10777, 6787, 10565, 22891, 9320, 10776, 3257, 4131, 9325, 3275, 4139, 2140, 9734, 7763, 154214, 7465, 2488, 2241, 3981, 2491, 375298, 10643, 10644, 2494, 10642, 4338, 348093, 4666, 55344, 10785, 23768, 132864, 10818, 54809, 96459, 1108, 10795, 8925, 5536, 10540, 10296, 26268, 9750, 10135, 23301, 1620, 29993, 130507, 79789, 54623, 677, 115294, 84946, 53942, 4035, 56995, 56853, 1795, 1796, 83666, 26258, 6695, 1618, 134957, 10146, 80031, 29978, 10527, 4189, 55312, 10673, 1124, 1122, 9575, 9900, 9361, 1788, 10015, 55008, 1500, 283254, 79026, 10178, 23327, 5962, 23322, 10114, 6003, 29974, 26999, 57551, 1641, 57144, 55010, 29761, 29966, 54602, 388698, 80352, 6815, 394, 26135, 22866, 2776, 8661, 6667, 5813, 152006, 23435, 5573, 6672, |

| | | | | |
|---|---|---|---|---|
| | | | | 1634, 26136, 83737, 2186, 23331, 124454, 998, 151242, 2182, 23205, 127544, 1761, 64901, 5577, 57534, 8767, 53335, 10196, 3782, 5683, 55582, 8569, 6927, 55818, 6929, 65110, 23405, 50484, 120534, 5584, 56478, 3190, 5775, 132625, 2191, 3037, 5074, 639, 2064, 6733, 26524, 775, 6310, 60682, 372, 9397, 1758, 9402, 54960, 88455, 6938, 375743, 6249, 222658, 84230, 9960, 6446, 23414, 100127983, 55705, 10107, 5594, 64750, 54439, 1958, 25941, 761, 6240, 64759, 1749, 9847, 1027, 663, 361, 5080, 60506, 57118, 11176, 80124, 367, 3658, 6642, 5797, 4904, 55832, 22823, 659, 26091, 6432, 6714, 145482, 25950, 146713, 1736, 1021, 3164, 54014, 91754, 2535, 4919, 901 |
| GO:0030054 | Cell junction | 49 | 5E-05 | 375567, 59283, 2897, 23177, 3775, 9699, 80315, 22902, 10135, 1826, 9001, 29993, 203190, 23137, 1138, 54623, 1136, 100506658, 2892, 2890, 55752, 9867, 134957, 440279, 85439, 23312, 83706, 2562, 2561, 3747, 22999, 27185, 57706, 386617, 3357, 1131, 3222, 2558, 9635, 4131, 26999, 9378, 114088, 9899, 2241, 388662, 26136, 5100, 347731 |
| GO:0005654 | Nucleoplasm | 205 | 1E-04 | 4208, 3720, 23512, 7753, 6773, 29028, 7490, 27063, 9866, 256471, 59343, 2957, 6497, 23137, 204851, 219736, 79718, 8028, 10472, 51095, 7073, 7175, 342371, 10625, 406, 6603, 10622, 6607, 6606, 84295, 403341, 7227, 252983, 3840, 387893, 7334, 1993, 4090, 3399, 1848, 4092, 1845, 151987, 1846, 8428, 51585, 5897, 818, 10592, 59338, 80853, 10743, 10499, 7323, 10746, 257397, 51535, 92856, 983, 7994, 154007, 4760, 23231, 86, 153222, 5411, 51663, 2892, 96764, 9063, 4973, 6197, 23741, 9702, 9480, 4976, 8452, 5500, 9314, 2113, 85437, 4048, 10979, 5711, 11102, 55252, 2872, 286205, 10565, 23774, 9320, 10776, 9325, 9329, 23244, 2551, 3275, 2140, 9734, 7465, 7764, 3981, 2491, 23764, 2237, 466, 2494, 10556, 7559, 23047, 10785, 9338, 84142, 1108, 5536, 51008, 5534, 10135, 54623, 81620, 84946, 54629, 79697, 55193, 26258, 83666, 5451, 26354, 10527, 57050, 1122, 2627, 9575, 9361, 1788, 2274, 3222, 23327, 9169, 29974, 84515, 5966, 55010, 9425, 29966, 5567, 27107, 6667, 10933, 152006, 23435, 6672, 26137, 2186, 51503, 7022, 7707, 53335, 11193, 5683, 6927, 8569, 6929, 55818, 84248, 1054, 57332, 65110, 50484, 3190, 6921, 2066, 1859, 639, 6733, 5078, 6310, 1854, 57122, 9402, 3189, 3642, 6446, 23414, 5594, 64750, 5599, 1958, 6240, 1027, 5080, 254225, 11176, 367, 4904, 55832, 22823, 6432, 6434, 1736, 1021, 3164, 6917, 80012, 905 |
| GO:0005768 | Endosome | 27 | 6E-04 | 10818, 10890, 115123, 8417, 1027, 79689, 60592, 23011, 3775, 5869, 29993, 5584, 160851, 7251, 4074, 55737, 7037, 4035, 9402, 26258, 9182, 6249, 57561, 57448, 374868, 4642, 4915 |
| GO:0008021 | Synaptic vesicle | 15 | 8E-04 | 114088, 27185, 30061, 9899, 2890, 7416, 9900, 388662, 51305, 9001, 134957, 85439, 23312, 203190, 83992 |
| GO:0005667 | Transcription factor complex | 24 | 8E-04 | 1602, 7227, 55758, 2140, 9734, 2627, 5991, 9575, 27063, 51008, 6927, 6938, 6929, 3164, 10625, 406, 4090, 2113, 29777, 6497, 4092, 4808, 5966, 5454 |

| | | | | |
|---|---|---|---|---|
| GO:0030425 | Dendrite | 34 | 2E-03 | 3747, 92737, 57706, 2785, 3357, 3775, 22902, 80315, 1131, 23405, 23774, 2558, 729230, 1620, 10752, 659, 29966, 3572, 1136, 114088, 4035, 2776, 2890, 54407, 5028, 3352, 5813, 4976, 8725, 342371, 5100, 26047, 2186, 9312 |
| GO:0043204 | Perikaryon | 15 | 3E-03 | 3747, 146713, 57699, 5536, 4988, 3775, 4762, 22902, 23774, 729230, 26047, 6607, 6606, 9312, 5594 |
| GO:0005829 | Cytosol | 225 | 5E-03 | 55233, 4208, 56344, 4204, 5230, 6773, 55844, 159, 59345, 3728, 27063, 8738, 25791, 144165, 56341, 2013, 8874, 51361, 26007, 7879, 51091, 164, 10096, 7074, 10622, 5218, 1491, 8411, 6607, 7204, 6606, 29907, 283209, 3845, 3842, 5611, 3840, 23019, 7334, 8440, 4090, 55526, 729230, 1848, 4092, 1845, 55737, 8428, 818, 10592, 136319, 9182, 7323, 64087, 5238, 121512, 64089, 526, 10746, 989, 257397, 55276, 7456, 983, 64841, 10588, 7447, 8835, 8837, 7257, 51271, 4077, 2107, 10869, 9061, 96764, 4848, 4747, 6197, 9702, 8452, 5500, 586, 4642, 2561, 4048, 23259, 10979, 5711, 60592, 1806, 5203, 5515, 140838, 10565, 9320, 26960, 4131, 9325, 23244, 3275, 9732, 154214, 2241, 2491, 10643, 2549, 10644, 10642, 4338, 7095, 23047, 10785, 192668, 81631, 5537, 8925, 5536, 10540, 5534, 10135, 29993, 57162, 677, 81620, 84946, 3552, 1796, 90427, 83666, 26258, 10146, 10527, 55312, 5144, 22999, 1124, 1122, 10015, 55008, 1500, 3418, 79026, 23327, 6001, 5137, 23322, 6003, 26999, 10802, 1641, 5966, 57551, 6173, 6815, 394, 5567, 8661, 8667, 23433, 5573, 1121, 11127, 7532, 83737, 998, 23205, 127544, 22931, 5577, 8767, 57534, 10196, 6517, 57132, 5683, 8569, 9698, 65110, 23405, 9798, 5584, 56478, 6921, 1756, 2066, 3306, 257106, 26524, 25936, 372, 57122, 9397, 9402, 54960, 51479, 5495, 6249, 6446, 55705, 5594, 262, 64750, 5599, 761, 2935, 6240, 663, 1027, 367, 3658, 6642, 26249, 26091, 6134, 6714, 11164, 137886, 5291, 1021, 3162, 91754, 55571, 3067, 4915 |
| GO:0014069 | Postsynaptic density | 21 | 5E-03 | 659, 55737, 6714, 1136, 775, 27185, 818, 57534, 22866, 2890, 59283, 5727, 2785, 5028, 8825, 9867, 55327, 6695, 80315, 4915, 4131 |
| GO:0005911 | Cell-cell junction | 20 | 5E-03 | 659, 10256, 10818, 23705, 10890, 5411, 100506658, 6523, 9900, 3728, 8825, 8322, 55327, 1500, 10096, 7074, 5797, 23322, 998, 23768 |
| GO:0045211 | Postsynaptic membrane | 23 | 5E-03 | 1138, 1756, 2562, 2561, 3747, 1136, 27185, 2892, 22866, 2890, 59283, 386617, 5028, 2897, 8825, 9867, 55327, 80315, 1131, 2558, 5100, 347731, 4915 |
| GO:0043005 | Neuron projection | 25 | 5E-03 | 132864, 57699, 9900, 79608, 4988, 80315, 10178, 203190, 26999, 1641, 6714, 23705, 8874, 9899, 22866, 2693, 8825, 493, 55327, 85439, 4642, 6607, 6606, 998, 23768 |
| GO:0005901 | Caveola | 10 | 1E-02 | 659, 6714, 3162, 493, 4040, 3778, 64805, 2013, 5727, 5594 |
| GO:0043025 | Neuronal cell body | 29 | 2E-02 | 4040, 5577, 92737, 6240, 6323, 3782, 57706, 79608, 729230, 1620, 6001, 659, 9378, 29966, 3572, 1136, 1363, 4035, 8874, 22866, 2890, 54407, 5813, 7074, 11127, 26047, 4642, 998, 2182 |

| | | | | |
|---|---|---|---|---|
| GO:0005891 | Voltage-gated calcium channel complex | 6 | 2E-02 | 781, 783, 10345, 5144, 775, 59283 |
| GO:0016605 | PML body | 12 | 3E-02 | 2872, 204851, 59343, 7994, 406, 6672, 56478, 55743, 6497, 10114, 23137, 9063 |
| GO:0000790 | Nuclear chromatin | 19 | 3E-02 | 5074, 2551, 7227, 1108, 1749, 5991, 6667, 2241, 5080, 6938, 6929, 57332, 367, 6603, 9960, 10622, 9314, 3190, 86 |
| GO:0016323 | Basolateral plasma membrane | 18 | 3E-02 | 2066, 2064, 23705, 26504, 3747, 4363, 7037, 30061, 361, 84466, 5028, 8825, 3953, 493, 55327, 1131, 6563, 4642 |
| GO:0045202 | Synapse | 18 | 3E-02 | 1756, 2562, 132864, 2561, 23705, 4040, 3747, 375567, 3357, 3775, 80315, 1131, 2558, 8440, 1826, 29993, 23768, 26999 |
| GO:0030904 | Retromer complex | 5 | 4E-02 | 55737, 51479, 6642, 7879, 4074 |
| GO:0000139 | Golgi membrane | 46 | 4E-02 | 10678, 1571, 84912, 10890, 64131, 5537, 60592, 79689, 8459, 57531, 4247, 51026, 9554, 8128, 84002, 10396, 64841, 79608, 10565, 2526, 5584, 9953, 205717, 10802, 259230, 5862, 54602, 55032, 2013, 2890, 372, 283464, 2591, 340481, 205428, 9867, 164, 54947, 8452, 9217, 9331, 6482, 56900, 51311, 998, 8706 |
| GO:0030018 | Z disc | 13 | 4E-02 | 1756, 27295, 493, 91624, 10529, 2274, 775, 6323, 6607, 3728, 6606, 8470, 6334 |
| GO:1990454 | L-type voltage-gated calcium channel complex | 3 | 4E-02 | 781, 783, 775 |
| GO:1904115 | Axon cytoplasm | 6 | 5E-02 | 26258, 4976, 9001, 11127, 23011, 4747 |
| **Molecular Function** | | | | |
| GO:0005515 | Protein binding | 593 | 4E-07 | 55233, 10890, 23512, 5230, 6773, 8417, 27069, 1316, 159, 7490, 27063, 5376, 8738, 8322, 22902, 59343, 8325, 6354, 2957, 1826, 339745, 6497, 22807, 84083, 204851, 219736, 2013, 222546, 6091, 8874, 100506658, 79718, 23507, 8028, 51361, 26007, 50618, 140462, 164, 59350, 8725, 10096, 25780, 55366, 406, 5218, 8411, 58497, 7204, 29907, 8863, 84295, 1435, 7227, 27185, 22918, 8100, 63901, 55218, 387893, 7334, 919, 8440, 3399, 8853, 8428, 55109, 818, 196483, 59338, 27097, 10743, 27295, 118429, 22915, 6383, 63908, 7323, 526, 10746, 374986, 64795, 7220, 989, 10950, 225689, 84171, 55276, 23554, 10957, 7456, 983, 8050, 7447, 8835, 8837, 7257, 2247, 10651, 51271, 6397, 2107, 7251, 131034, 10869, 57496, 79755, 2890, 96764, 2259, 444, 8825, 6197, 23741, 4973, 7381, 4976, 2882, 8452, 9217, 5314, 9313, 2113, 9314, |

| | | | | | |
|---|---|---|---|---|---|
| | | | | | 85439, 26047, 5125, 63941, 55156, 7572, 1602, 387338, 865, 257194, 10979, 60592, 2872, 1806, 5203, 4988, 5015, 8813, 9320, 51257, 3257, 9325, 9329, 3275, 57470, 2140, 953, 7465, 2241, 55349, 84063, 10643, 2237, 466, 10644, 8803, 10642, 84970, 348093, 10785, 374946, 8470, 51142, 81631, 54809, 192668, 4040, 96459, 159195, 3684, 9554, 9749, 10298, 9753, 9750, 1620, 29993, 79583, 79596, 54623, 677, 23705, 81620, 4035, 3552, 1795, 1796, 79697, 90427, 83666, 11006, 1618, 166824, 81618, 29978, 4189, 10673, 83899, 2627, 93474, 9575, 9361, 1788, 2274, 1500, 79026, 23327, 29777, 9169, 23322, 29974, 4808, 1641, 9378, 6173, 55010, 399947, 9425, 29761, 29966, 54602, 6815, 56970, 2776, 9758, 283461, 9767, 23435, 158297, 11127, 1634, 2186, 83737, 998, 783, 23200, 10196, 11193, 10396, 9695, 6927, 3778, 6929, 9699, 128553, 9698, 23405, 3632, 9798, 120534, 9001, 3190, 55810, 6921, 2191, 55719, 1859, 29887, 1756, 2066, 2064, 639, 6733, 3306, 775, 80129, 9818, 1854, 9402, 131566, 54960, 3189, 6938, 6249, 23608, 23609, 23414, 55705, 2935, 6240, 663, 54838, 11176, 3658, 222663, 4904, 55832, 659, 6714, 6134, 3572, 11164, 83786, 1736, 9839, 83640, 3162, 3164, 6917, 80012, 2535, 4919, 3067, 4208, 253943, 147686, 10345, 56344, 4204, 7753, 92906, 4103, 55743, 3728, 64399, 144165, 84675, 348110, 23137, 56341, 55758, 3096, 3093, 55752, 7879, 54790, 51091, 10472, 342371, 7175, 4601, 7074, 6603, 1491, 6607, 6606, 19, 3845, 23299, 3842, 23011, 252983, 3840, 23019, 55668, 1993, 4090, 4092, 151987, 55737, 3730, 114088, 10592, 5629, 30061, 9899, 10499, 9182, 64087, 7058, 124359, 4148, 257397, 51535, 4753, 89890, 92856, 23177, 4253, 4254, 4762, 80315, 7994, 4760, 4153, 4077, 86, 153222, 4360, 11236, 51663, 64062, 4848, 4257, 4747, 9063, 3953, 9702, 5500, 8099, 4048, 4879, 92737, 220965, 5711, 10363, 55769, 9098, 84458, 84456, 286205, 5515, 7767, 10565, 22891, 4131, 23244, 10256, 2551, 23141, 9734, 7763, 7764, 9732, 23142, 2488, 5727, 51642, 3981, 5523, 9728, 2491, 2549, 2494, 4060, 10556, 3977, 93649, 254170, 10558, 23047, 10457, 10818, 10130, 5165, 1108, 84142, 5537, 5536, 51008, 10540, 5534, 26268, 10927, 6565, 10135, 57162, 84946, 4318, 55193, 5028, 26258, 57561, 84333, 54329, 92703, 5451, 10146, 26354, 5454, 10527, 10424, 84912, 10529, 5144, 22999, 1124, 57050, 6400, 7416, 2785, 10015, 55165, 1131, 91833, 26994, 5962, 151254, 84515, 5828, 26999, 57551, 10802, 5966, 57144, 84923, 23592, 26051, 134553, 26135, 394, 5567, 1112, 8661, 10418, 6667, 8667, 151647, 54407, 5813, 10933, 493, 10026, 5573, 6672, 26747, 26136, 7532, 51503, 84528, 5955, 7022, 127544, 7024, 91526, 22931, 7707, 5577, 57534, 8767, 6517, 5991, 254251, 57531, 57132, 5683, 124401, 8569, 84248, 1054, 57332, 25862, 65110, 50484, 6660, 56478, 50486, 5584, 56479, 5775, 219333, 1138, 5074, 90957, 6646, 26524, 7037, 6310, 25936, 7504, 22843, 57122, 51479, 5495, 8785, 84230, 6446, 56681, 10107, 5789, 5594, 64750, 5599, 1958, 64759, 1027, 5080, 91404, 57515, 367, 5869, 6642, 5797, 26249, 6432, 5862, 6434, 163081, |

| | | | | |
|---|---|---|---|---|
| | | | | 25950, 6523, 137886, 6334, 5291, 1021, 57105, 91754, 55571, 57448, 905 |
| GO:0003730 | mRNA 3'-UTR binding | 13 | 2E-05 | 677, 132864, 26135, 1996, 9728, 10643, 23435, 80315, 9698, 10644, 1993, 10642, 1618 |
| GO:0008270 | Zinc ion binding | 102 | 5E-05 | 1108, 55743, 170692, 7490, 9866, 64399, 144165, 6497, 84675, 130507, 7748, 79596, 84083, 4312, 4318, 8028, 54790, 9867, 54796, 84333, 8411, 84295, 10529, 7227, 22998, 64924, 7799, 2627, 2274, 26994, 9169, 5828, 10802, 80352, 5897, 114088, 56970, 80853, 10743, 27295, 152006, 9767, 6672, 26136, 2186, 124454, 127544, 257397, 57534, 25893, 11193, 166336, 7994, 25862, 4077, 79752, 1756, 11236, 51663, 55892, 9063, 6249, 9313, 23608, 9960, 9314, 23414, 23609, 85437, 10107, 7572, 9317, 4048, 1958, 115123, 761, 11174, 254225, 57786, 9678, 11176, 367, 23774, 51257, 4325, 1368, 9325, 22823, 6432, 145482, 1363, 8609, 7763, 7761, 340481, 285498, 3164, 2494, 6917, 80012, 91612, 7092 |
| GO:0003700 | Transcription factor activity, sequence-specific DNA binding | 85 | 1E-04 | 4208, 7022, 147686, 7024, 4204, 1761, 7594, 7753, 6773, 10795, 285349, 5991, 90987, 7490, 9753, 6927, 6929, 55818, 4762, 4760, 3234, 6660, 9496, 10716, 22807, 132625, 153222, 677, 30813, 639, 128611, 22887, 55758, 56995, 8028, 55892, 10472, 6938, 406, 9314, 2113, 3642, 5451, 5454, 7572, 1958, 865, 7227, 1747, 55769, 2627, 7799, 10363, 9575, 5080, 388561, 5015, 7767, 367, 3222, 3399, 4090, 64864, 4092, 4904, 5966, 2551, 55205, 163081, 29966, 5629, 8609, 1112, 27097, 7764, 27107, 6667, 5813, 10743, 23764, 466, 23435, 2494, 7559, 9338 |
| GO:0001077 | Transcriptional activator activity, RNA polymerase II core promoter proximal region sequence-specific binding | 27 | 2E-03 | 4208, 7022, 1958, 1747, 1749, 5080, 7490, 6927, 5015, 1054, 367, 4760, 55810, 3190, 4904, 2551, 79755, 4603, 23261, 9480, 6938, 3164, 2494, 93649, 9314, 253738, 5454 |
| GO:0044212 | Transcription regulatory region DNA binding | 25 | 2E-03 | 2066, 4208, 30813, 2551, 55205, 128611, 163081, 222546, 55758, 3096, 5629, 79718, 1749, 2627, 7764, 5991, 6667, 7490, 6927, 6938, 2494, 367, 9314, 22807, 4092 |
| GO:0046872 | Metal ion binding | 153 | 2E-03 | 55233, 5169, 147686, 221833, 23512, 7753, 10795, 285349, 1316, 5537, 79689, 90987, 5536, 7490, 114826, 3684, 9753, 9931, 389114, 54816, 11019, 22807, 126017, 84076, 7748, 677, 3096, 100528021, 219539, 26007, 10472, 59350, 54796, 440279, 8411, 58497, 253738, 403341, 728116, 7227, 55312, 283209, 5144, 22999, 5145, 1124, 3845, 7799, 93474, 80264, 388561, 1788, 57615, 283254, 4090, 152485, 9635, 5137, 4092, 8853, 9378, 8428, 55205, 101060200, 2776, 27107, 10418, 6667, 54482, 493, 118429, 26747, 26137, 5238, 121512, 10746, 64282, 781, 388403, 1761, 7594, 7707, 53335, 10196, 126068, 25925, 10588, 168451, 3778, 9699, 128553, 7257, 23405, 154007, 50484, 5584, 160851, 132625, 90957, 639, 128611, 10869, 341640, 26524, 775, 79755, 339500, 60682, 22843, 64062, 55892, 51479, 6249, 5500, 23608, 3642, 9314, 23414, |

| | | | | |
|---|---|---|---|---|
| | | | | 64172, 8515, 7572, 55156, 54439, 1958, 23259, 64759, 83591, 55769, 90874, 55252, 2872, 1806, 5515, 7767, 3658, 659, 163081, 2140, 57711, 8609, 9734, 51646, 154214, 7764, 9839, 3981, 7761, 3162, 91754, 8803, 9331, 7559, 8470 |
| GO:0003677 | DNA binding | 127 | 2E-03 | 4208, 221833, 4204, 7753, 1108, 6773, 10795, 1316, 90987, 27063, 2957, 389114, 85464, 54816, 10716, 126017, 30813, 677, 81620, 204851, 55758, 3096, 8028, 55193, 54790, 4603, 84542, 10472, 54796, 342371, 4601, 406, 10146, 253738, 84295, 403341, 728116, 2627, 93474, 7799, 9575, 388561, 1788, 144108, 152485, 29777, 26994, 84515, 55010, 23592, 101060200, 5897, 5629, 6667, 26747, 6672, 26137, 64282, 51311, 7022, 7024, 7594, 126068, 25925, 5991, 254251, 168451, 6927, 6929, 128553, 1054, 7257, 4762, 7994, 6660, 9496, 3190, 132625, 7251, 5078, 51663, 5411, 339500, 6310, 55892, 9063, 2113, 375748, 23414, 5594, 1602, 7572, 1958, 865, 83591, 55769, 90874, 10363, 84458, 5080, 55252, 11176, 5015, 7767, 367, 64864, 205717, 4904, 22823, 9329, 2551, 146713, 81789, 57711, 8609, 7763, 9839, 3981, 7761, 2237, 3164, 2494, 6917, 80012, 7559, 23047, 4666 |
| GO:0016874 | Ligase activity | 29 | 3E-03 | 64750, 127544, 115123, 25893, 57534, 55743, 8925, 57531, 254225, 23327, 9320, 51257, 130507, 57162, 26091, 11236, 114088, 154214, 9063, 3981, 2180, 152006, 9867, 285498, 8803, 23608, 64087, 57448, 23609 |
| GO:0043565 | Sequence-specific DNA binding | 47 | 3E-03 | 7572, 7024, 1958, 7227, 7707, 27185, 1747, 153572, 9575, 5080, 9361, 7490, 9753, 7257, 5015, 1054, 367, 4760, 3222, 3234, 55810, 153222, 30813, 2551, 55205, 128611, 22887, 5078, 3096, 1112, 727940, 6667, 136319, 10472, 23261, 23764, 466, 3164, 2494, 406, 2113, 5451, 283078, 2186, 84528, 51142, 2001 |
| GO:0004842 | Ubiquitin-protein transferase activity | 33 | 4E-03 | 64750, 127544, 57534, 8925, 55743, 89890, 5080, 57531, 10296, 26268, 55008, 7334, 23327, 9320, 6921, 51257, 26994, 130507, 26249, 26091, 11236, 114088, 154214, 3093, 84541, 152006, 9867, 8452, 7323, 23608, 57448, 83737, 64795 |
| GO:0035035 | Histone acetyltransferase binding | 7 | 5E-03 | 23741, 1958, 93649, 2113, 6667, 5080, 9325 |
| GO:0030332 | Cyclin binding | 6 | 5E-03 | 1021, 8452, 5218, 3642, 5727, 983 |
| GO:0003723 | RNA binding | 48 | 6E-03 | 253943, 23512, 92906, 24138, 7490, 5536, 51008, 11102, 1996, 11176, 9698, 1993, 65110, 3658, 29777, 3190, 2107, 29974, 4904, 55109, 57470, 146713, 6310, 1736, 8661, 64062, 25973, 5813, 83640, 51095, 11022, 3189, 7073, 10643, 342371, 23435, 10556, 1618, 5939, 26747, 10642, 6607, 85437, 124454, 6606, 51503, 51311, 166863 |
| GO:0000978 | RNA polymerase II core promoter proximal region sequence-specific DNA binding | 34 | 6E-03 | 4208, 7707, 1108, 53335, 1749, 5991, 9575, 80264, 5080, 6929, 5015, 367, 57615, 4760, 6497, 64864, 55810, 3190, 86, 4904, 7748, 2551, 639, 222546, 7764, 6667, 4603, 10499, 9480, 6938, 2494, 6603, 3642, 5454 |

| | | | | |
|---|---|---|---|---|
| GO:0003729 | mRNA binding | 16 | 7E-03 | 677, 6434, 4204, 51585, 146713, 92906, 56853, 8661, 1996, 84248, 7175, 10644, 10642, 56478, 348093, 10146 |
| GO:0000900 | Translation repressor activity, nucleic acid binding | 4 | 2E-02 | 5813, 132864, 80315, 56853 |
| GO:0071837 | HMG box domain binding | 5 | 2E-02 | 4208, 6938, 1749, 6667, 5080 |
| GO:0086007 | Voltage-gated calcium channel activity involved in cardiac muscle cell action potential | 3 | 2E-02 | 781, 783, 775 |
| GO:0001105 | RNA polymerase II transcription coactivator activity | 7 | 2E-02 | 57496, 4760, 93649, 23414, 27063, 116931, 5454 |
| GO:0019904 | Protein domain specific binding | 21 | 2E-02 | 4204, 5577, 100506658, 25925, 57132, 5534, 4747, 23177, 4988, 23019, 55327, 59343, 6672, 23405, 9798, 3399, 7532, 5584, 5962, 6497, 4642 |
| GO:0019888 | Protein phosphatase regulator activity | 6 | 3E-02 | 5523, 26051, 23141, 55844, 10776, 151987 |
| GO:0017137 | Rab GTPase binding | 15 | 3E-02 | 22999, 1122, 23299, 57706, 57531, 51479, 164, 9699, 1121, 25782, 134957, 11127, 9515, 23312, 55654 |
| GO:0016301 | Kinase activity | 23 | 3E-02 | 10256, 6714, 2064, 5599, 4139, 5577, 8767, 22866, 5567, 4753, 7465, 1027, 1796, 548596, 6197, 5291, 375298, 5573, 29993, 4915, 5594, 57551, 259230 |
| GO:0001078 | Transcriptional repressor activity, RNA polymerase II core promoter proximal region sequence-specific binding | 13 | 3E-02 | 1602, 7022, 639, 7707, 5629, 53335, 7764, 6929, 128553, 26137, 3642, 6497, 23414 |
| GO:0010385 | Double-stranded methylated DNA binding | 3 | 3E-02 | 4204, 1958, 7490 |
| GO:0016868 | Intramolecular transferase activity, phosphotransferases | 3 | 3E-02 | 283209, 5238, 55276 |
| GO:0004222 | Metalloendopeptidase activity | 13 | 3E-02 | 64172, 4312, 128553, 4318, 9986, 9313, 11174, 9635, 4325, 170692, 7092, 2191, 79875 |
| GO:0035064 | Methylated histone binding | 8 | 3E-02 | 22823, 9425, 10927, 57332, 23512, 5897, 80853, 124359 |
| GO:0003713 | Transcription coactivator activity | 23 | 4E-02 | 2551, 3275, 57496, 865, 56970, 8031, 8609, 27097, 3728, 9063, 10499, 6929, 2274, 7994, 4760, 93649, 6672, 2957, 6603, 4666, 29777, 86, 9325 |

| | | | | |
|---|---|---|---|---|
| GO:0042813 | WNT-activated receptor activity | 5 | 4E-02 | 4040, 8322, 8325, 2535, 23554 |
| GO:0004683 | Calmodulin-dependent protein kinase activity | 5 | 4E-02 | 8569, 2872, 57118, 818, 1641 |
| GO:0001046 | Core promoter sequence-specific DNA binding | 7 | 4E-02 | 4208, 79755, 5629, 93649, 9960, 9575, 6667 |
| GO:0045182 | Translation regulator activity | 3 | 4E-02 | 10643, 10644, 10642 |
| GO:0003747 | Translation release factor activity | 3 | 4E-02 | 2935, 91574, 2107 |
| GO:0005085 | Guanyl-nucleotide exchange factor activity | 13 | 4E-02 | 9771, 64805, 96459, 25780, 7074, 25791, 8874, 10565, 9732, 121512, 1795, 3257, 7204 |
| GO:0008134 | Transcription factor binding | 25 | 5E-02 | 4204, 55758, 9734, 2627, 6667, 5080, 9063, 5813, 10499, 6938, 6929, 1054, 2274, 7994, 367, 93649, 6672, 4760, 2957, 3399, 2113, 23414, 2186, 51142, 5594 |
| GO:0070888 | E-box binding | 6 | 5E-02 | 6938, 6929, 4762, 406, 4760, 9575 |
| GO:0030742 | GTP-dependent protein binding | 5 | 5E-02 | 9771, 164, 5869, 8411, 998 |

| KEGG ID | KEGG Term | Count | PValue | Genes (Entrez ID) |
|---|---|---|---|---|
| hsa04550 | Signaling pathways regulating pluripotency of stem cells | 22 | 4E-05 | 659, 3572, 3720, 3845, 1749, 5080, 3626, 6927, 5291, 6929, 8322, 2247, 4762, 7994, 8325, 3977, 4090, 3399, 2535, 84333, 9314, 5594 |
| hsa04010 | MAPK signaling pathway | 28 | 1E-03 | 781, 4208, 783, 5599, 3845, 59283, 5536, 5534, 2872, 8569, 2247, 1848, 1845, 57551, 1846, 3306, 775, 5567, 3552, 2259, 6197, 5495, 3164, 25780, 4915, 10746, 998, 5594 |
| hsa05205 | Proteoglycans in cancer | 23 | 2E-03 | 2066, 10818, 2064, 6714, 4318, 818, 3845, 5567, 5727, 5291, 8322, 2549, 2247, 6383, 4060, 8325, 7074, 5500, 2535, 1634, 5962, 998, 5594 |
| hsa04012 | ErbB signaling pathway | 13 | 4E-03 | 57144, 10298, 2066, 5291, 6714, 2064, 5599, 2549, 818, 8440, 3845, 1027, 5594 |
| hsa04728 | Dopaminergic synapse | 16 | 6E-03 | 5599, 818, 775, 6323, 2892, 55844, 2776, 5567, 2890, 59345, 9575, 2785, 5523, 5515, 5500, 406 |

| | | | | |
|---|---|---|---|---|
| hsa04723 | Retrograde endocannabinoid signaling | 13 | 1E-02 | 2562, 2561, 5599, 775, 22999, 2892, 2776, 5567, 2890, 59345, 2785, 2558, 5594 |
| hsa04720 | Long-term potentiation | 10 | 1E-02 | 6197, 818, 775, 5500, 5567, 3845, 2776, 2890, 5594, 5534 |
| hsa04912 | GnRH signaling pathway | 12 | 1E-02 | 6714, 5599, 818, 775, 5567, 3845, 2776, 2798, 2488, 10746, 998, 5594 |
| hsa04950 | Maturity onset diabetes of the young | 6 | 1E-02 | 6927, 5078, 222546, 2494, 4760, 5080 |
| hsa04725 | Cholinergic synapse | 13 | 2E-02 | 1136, 818, 775, 2776, 3845, 5567, 59345, 2785, 5291, 1131, 3786, 56479, 5594 |
| hsa04910 | Insulin signaling pathway | 15 | 3E-02 | 5599, 5577, 3845, 5567, 6517, 2872, 5291, 8569, 8835, 23433, 5573, 2538, 5500, 5584, 5594 |
| hsa05211 | Renal cell carcinoma | 9 | 3E-02 | 10298, 57144, 5291, 2549, 3845, 2113, 6921, 998, 5594 |
| hsa04014 | Ras signaling pathway | 21 | 3E-02 | 57144, 1435, 5599, 3845, 5567, 59345, 2785, 9771, 10298, 2259, 5291, 4254, 2549, 2247, 25780, 7074, 5869, 2113, 998, 5966, 5594 |
| hsa04144 | Endocytosis | 22 | 3E-02 | 55737, 8853, 64750, 6714, 3306, 10890, 7037, 60682, 7879, 7456, 57132, 10015, 10565, 9798, 5869, 6642, 23327, 8411, 5584, 83737, 998, 7251 |
| hsa03013 | RNA transport | 17 | 4E-02 | 5411, 9818, 8661, 96764, 8667, 57122, 11102, 54960, 51095, 7175, 59343, 65110, 10556, 285190, 6607, 6606, 26999 |
| hsa04350 | TGF-beta signaling pathway | 10 | 5E-02 | 659, 64750, 5515, 4090, 3399, 1634, 6667, 3626, 4092, 5594 |
